# Single-strand mismatch and damage patterns revealed by single-molecule DNA sequencing

**DOI:** 10.1101/2023.02.19.526140

**Authors:** Mei Hong Liu, Benjamin Costa, Una Choi, Rachel C. Bandler, Emilie Lassen, Marta Grońska-Pęski, Adam Schwing, Zachary R. Murphy, Daniel Rosenkjær, Shany Picciotto, Vanessa Bianchi, Lucie Stengs, Melissa Edwards, Caitlin A. Loh, Tina K. Truong, Randall E. Brand, Tomi Pastinen, J. Richard Wagner, Anne-Bine Skytte, Uri Tabori, Jonathan E. Shoag, Gilad D. Evrony

**Affiliations:** Center for Human Genetics and Genomics, New York University Grossman School of Medicine, USA; Department of Pediatrics, Department of Neuroscience & Physiology, Institute for Systems Genetics, Perlmutter Cancer Center, and Neuroscience Institute, New York University Grossman School of Medicine, USA; Cryos International Sperm and Egg Bank, Denmark; Department of Urology, University Hospitals Cleveland Medical Center, Case Western Reserve University School of Medicine, USA; Program in Genetics and Genome Biology, Peter Gilgan Centre for Research and Learning, The Hospital for Sick Children, Canada; Department of Medicine, University of Pittsburgh School of Medicine, USA; Genomic Medicine Center, Children’s Mercy Kansas City, USA; Department of Nuclear Medicine and Radiobiology, Université de Sherbrooke, Canada; Division of Haematology/Oncology, Arthur and Sonia Labatt Brain Tumour Research Centre, The Hospital for Sick Children, Canada

## Abstract

Mutations accumulate in the genome of every cell of the body throughout life, causing cancer and other genetic diseases^1-4^. Almost all of these mosaic mutations begin as nucleotide mismatches or damage in only one of the two strands of the DNA prior to becoming double-strand mutations if unrepaired or misrepaired^5^. However, current DNA sequencing technologies cannot resolve these initial single-strand events. Here, we developed a single-molecule, long-read sequencing method that achieves single-molecule fidelity for single-base substitutions when present in either one or both strands of the DNA. It also detects single-strand cytosine deamination events, a common type of DNA damage. We profiled 110 samples from diverse tissues, including from individuals with cancer-predisposition syndromes, and define the first single-strand mismatch and damage signatures. We find correspondences between these single-strand signatures and known double-strand mutational signatures, which resolves the identity of the initiating lesions. Tumors deficient in both mismatch repair and replicative polymerase proofreading show distinct single-strand mismatch patterns compared to samples deficient in only polymerase proofreading. In the mitochondrial genome, our findings support a mutagenic mechanism occurring primarily during replication. Since the double-strand DNA mutations interrogated by prior studies are only the endpoint of the mutation process, our approach to detect the initiating single-strand events at single-molecule resolution will enable new studies of how mutations arise in a variety of contexts, especially in cancer and aging.

## Main

Mosaic mutations are ubiquitous in the body and accumulate throughout life in every cell^3,6^. Most mosaic mutations begin as nucleotide mismatches or damage in only one of the two strands of the DNA double helix^5,7,8^. When these single-strand DNA (ssDNA) events are misrepaired, or when they are replicated during the cell cycle prior to repair, they then become permanent double-strand DNA (dsDNA) mosaic mutations^5^. However, these ssDNA events, which are the origin of most mutations in the body, have remained invisible to current DNA profiling methods, which only reliably detect dsDNA mutations. This is because all current methods for profiling mosaic mutations—single-cell genome sequencing^9-11^, *in vitro* cloning of single cells^12,13^, microdissection or biopsy of clonal populations^14,15^, and duplex sequencing^16-19^—amplify the original DNA molecules before sequencing, either prior to or on the sequencer itself. Amplification of DNA prior to sequencing masks true ssDNA events by either transforming existing ssDNA mismatches and damage to dsDNA mutations, or by introducing artifactual ssDNA mismatches and damage^16^.

Mosaic dsDNA mutations are the result of the interaction between ssDNA mismatch and damage events, DNA repair, and DNA replication^5,20^. For example, dsDNA mutational signatures (i.e., the sequence contexts of mutations) may not reflect the patterns of the originating ssDNA events, but rather only of the ssDNA events that are misrepaired or unrepaired prior to replication^8^. dsDNA mutation profiling also does not resolve on which strands the initiating mutational processes are occurring. Therefore, a complete understanding of the process of mutation requires profiling of ssDNA mismatches and damage^5,21^. Here, to study the ssDNA origins of mosaic mutations, we developed an approach for direct sequencing of single DNA molecules without any prior amplification that achieves, for single-base substitutions, single-molecule fidelity detection of dsDNA mutations simultaneously with ssDNA mismatches and damage.

### Hairpin Duplex Enhanced Fidelity Sequencing (HiDEF-seq)

Profiling dsDNA mosaic mutations in human tissues requires single-molecule fidelity of < 1 error per 1 billion bases (10^-9^), and profiling ssDNA mismatch and damage events would likely require similar or greater fidelity^16,21-23^. However, no technology, to date, has achieved this fidelity when directly sequencing unamplified single DNA molecules. To achieve this, we developed Hairpin Duplex Enhanced Fidelity Sequencing (HiDEF-seq). HiDEF-seq dramatically increases the fidelity of single-molecule sequencing by: (1) increasing the number of independent sequencing passes for each molecule (median 32 passes with median 1.7 kilobase (kb) molecules) relative to standard single-molecule sequencing^24,25^ to create a high-quality consensus sequence for each strand; (2) eliminating *in vitro* artifacts during library preparation, initially using the NanoSeq A-tailing approach^16^ (HiDEF-seq version 1) and subsequently with an improved protocol that removes residual artifacts (HiDEF-seq version 2); and (3) a computational pipeline that avoids analytic artifacts (**Figs. 1a,b, Extended Data Figs. 1-5, and Methods**). HiDEF-seq libraries are sequenced on Pacific Biosciences (PacBio) single-molecule, long-read sequencers. The computational pipeline analyzes single base substitutions, since these have an orthogonal error profile to the prevalent insertion and deletion sequencing errors of single-molecule sequencing^26^, and it analyzes each strand separately to distinguish dsDNA from ssDNA events (**Methods**). Further, it utilizes the telomere-to-telomere human reference genome, which was itself constructed using long reads^27^. Germline variants can be filtered using either standard short-read or long-read genome sequencing of the same individual (**Extended Data Figs. 3k,l**).

**Figure 1.**
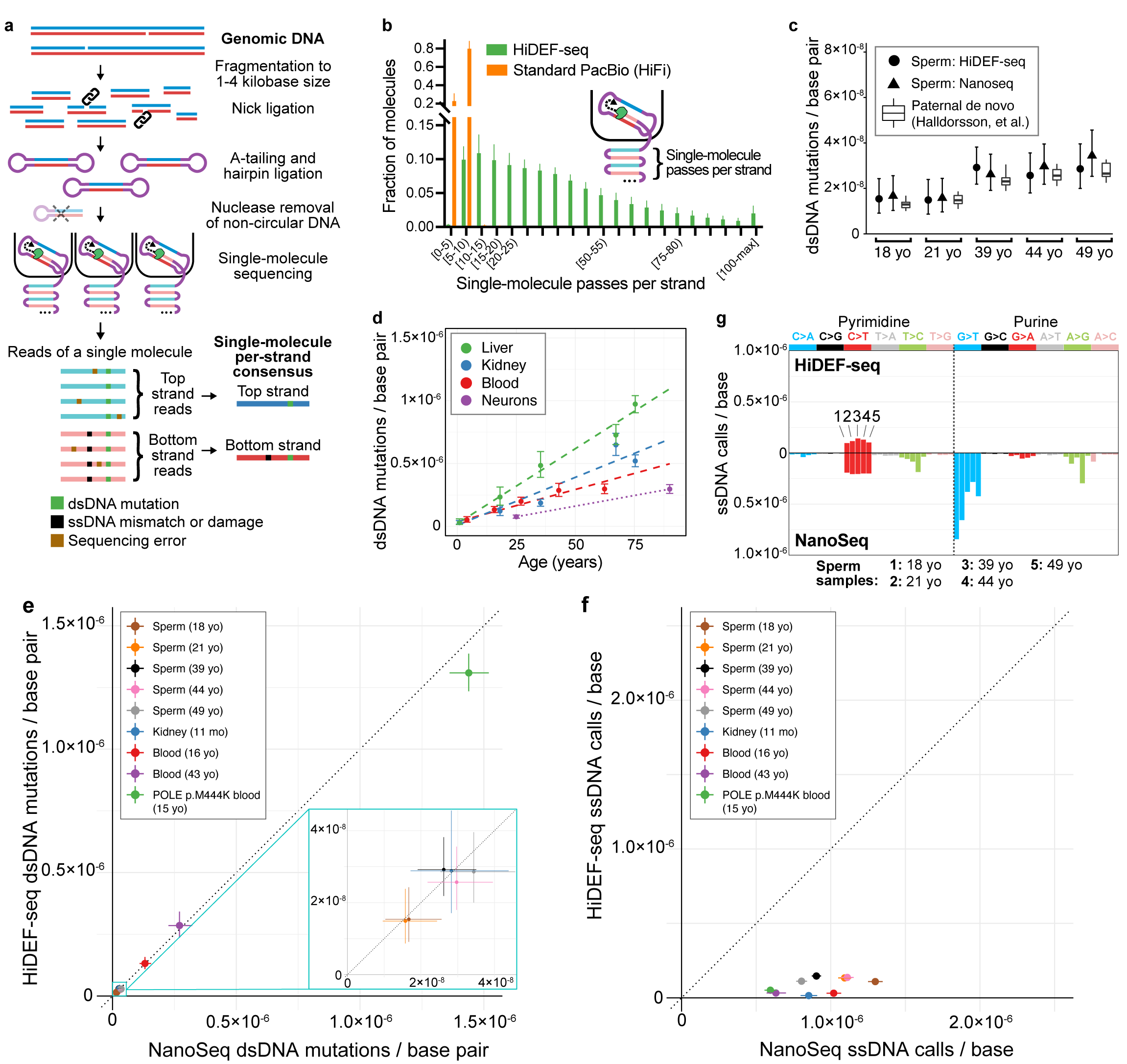
Overview of method. **a**, Schematic of library preparation and sequencing. A-tailing is performed with a polymerase, dATP and non-A dideoxynucleotides to block residual nicks^16^ (not illustrated), except for fragmented DNA samples that utilize only dideoxynucleotides (without dATP) in this step to avoid misincorporation of dATP at these samples’ more numerous residual nick sites (**Extended Data Fig. 5 and Methods**). Sequencing reads are reverse complements of the template molecule. **b**, Histogram of the average number of passes per strand (**Methods**) in single-molecule sequencing of representative HiDEF-seq samples (n=51) and standard Pacific Biosciences (PacBio HiFi) samples (n=10). The average percentage of molecules with ≥ 5 and ≥ 20 passes per strand is: 99.8% and 70% for HiDEF-seq, respectively, and 78.7% and 0.1% for HiFi, respectively. Plot shows HiDEF-seq molecules output by the primary data processing step of the analysis pipeline. X-axis square brackets and parentheses signify inclusion and exclusion of interval endpoints, respectively. **c,** dsDNA mutation burdens in sperm samples (left to right: SPM-1013, SPM-1002, SPM-1004, SPM-1020, SPM-1060) profiled by both HiDEF-seq v2 and NanoSeq, compared for each age (yo, years old) to paternally-phased de novo mutations in children from a prior study of 2,976 trios^28^. **d,** dsDNA mutation burdens versus age measured by HiDEF-seq v2 in samples from individuals without cancer predisposition. Dashed lines (liver, kidney, blood): weighted least-squares linear regression. Dotted line (neurons): these only connect two data points to aid visualization of burden difference, since regression cannot be performed with two samples. **e,** Comparison of HiDEF-seq versus NanoSeq dsDNA mutations per base pair for samples profiled by both methods. Samples (top to bottom in legend) are: SPM-1013, SPM-1002, SPM-1004, SPM-1020, SPM-1060, 1443, 1105, 6501, 63143. Note, all samples except for 63143 (POLE p.M444K) are from individuals without a cancer predisposition syndrome. Dashed line diagonal, y = x, is the expectation for concordance. **f,** Comparison of HiDEF-seq versus NanoSeq ssDNA calls per base for samples profiled by both methods. These are the same samples as in (**e**). **g,** Comparison of HiDEF-seq versus NanoSeq ssDNA calls per base, separated by call type. For each call type (i.e., C>A, C>G, etc.), each bar represents a different sperm sample. Samples for each call type, from left to right, are SPM-1013, SPM-1002, SPM-1004, SPM-1020, SPM-1060. **b,** Error bars: standard deviation. **c-f,** Error bars: Poisson 95% confidence intervals. **c,** Box plots: middle line, median; boxes, 1st and 3rd quartiles; whiskers, 5% and 95% quantiles. **c,e,f,** For each sample, HiDEF-seq and NanoSeq confidence intervals were normalized to reflect an equivalent number of interrogated base pairs (c,e) or bases (f) (**Methods**). **e-g,** yo, years old; mo, months old.

With the first version of HiDEF-seq (v1), we profiled purified human sperm as the most rigorous test of fidelity for detecting dsDNA mosaic mutations—since sperm harbor the lowest dsDNA mutation burden of any readily accessible human cell type^3^. Sperm dsDNA mutation burdens as measured by HiDEF-seq v1 were concordant with a prior study of de novo mutations^28^ and with NanoSeq profiling^16^ (a recently developed method for sequencing mosaic dsDNA mutations) that we performed for the same samples (**Extended Data Figs. 4a**). HiDEF-seq v1 also measured the expected dsDNA mutational signatures and increase in dsDNA mutation burden with age in other primary human tissues (kidney, liver, blood, and cerebral cortex neurons)^18,20^ (**Extended Data Figs. 4b,c**). Notably, relaxing from a threshold of ≥ 20 to ≥ 5 sequencing passes per strand, while keeping our optimized computational filters, produced concordant results (**Extended Data Fig. 3e**). This suggests that with our computational approach alone, PacBio sequencing can achieve a higher per-pass fidelity for substitutions than estimated by prior studies^24^. Accordingly, for ultra-high fidelity analysis of dsDNA mutations, we used this lower threshold of ≥ 5 passes per strand as this increases the percentage of molecules included in the analysis from 70% to 99.8% (of molecules that pass primary data processing), and it increases the percentage of bases that are interrogated by 11%. We also successfully quantified the dsDNA mutation burden of sperm using HiDEF-seq with larger DNA fragments (median 4.2 kb), which have correspondingly fewer passes per strand (median 15 passes) (**Supplementary Note 1**). However, for this study, we proceeded with HiDEF-seq with the smaller median 1.7 kb fragments, since a higher threshold of ≥ 20 passes per strand was required for ssDNA analysis.

Next, we proceeded to analyze ssDNA calls. Importantly, ssDNA calls may include not only ssDNA mismatches, but also damaged bases that alter base pairing properties and lead to mis-incorporation of nucleotides by the sequencer polymerase. The latter is potentially advantageous as it would enable high fidelity detection of ssDNA damage. However, whereas dsDNA mutation analysis can take advantage of information in both strands (duplex error correction) and its fidelity can be confirmed using the expected mutation burden of sperm, duplex error correction is not possible for ssDNA calls, and ssDNA mismatch burdens are unknown. Hence, for ssDNA calling we optimized key analytic parameters by identifying filter thresholds above which ssDNA burden estimates are stable and identifying any patterns that may suggest artifacts (**Extended Data Figs. 3i,j and Methods**).

Upon initial analysis of HiDEF-seq v1’s ssDNA calls, we identified that approximately 60% of calls were T>A changes at a motif corresponding to half of the recognition sequence of the restriction enzyme that we use to fragment the input DNA, likely because the enzyme normally operates as a dimer while creating rare ssDNA nicks as a monomer (**Extended Data Figs. 4d-e**). At these ssDNA nicks, mismatched deoxyadenosines can then be introduced during the A-tailing step of library preparation (**Extended Data Fig. 4f**). We were able to fully eliminate these artifactual ssDNA mismatches by adding a ssDNA nick ligation step after restriction enzyme fragmentation (**Extended Data Figs. 4d,f,g**). In a larger set of sperm samples, this improved version of HiDEF-seq (v2) again measured the expected dsDNA mutation burdens (**Fig. 1c**)^28^. The fidelity of HiDEF-seq v2 for dsDNA mutations (estimated as the probability of complementary single-strand mismatches occurring at the same position; **Methods**) is < 1 error per 7⋅10^16^ base pairs with ≥ 5 passes per strand and < 1 error per 10^17^ base pairs with ≥ 20 passes per strand.

Across all sample types profiled in this study with HiDEF-seq v2 (kidney, liver, blood, sperm, cerebral cortex neurons, primary fibroblasts, and lymphoblastoid cell lines), we found that post-mortem kidney and liver samples still exhibited a significant burden of T>A ssDNA calls despite this nick ligation step (average 67% of calls and average 1 call per 3.5 million bases) that correlated with the extent of post-mortem DNA fragmentation (**Extended Data Figs. 5a-c and Supplementary Table 3**). These presumed artifactual ssDNA T>A calls do not correspond to any recognizable sequence motif and likely arise from post-mortem ssDNA nicks that HiDEF-seq v2’s nick ligation step is unable to seal due to damaged nucleotides at the nick sites (**Extended Data Fig. 5d**). We trialed a variety of approaches to remove these T>A ssDNA artifacts so that HiDEF-seq v2 could also be used to profile post-mortem tissues with fragmented DNA (**Extended Data Fig. 5e and Methods**). We discovered that altogether removing A-tailing from the protocol completely eliminated these T>A ssDNA artifacts, and further, that retaining a polymerase nick extension step with non-A dideoxynucleotides^16^ removes low-level random-context artifactual ssDNA calls to produce final ssDNA call patterns similar to non-post mortem tissues and without any discernible artifacts (**Extended Data Figs. 5e-g and Supplementary Table 3**). We also profiled sperm with this non-A-tailing HiDEF-seq v2 protocol and confirmed its dsDNA mosaic mutation fidelity (**Extended Data Fig. 5h**). The only disadvantage to removing A-tailing from HiDEF-seq v2 is a requirement for approximately double the amount of input DNA due to the lower efficiency of subsequent blunt adapter ligation (**Methods**). We therefore utilized standard HiDEF-seq v2 for nearly all samples, except for post-mortem kidney, post-mortem liver, and tumor samples for which we utilized HiDEF-seq v2 without A-tailing (**Supplementary Table 1**).

Similar to HiDEF-seq v1, profiling of primary human tissues (kidney, liver, blood, and cerebral cortex neurons) with HiDEF-seq v2 exhibited the expected dsDNA mutational signatures and linear increase in dsDNA mutation burden with age^16,18^ (**Fig. 1d and Extended Data Fig. 5i**). For simplicity, unless otherwise specified, we subsequently refer to HiDEF-seq v2 (both with and without A-tailing) as HiDEF-seq.

To compare ssDNA calls between HiDEF-seq and NanoSeq, we profiled 9 samples with both methods. While HiDEF-seq and NanoSeq dsDNA mutation burdens and patterns were concordant, HiDEF-seq measured on average 18-fold lower ssDNA call burdens than Nanoseq, with distinct patterns, and 5-fold lower when considering only C>T calls (**Figs. 1e-g and Extended Data Figs. 6a-c**). This suggests that while NanoSeq achieves ultra-high fidelity for dsDNA mutations, its ssDNA calls are largely artifactual as suggested by its developers^16^. The HiDEF-seq ssDNA burden measured for cerebral cortex neurons was also ∼13-fold lower than estimated by the recently developed Meta-CS single-cell duplex sequencing method^29^, with a distinct pattern, and ∼4-fold lower when considering only C>T calls (**Supplementary Tables 2-3**). Altogether, by direct interrogation of unamplified single molecules, HiDEF-seq achieves the highest fidelity for single base changes of any DNA sequencing method to date.

### Single-strand mismatch patterns in cancer predisposition syndromes

Since there is no prior method for sequencing ssDNA mismatches with single-molecule fidelity, we sought to confirm the veracity of HiDEF-seq’s ssDNA calls by profiling samples and cell lines from individuals with inherited cancer predisposition syndromes that may have elevated ssDNA mismatch burdens. We profiled with HiDEF-seq 17 blood, primary fibroblast, and lymphoblastoid cell line samples from 8 different cancer predisposition syndromes, including defects in nucleotide excision repair, mismatch repair, polymerase proofreading, and base excision repair (**Supplementary Tables 1-2**). In these samples, we first confirmed HiDEF-seq’s fidelity for dsDNA mutations by measuring the expected dsDNA mutation burdens and signatures based on prior studies^30-34^—except for *MUTYH* blood samples from which we were unable to recover its known signatures, since as seen in prior studies, *MUTYH* blood has near normal mutation burdens^34^ (**Extended Data Figs. 7a-d, and Supplementary Tables 2 and 4**). In *ERCC6* and *ERCC8* mutant cell lines, whose mutational patterns are unknown, we identified a signature similar to the COSMIC^35^ SBS36 signature (SBS, single base substitution; cosine similarity 0.82) (**Extended Data Figs. 7b-c**). These data further illustrate the single-molecule fidelity of HiDEF-seq for dsDNA mutations.

Notably, compared to non-cancer predisposition samples, we detected an increase in ssDNA calls per base in two cancer predisposition syndromes: a 2.6-fold increase (95% confidence interval 2.3-3.0, p<10^-15^, Poisson rates ratio test) in *POLE* polymerase proofreading-associated polyposis syndrome samples (PPAP; germline heterozygous exonuclease domain mutations in *POLE*, which encodes polymerase epsilon that is responsible for leading strand genome replication^36,37^), and a 1.6-fold increase (95% confidence interval 1.4-1.9, p = 8⋅10^-11^) in congenital mismatch repair deficiency syndrome samples (CMMRD; *MSH2*, *MSH6*, and *PMS2* germline bi-allelic loss-of function) (**Fig. 2a**).

**Figure 2.**
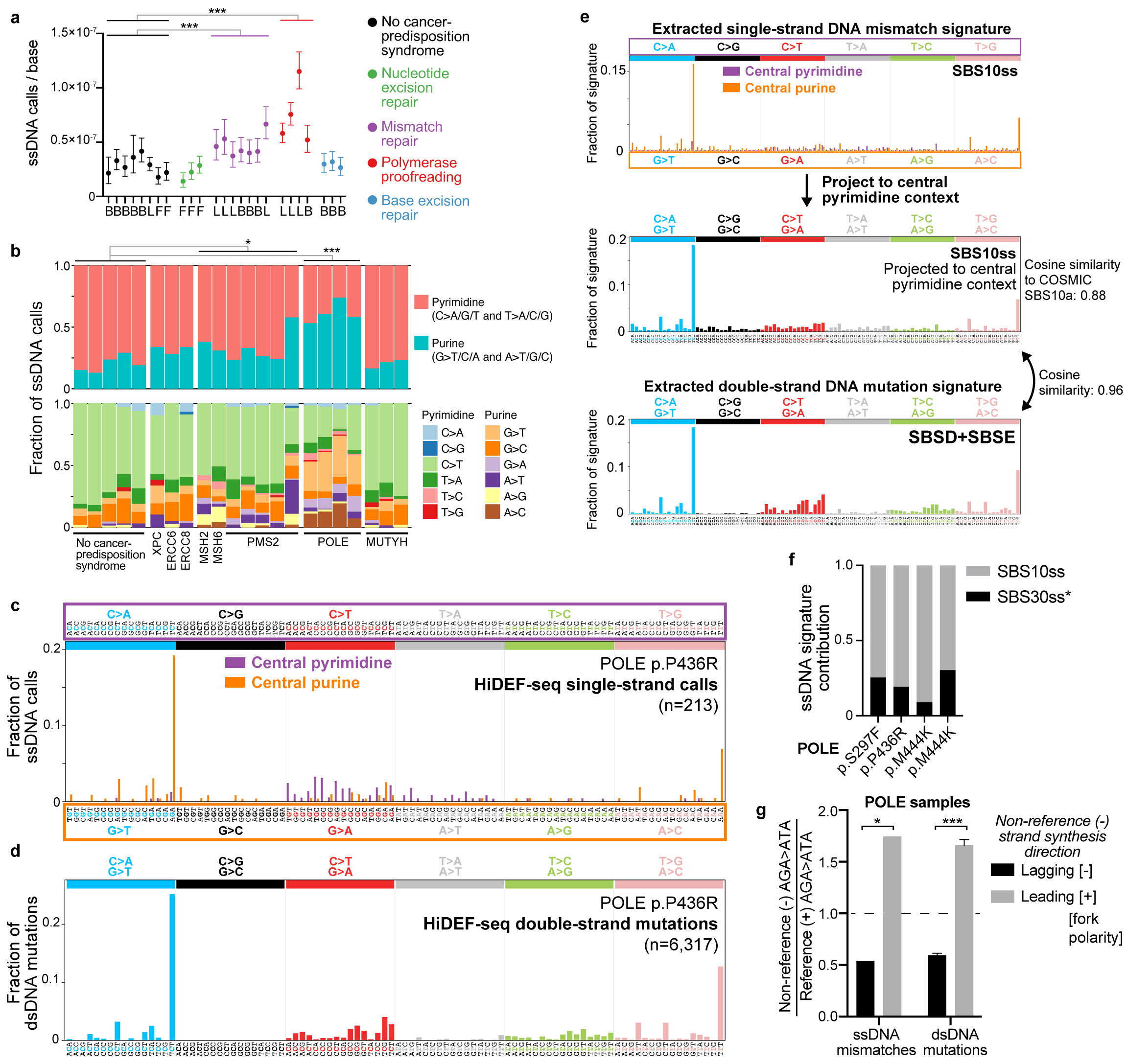
ssDNA call burdens and patterns in cancer-predisposition syndromes. **a,** Burdens of ssDNA calls in blood (B), fibroblasts (F), and lymphoblastoid cell lines (L) from individuals without and with cancer predisposition syndromes. Call burdens are corrected for trinucleotide context opportunities and detection sensitivity (**Methods**). ***, p = 8⋅10^-11^ for mismatch repair versus non-cancer predisposition samples and p < 10^-15^ for polymerase proofreading versus non-cancer predisposition samples (Poisson rates ratio test, using combined counts of calls and interrogated bases from each group). Results were still significant when including only blood samples. From left to right, non-cancer predisposition samples are: 5203, 1105, 1301, 6501, 1901, GM12812, GM02036, GM03348; cancer predisposition samples are: GM16381, GM01629, GM28257, 55838, 58801, 57627, 1400, 1324, 1325, 60603, 59637, 57615, 63143 (L), 63143 (B), CC-346-253, CC-388-290, CC-713-555. For cancer predisposition samples, the affected genes are in the same left-right order as for cancer predisposition samples in (b). **b,** Fraction of ssDNA call burdens by context, corrected for trinucleotide context opportunities. We include only non-cancer predisposition samples with > 30 ssDNA calls (1105, 1301, 1901, GM12812, GM03348) for reliable fraction estimates. However, the cancer predisposition sample GM16381 (XPC) with < 30 ssDNA calls is included for completeness to show all cancer predisposition samples. The cancer predisposition syndrome samples are in the same order as in (a). **c,d**, ssDNA (c) and dsDNA (d) call spectra for representative *POLE* sample 57615, corrected for trinucleotide context opportunities. Parentheses show total number of calls. **e,** Top, ssDNA mismatch signature SBS10ss extracted from all *POLE* samples. The signature was extracted de novo while simultaneously fitting SBS30ss* (see **Fig. 4e**). Middle, SBS10ss projected to central pyrimidine context by summing central pyrimidine and their reverse complement central purine values to allow comparison to dsDNA signatures. Bottom, dsDNA mutational signature (sum of SBSD and SBSE) extracted de novo from all *POLE* samples, while simultaneously fitting SBS1 and SBS5. **f,** Fraction of ssDNA calls attributed to each ssDNA signature in *POLE* samples (left to right): 59637, 57615, and 63143 lymphoblastoid cell lines, and 63143 blood. Protein-level POLE mutation is annotated below. Cosine similarities of original spectra of samples to spectra reconstructed from component signatures are (left to right): 0.94, 0.97, 0.97, 0.85. See **Fig. 4e** for details of SBS30ss*. **g,** In *POLE* samples, AGA>ATA ssDNA mismatches and AGA>ATA dsDNA mutations occur more often on the non-reference (-) than on the reference (+) strand in regions where the non-reference strand is synthesized more frequently in the leading direction (i.e., positive fork polarity), based on replication timing data (**Methods**). Reference (+) strand refers to the plus strand of the human reference genome. See **Extended Data Fig. 7e** for plots of dsDNA mutations separated by fork polarity quantiles (rather than positive versus negative polarity), which cannot be plotted for ssDNA mismatches due to the low number of ssDNA mismatches per quantile. Y-axis is the ‘strand ratio’, calculated as the fraction of all AGA>ATA non-reference strand events that have the specified fork polarity divided by the fraction of all AGA>ATA reference strand events that have the specified fork polarity. For ssDNA analysis, the strand ratio is calculated using the ssDNA mismatches of all *POLE* samples, since there are not enough ssDNA mismatches to quantify this reliably for each sample separately. For dsDNA analysis, strand ratios were calculated for each sample separately, and the plot shows average and standard deviation (error bars) across these samples. Dashed line at 1.0 is the expected ratio in the absence of strand asymmetry. *, p = 0.015 (chi-squared test, n = 73 ssDNA AGA>ATA mismatches); ***, p < 10^-15^ (chi-squared test of all 3,871 dsDNA AGA>ATA mutations across all *POLE* samples). An analysis excluding mismatches and mutations overlapping genes, to exclude biases due to transcription strand, was still significant for dsDNA mutations (p < 10^-15^) but not for ssDNA mismatches, but this analysis has significantly reduced power due to the 55% reduction in the number of ssDNA mismatches remaining for analysis. **a,b**, See further disease and sample details, including genotypes, in **Supplementary Tables 1-2. a**, Error bars, Poisson 95% confidence intervals.

Next, we examined the patterns of ssDNA calls. The percentage of purine ssDNA calls (G>T/C/A and A>T/G/C) was elevated in PPAP samples to an average of 61% (range 53-74%) compared to 20% (range 13-29%) in non-cancer-predisposition samples (**Fig. 2b**; p = 0.0004, heteroscedastic two-tailed t-test; analysis excludes non-cancer predisposition samples with less than 30 ssDNA calls as their call patterns are not reliably ascertained). This increase in purine ssDNA calls in PPAP was largely due to an increase in the fraction of G>T, G>A, and A>C ssDNA calls (**Fig. 2b**). There was no significant correlation of ssDNA call contexts with specific *POLE* mutations in PPAP samples (**Fig. 2b and Supplementary Table 3**). The percentage of purine ssDNA calls was also elevated to a lesser degree in CMMRD samples to an average of 33% (range 23-58%, p = 0.04), though without a clear enrichment of a specific sequence context except for one *PMS2* loss-of-function sample with increased A>T ssDNA calls (**Fig. 2b**). These data indicate that most ssDNA calls in PPAP samples, and at least some calls in CMMRD samples, are bona fide ssDNA mismatches.

To further characterize the patterns of ssDNA mismatches in *POLE* PPAP samples, we plotted their 192-trinucleotide context spectra (standard 96-trinucleotide context spectrum, separated by central pyrimidine versus central purine). This revealed a distinct pattern, with two large peaks for AGA>ATA and AAA>ACA accounting for ∼15-20% and ∼5-10% of ssDNA mismatches, respectively, in addition to smaller peaks with G>T, G>A, A>C, and C>T contexts (**Fig. 2c and Supplementary Table 3**). The ssDNA mismatch spectra were highly concordant with these same samples’ dsDNA mutation spectra (**Fig. 2d and Supplementary Table 4**), confirming these are true ssDNA mismatches and that these initial mismatch events—due to polymerase epsilon nucleotide misincorporation—lead to the subsequent pattern of accumulated dsDNA mutations. We then performed de novo extraction of ssDNA mismatch signatures from *POLE* PPAP samples, which produced a signature we name SBS10ss (ss, single-strand) (**Fig. 2e**). Note, as this is the first ssDNA signature, we propose a nomenclature with suffix ‘ss’ to distinguish ssDNA from dsDNA signatures. Projecting SBS10ss to central pyrimidine contexts, by summing central purine and central pyrimidine spectra, produced a spectrum remarkably similar (cosine similarity 0.96) to the dsDNA signatures extracted de novo (SBSD+SBSE) from these same samples (**Fig. 2e**), again indicating that the ssDNA mismatches are the inciting events subsequently leading to dsDNA mutations. SBS10ss also had high similarity to COSMIC SBS10a (cosine similarity 0.88) that has been previously associated with *POLE* PPAP^30^. SBS10ss accounted for an average of 79% (range 75 – 91%) of ssDNA calls in *POLE* PPAP samples, with the remaining attributed to SBS30ss*, a ssDNA cytosine deamination damage signature (*, indicates damage) described in a subsequent section (**Fig. 2f**). For CMMRD samples, the number of ssDNA calls was too low to extract a signature.

The two most frequent ssDNA mismatch contexts in *POLE* PPAP samples are also notable for the asymmetry of their prevalence relative to their reverse complements: AGA>ATA versus TCT>TAT (73 vs. 10 mismatches across all *POLE* samples; chi-squared p < 0.0001) and AAA>ACA versus TTT>TGT (26 vs. 2 mismatches; chi-squared p < 0.0001). These data provide the first direct observation that the dsDNA mutational context of AGA>ATA / TCT>TAT that is prevalent in *POLE* PPAP arises significantly more frequently from C:dT (template base:polymerase incorporated base) misincorporations rather than G:dA misincorporations, and that the dsDNA mutational context of AAA>ACA / TTT>TGT arises more frequently from T:dC than A:dG misincorporations. Importantly, these results are consistent with prior studies that indirectly inferred this asymmetry using yeast^38^ and human tumors^39-41^ harboring polymerase epsilon exonuclease domain mutations by identifying asymmetries in the prevalence of dsDNA mutation contexts relative to their reverse complement contexts depending on whether the mutation locus is preferentially replicated via leading versus lagging strand synthesis. In contrast to these studies that rely on replication timing data that imperfectly estimates the probability of leading versus lagging strand replication in a bulk sample to measure this asymmetry, our single-molecule detection of nucleotides that were misincorporated *in vivo* by replicative polymerases allows us to measure this asymmetry directly. We also applied the above studies’ indirect replication timing approach and similarly found replication strand asymmetry for our *POLE* PPAP samples’ AGA>ATA dsDNA mutations (**Fig. 2g and Extended Data Fig. 7e**). We further show that the AGA>ATA ssDNA mismatches in these samples occur more frequently on the strand that is synthesized in the leading, rather than lagging direction, consistent with the role of polymerase epsilon in leading strand synthesis^36,37^ (**Fig. 2g**). Altogether, these results represent the first direct measurements of *in vivo* ssDNA mismatch burdens and patterns.

### Single-strand mismatch patterns of tumors deficient in both mismatch repair and polymerase proofreading

To further study the interaction between ssDNA mismatches introduced during replication and mismatch repair, we profiled 3 brain tumors from individuals with CMMRD whose tumors also harbored somatic mutations affecting polymerase proofreading. One of the tumors (Tumor 3) was excluded from further analysis as it had a very high ssDNA C>T burden attributed to SBS30ss*, a ssDNA cytosine deamination damage signature described in the next section that likely arose from *ex vivo* DNA damage (**Supplementary Tables 2 and 3**). The other two tumors, a medulloblastoma and a glioblastoma—both with bi-allelic germline *PMS2* mutations and heterozygous somatic *POLE* exonuclease domain mutations—had higher burdens and different patterns of dsDNA mutations and ssDNA calls than samples deficient in only mismatch repair or only polymerase proofreading (**Figs. 2a-d, 3a-d, Extended Data Figs. 7a-b, and Supplementary Tables 2-3**). Additionally, the tumors’ dsDNA mutation spectra matched those found in prior studies of tumors and cell lines deficient in both mismatch repair and polymerase proofreading (**Fig. 3d**)^42-45^. Most dsDNA mutations were attributed to a signature we extracted *de novo* that most resembled COSMIC SBS14 (cosine similarity 0.85) (**Fig. 3f**)^44^.

**Figure 3.**
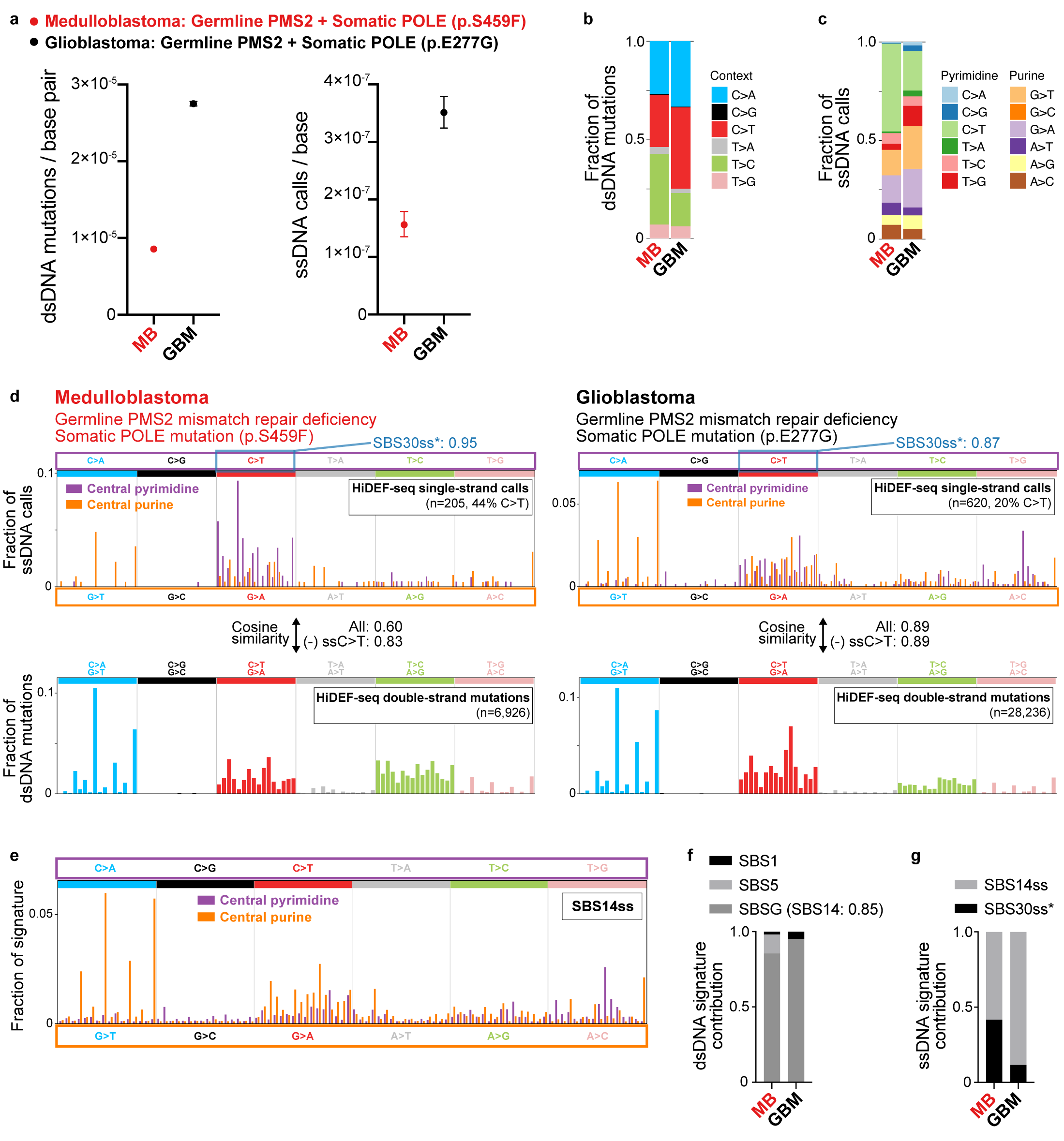
Tumors deficient in both mismatch repair and polymerase proofreading. **a,** Burdens of dsDNA mutations (left) and ssDNA calls (right) in a medulloblastoma (ID: Tumor 8) and glioblastoma (ID: Tumor 10). See **Supplementary Table 1** for sample details. Burdens are corrected for trinucleotide context opportunities and detection sensitivity (**Methods**). **b,c,** Fraction of dsDNA mutation burdens (b) and ssDNA call burdens (c) by context, corrected for trinucleotide context opportunities. **d,** Spectra of ssDNA calls (top) and dsDNA mutations (bottom) in tumor samples corrected for trinucleotide context opportunities. Parentheses show the total number of raw calls, and the percentage of calls that are C>T after correction for trinucleotide context opportunities. Blue annotation on the top right of each ssDNA spectrum is the cosine similarity of only the ssDNA C>T calls to SBS30ss* (see **Fig. 4e** for details of SBS30ss*). Also annotated are the cosine similarities of each sample’s full ssDNA call spectrum (projected to central pyrimidine context) to its dsDNA mutation spectrum, for all ssDNA calls and excluding ssDNA C>T calls (most of which are due to the SBS30ss* cytosine deamination process). **e,** ssDNA mismatch signature SBS14ss extracted from tumor samples. The signature was extracted de novo while simultaneously fitting SBS30ss*. **f,** Fraction of dsDNA mutations attributed to each dsDNA signature in tumor samples. Cosine similarity of the de novo extracted signature SBSG to the best matching COSMIC SBS signature is shown in parentheses. Cosine similarities of original spectra of samples to spectra reconstructed from component signatures are (left to right): 0.94 and 0.998. **g,** Fraction of ssDNA calls attributed to each ssDNA signature in tumor samples. Cosine similarities of original spectra of samples to spectra reconstructed from component signatures are (left to right): 0.91 and 0.98. **a,** Error bars, Poisson 95% confidence intervals. **a-c,f,g,** MB, medulloblastoma (ID: Tumor 8); GBM, glioblastoma (ID: Tumor 10).

Importantly, the tumors’ ssDNA call spectra largely matched their dsDNA mutation spectra (**Figs. 3b-d**)—except for ssDNA C>T calls related to SBS30ss* (**Figs. 3d,g**). Consequently, the tumors’ ssDNA call spectra harbored notable differences relative to ssDNA call spectra of samples deficient in only mismatch repair or polymerase proofreading, including increases in ssDNA AG>AT calls flanked by 3’ C/G/T, and increases in ssDNA G>A, A>G, and T>C calls (**Figs. 2c, 3d**). Note that the relative increase in ssDNA C>T calls in the tumors largely arose from cytosine deamination damage rather than polymerase misincorporation (**Figs. 3d,g and 4e**). These differences in ssDNA call spectra of polymerase proofreading-deficient samples with and without mismatch repair deficiency are consistent with prior studies suggesting that that the mismatch repair system is more efficient at repairing certain mismatches caused by deficient polymerase proofreading^45-47^. The ssDNA call spectra further resolve the identity of the nucleotides misincorporated by proofreading-deficient polymerase epsilon that lead to the tumors’ observed dsDNA mutation spectrum—for example, these data indicate that the C>T / G>A dsDNA mutations of COSMIC SBS14 largely arise from C:dA (template base:polymerase incorporated base) misincorporations rather than G:dT misincorporations. We extracted a ssDNA mismatch signature from tumor samples that we name SBS14ss, since its most similar COSMIC dsDNA signature is SBS14 (cosine similarity 0.73, after projecting SBS14ss to central pyrimidine contexts) (**Fig. 3e**). SBS14ss accounted for most ssDNA calls in both tumors, with the remaining attributed to SBS30ss* (**Fig. 3g**).

### Single-strand patterns of cytosine deamination damage

We reasoned that HiDEF-seq may also enable detection of rare ssDNA damage events with single-molecule fidelity—specifically, base damage that leads to nucleotide misincorporation by the sequencer polymerase. Detecting these rare events would be useful for characterizing processes that damage DNA. A common form of DNA damage that leads to mutations is the deamination of cytosine (with or without preceding oxidation) to uracil, uracil glycol, 5-hydroxyuracil, or 5-hydroxyhydantoin (uracil-species)^48-54^. When these lesions are not repaired, they result in dsDNA C>T transition mutations^48^. We hypothesized that HiDEF-seq may detect these ssDNA cytosine to uracil-species events despite their low levels (estimated by mass spectrometry at < 1 per 1 million bases in mammalian cells^55^), since the damaged cytosines would be mis-sequenced as thymine.

We began by investigating the burden and pattern of ssDNA C>T calls in blood DNA of normal (i.e., non-cancer predisposition) individuals, since blood can be processed rapidly without potential post-mortem DNA damage. We also extracted the DNA with only room temperature incubations to avoid heat-induced deamination damage^56^. Blood DNA had 2.0⋅10^-8^ ssDNA C>T calls per base (mean of n=9 samples from n=5 individuals; range 9.6⋅10^-9^ – 3.1⋅10^-8^), which comprised on average 71% of these samples’ ssDNA calls (**Extended Data Fig. 8a and Supplementary Tables 2 and 3**). This burden, which may have either been present *in vivo* or may have partly arisen during laboratory processing of the blood or DNA, suggests there are fewer than 200 cytosine to uracil-species deaminated bases per cell in blood leukocytes. This level of detection at 1 event per 50 million bases is on par with the most sensitive mass spectrometry methods^55,57,58^—which cannot determine the sequence context of damaged bases—and provides a low background for studying cytosine deamination processes. Notably, combining all the ssDNA calls across these samples and projecting their ssDNA trinucleotide context spectrum to the corresponding dsDNA trinucleotide spectrum produced a spectrum similar to COSMIC^35^ SBS30 (cosine similarity 0.83) (**Figs. 4a,c**), a signature associated with cytosine oxidative deamination damage repaired by DNA glycosylases^31,59-62^. Surprisingly, there was no signal in these blood samples for the commonly oxidized base 8-oxoguanine that would be expected to lead to G>T ssDNA calls, which were very infrequent in these blood samples (average 6% of ssDNA calls; 1.6⋅10^-9^ ssDNA calls per base; range 0 – 2.9⋅10^-9^). This is likely due to the sequencer polymerase (a derivative of phi29 polymerase) incorporating correctly dC rather than misincorporating dA across from 8-oxoguanine bases^63^.

**Figure 4.**
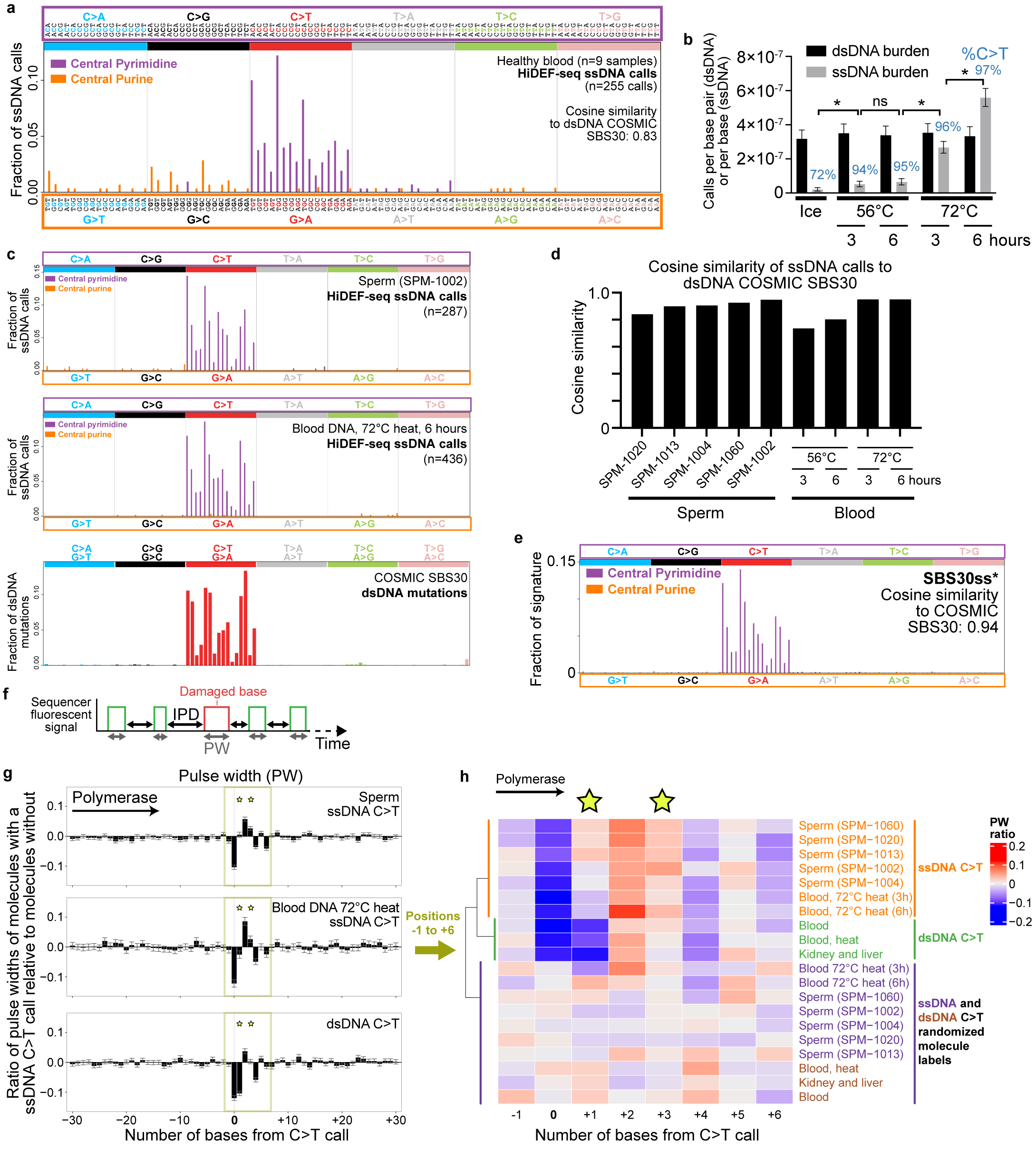
ssDNA damage signatures in sperm and after heat treatment. **a,** Spectrum of all ssDNA calls of non-cancer predisposition (healthy) blood samples (1 sample each from individuals 1105, 1301, 5203, 6501, and 5 samples from individual 1901). Cosine similarity to the dsDNA COSMIC signature SBS30 is calculated after projecting the ssDNA spectrum to a central pyrimidine trinucleotide spectrum (by summing values of central pyrimidine and their reverse complement central purine contexts). **b,** dsDNA mutation and ssDNA call burdens of heat-treated DNA. Non-heat-treated DNA was placed on ice for 6 hours. DNA heat-treated for 3 hours was subsequently placed on ice for 3 hours. The percentage of ssDNA sequencing calls that are C>T are annotated above each sample. **c,** Spectra of ssDNA calls for representative sperm and heat-treated blood DNA samples, and COSMIC SBS30 for comparison. **d,** Cosine similarity of ssDNA call spectra of each individual sperm and heat-treated blood sample to COSMIC SBS30, after projecting ssDNA calls to central pyrimidine trinucleotide contexts. **e,** SBS30ss* obtained by de novo signature extraction from central pyrimidine ssDNA calls of sperm and heat-treated samples. Cosine similarity to SBS30 is calculated after projecting to central pyrimidine trinucleotide context. **f,** Schematic of pulse width (PW) and interpulse duration (IPD) measured by the sequencer for each incorporated base. **g,** Average ratio of pulse widths at C>T call locations and 30 flanking bases of each molecule with a C>T call relative to molecules aligning to the same locus without the call. Data shows the average of the ratios for all ssDNA C>T calls in sperm samples (n=1799 calls), blood DNA samples that were heat-treated at 72C for 3 or 6 hours (n=626 calls), and dsDNA C>T mutations in a larger set of samples (non-heat treated blood DNA, 56C and 72C heat treated blood DNA, sperm, kidney, and liver; n=1217 mutations). The distinct profile of ssDNA C>T calls versus dsDNA C>T mutations, most notably at positions +1 and +3 (stars), indicates the ssDNA calls are damaged cytosines rather than cytosine to thymine mutations. **h,** Heat map of average pulse width ratios for C>T ssDNA calls and C>T dsDNA mutations for positions −1 to +6. Unbiased clustering of kinetic profiles (dendrogram) separates ssDNA from dsDNA calls and from kinetic profiles after randomizing labels of molecules with and without the calls. ssDNA ‘Blood, 72C heat (3h and 6h)’ (h, hours): heat-treated blood DNA. dsDNA ‘Blood, heat’: blood DNA heat-treated at 56C and 72C (both 3h and 6h for each); dsDNA ‘Blood’: 4 samples, not heat treated. dsDNA ‘Kidney and liver’: 10 samples, not heat treated. Star indicates positions +1 and +3 that best discriminate ssDNA C>T damage from dsDNA C>T mutations. **b**, Error bars, Poisson 95% confidence intervals. *, p < 0.005; ns, p>0.05; Poisson rates ratio test. **a,c-e**, HiDEF-seq spectra are corrected for trinucleotide context opportunities (**Methods**). **g**, Error bars, standard error of the mean.

Given the high sensitivity of HiDEF-seq’s ssDNA C>T detection, we sought to further elucidate the pattern of cytosine deamination damage via a larger number of events by investigating the effect of heat, an important source of laboratory-based cytosine deamination artifacts (since most DNA extraction methods utilize heat)^56,64^. We profiled purified blood DNA after heat incubation at 56 °C and 72 °C, each for 3 and 6 hours. While heat did not affect the dsDNA mutation burden, HiDEF-seq measured a significant increase in ssDNA calls (29-fold for 72 °C, 6 hour treatment), specifically C>T calls, with increasing temperature and time (**Fig. 4b and Supplementary Tables 2 and 3**). The effect of temperature was larger than the effect of time, and increased time had a larger effect at higher temperature (**Fig. 4b**). After 56 °C and 72 °C heat treatments, 94% and 97% of ssDNA calls, respectively, were C>T. Observation of this effect of heat led us to profile almost all samples in this study except for four samples (neurons of individual 5344 and 3 tumor samples) at least once with a room temperature DNA extraction (i.e., without heat incubation) (**Methods and Supplementary Table 1**). Notably, the temperature during HiDEF-seq library preparation does not exceed 37 °C (**Methods**).

Across all the healthy tissues and cell lines that we profiled, only sperm had a similarly high percentage (average 94%) of ssDNA calls that were C>T (**Extended Data Fig. 8a**). Sperm also had a high ssDNA C>T burden relative to other sample types (average 1.4⋅10^-7^ C>T calls per base) (**Extended Data Fig. 8a**). This suggests these are also cytosine deamination events and that sperm DNA either undergoes a greater degree of *in vivo* cytosine deamination than DNA of other tissues, or that it incurs this damage *ex vivo* prior to sperm purification from semen, during sperm purification, and/or during DNA extraction. To distinguish among these possibilities, we processed blood DNA with a known low ssDNA C>T burden with the same process used to extract DNA from sperm (**Methods**). This did not produce an increase in ssDNA C>T burden, indicating that our method of sperm DNA extraction is not the cause of sperm DNA’s high burden of ssDNA C>T events (**Supplementary Table 2**). To assess the possible contribution of the sperm purification process to the ssDNA C>T burden^65^, we purified sperm from semen of two additional individuals via filter chips that mimic physiologic separation of motile sperm, in parallel with the standard density gradient centrifugation method we used for the prior sperm samples (**Methods**). For each individual, sperm purified via the filter chip and sperm purified via density gradient centrifugation harbored the same percentage of ssDNA calls that were C>T (97% vs. 97% for the two methods for the first individual, and 87% vs. 87% for the second individual) and similar ssDNA C>T call burdens (1.4⋅10^-7^ vs. 1.1⋅10^-7^ for the first individual, and 9.0⋅10^-8^ vs. 9.0⋅10^-8^ for the second individual; p > 0.05 for both comparisons) (**Supplementary Tables 2-3**). These results suggest that the higher cytosine deamination burden in sperm occurs either *in vivo* or *ex vivo* during the time (< 1 hour) that semen liquefies in the laboratory prior to sperm purification. In both cases, the elevated cytosine deamination burden in sperm would likely be present in sperm fertilizing the egg, which would then be repaired by the DNA repair machinery of eggs after fertilization^66,67^.

Next, we analyzed the patterns (i.e., trinucleotide context spectra) of ssDNA C>T calls of sperm and heat-treated blood DNA samples. Strikingly, all sperm and heat-treated samples exhibited very similar ssDNA C>T spectra, and moreover, after projecting these ssDNA spectra to their corresponding dsDNA trinucleotide spectra, they again closely matched the COSMIC dsDNA signature SBS30 (average cosine similarity 0.90 and 0.95 for sperm and 72 °C heat samples, respectively) (**Figs. 4c,d**). Using all the above sperm and heat damage samples, we then extracted this ssDNA signature, which we term SBS30ss* (ss, single-strand; * indicates damage; cosine similarity 0.94 to SBS30) (**Fig. 4e**). The COSMIC dsDNA signature SBS30 has been previously associated with *NTHL1* and *UNG* biallelic loss of function mutations^31,59,60^ and with formalin fixation^68^. *NTHL1* and *UNG* encode DNA glycosylases that initiate base excision repair of oxidized pyrimidines, including uracil-species that result from cytosine oxidation^61,62^. Our finding that *in vitro* heat treatment of purified DNA leads to a ssDNA damage signature, SBS30ss*, that matches the *in vivo* dsDNA SBS30 signature, indicates that SBS30ss* and SBS30 reflect the nucleotide context bias of the primary biochemical process of cytosine deamination, likely via an oxidized intermediate, rather than a bias of base excision repair glycosylases to repair some trinucleotide contexts more efficiently. Moreover, the correspondence of SBS30ss* to SBS30 indicates that *in vitro* heat damage and formalin fixation converge on the same *in vivo* biochemical process that is revealed by loss of *NTHL1* and *UNG*.

To further confirm that ssDNA C>T calls in heat-treated DNA and sperm DNA are cytosine deamination damage (i.e., uracil-species) rather than ssDNA changes of cytosine to thymine, we took advantage of the single-molecule sequencer’s real-time polymerase kinetic data that records at a 10 millisecond frame rate both the duration of each individual nucleotide incorporation (pulse width, PW) and the time between consecutive nucleotide incorporations (inter-pulse duration, IPD) (**Fig. 4f**). The patterns of PW and IPD as the sequencing polymerase replicates and traverses a damaged base across the polymerase’s footprint of ∼8 nucleotides, encodes a unique kinetic signature for each canonical and damaged base^69,70^. Kinetic signatures have been previously identified for diverse base modifications in synthetic oligonucleotides, and they have been used to detect a small number of base modifications in genomic DNA such as cytosine methylation^69,71^. However, this approach has not yet been used to detect uracil-species in genomic DNA with single-molecule fidelity.

We began this kinetic analysis by extracting PW and IPD measurements from all ssDNA C>T calls of sperm and heat damage samples. We then controlled for local sequence context by normalizing the kinetic data of each molecule that has a C>T call using the kinetic data of all other molecules (across all samples) without C>T calls that aligned to the same locus (**Methods**). In parallel, we performed the same analysis for dsDNA C>T mutations (for the strand containing the thymine), as these are bona fide cytosine to thymine changes rather than cytosine damage. This analysis revealed a distinct kinetic signature for ssDNA C>T calls that differed from that of dsDNA C>T mutations (**Fig. 4g and Extended Data Fig. 8b**). Unbiased hierarchical clustering of PW of the −1 to +6 positions, which corresponds to the polymerase footprint in which the kinetic signal differs from the flanking baseline, separated the kinetic profile of individual samples’ ssDNA C>T calls from dsDNA C>T mutations (**Fig. 4h and Extended Data Fig. 8f**). Randomizing molecule labels abrogated these kinetic signals, further confirming their validity (**Fig. 4h and Extended Data Fig. 8c**). These results provide further evidence that the ssDNA C>T calls are uracil-species arising from cytosine deamination and definitively exclude the possibility that they are cytosine to thymine changes.

We further tested heat treatment of DNA in 5 different buffers and in water. All salt-containing buffers produced a similar burden of cytosine to uracil-species damage and the same SBS30ss* pattern as described above; however, samples that were heat-treated in water or in Tris-buffer without additional salt had a further 65-fold increase in cytosine damage (**Extended Data Fig. 8d and Supplementary Table 2**). Moreover, in these two samples, the damage pattern still closely matched SBS30ss*, except for an increased burden at 3’ T trinucleotide contexts and a decrease in burden for 5’ C and 3’ G contexts (**Extended Data Figs. 8e**). Since low or absent salt decreases DNA duplex stability at elevated temperatures and makes DNA more susceptible to deamination damage^72,73^, these results suggest that the *in vivo* mechanism of SBS30 and its corresponding SBS30ss* ssDNA signature is cytosine deamination of DNA while it is transiently single-stranded.

### Single-strand DNA calls in healthy tissues

We examined the burdens and patterns of ssDNA calls across the 29 healthy (i.e., non-cancer predisposition) samples that we profiled from sperm, liver, kidney, blood, cerebral cortex neurons, primary fibroblasts, and a lymphoblastoid cell line (n = 2,893 calls; 83% C>T). Except for sperm that exhibit significantly elevated ssDNA C>T calls from cytosine deamination damage as described above, we did not observe significant differences in ssDNA call burdens among tissue types (**Fig. 5a**). Liver samples, but not other tissues, showed a small but statistically significant increase in ssDNA call burden with age (5.8⋅10^-10^ calls per year; p = 0.0005) (**Fig. 5b**), and this correlation with age decreased but remained statistically significant after including post-mortem interval (PMI) as a covariate in a multiple linear regression model (5.4⋅10^-10^ calls per year; p = 0.002). This finding for liver tissue persisted when analyzing only ssDNA C>T calls (2.5⋅10^-10^ calls per year; p = 0.002) and only ssDNA non-C>T calls (2.9⋅10^-10^ calls per year; p = 0.002) when including PMI as a covariate (**Extended Data Figs. 9a,b**). However, ssDNA call burdens tended to increase with PMI in post-mortem kidney and liver samples (**Extended Data Fig. 9c**), and although this association with PMI was not statistically significant, since other tissues did not exhibit an increase in ssDNA burden with age, it is possible that PMI does not fully capture post-mortem effects that may explain the increase in ssDNA calls with age in liver.

**Figure 5.**
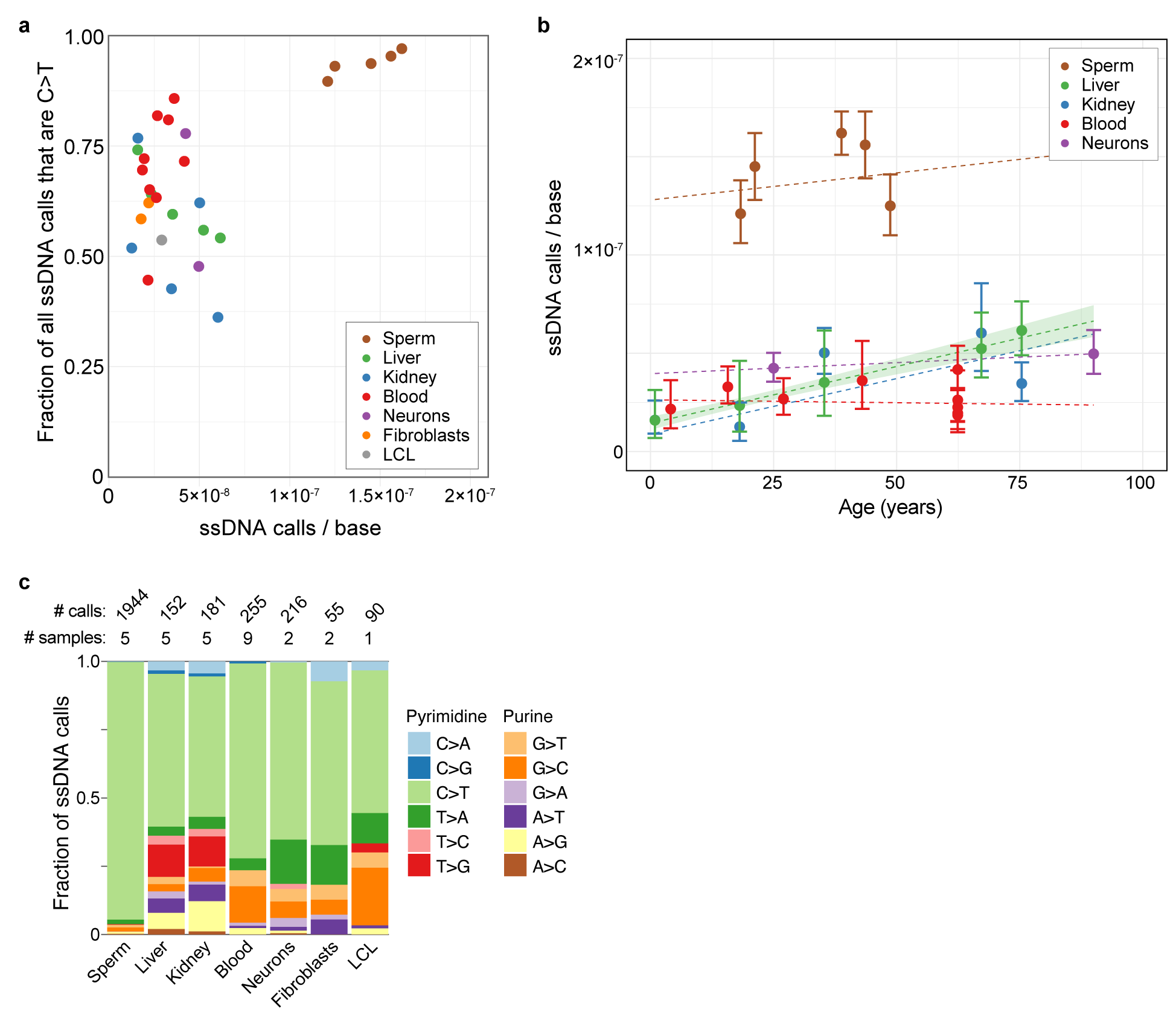
ssDNA call burdens and patterns in samples from healthy individuals. **a,** Fraction of ssDNA calls that are C>T (corrected for trinucleotide context opportunities) across all HiDEF-seq samples from healthy individuals and cell lines (i.e., excluding cancer-predisposition syndromes), versus the total ssDNA call burden. LCL, lymphoblastoid cell line. **b,** ssDNA call burden versus age across all HiDEF-seq v2 samples from healthy individuals (primary tissues only). Dashed lines: weighted least-squares linear regression, with a 95% confidence interval (shaded ribbon) shown for the statistically significant association for liver. **c,** Fraction of ssDNA call burdens by context for samples from healthy individuals and cell lines, after pooling calls separately for each tissue. Call burdens are corrected for trinucleotide context opportunities. See **Extended data Figs. 9d,e** for ssDNA and dsDNA call burdens by context for individual samples, and **Extended data Fig. 9f** for ssDNA spectra for each tissue. **b,** Error bars, Poisson 95% confidence intervals.

Analysis of sequence contexts of ssDNA calls across tissues was notable for the high fraction of C>T calls in sperm and a small increase in the fraction of T>G and A>G calls in post-mortem kidney and liver (**Fig. 5c and Extended Data Figs. 9d,e**). The latter A>G calls may be due to adenine oxidation or deamination damage occurring either pre-or post-mortem, leading the damaged adenine to mispair with cytosine during sequencing^74,75^. ssDNA call spectra of all tissues were similar to SBS30ss* (cosine similarities 0.72-0.93 in non-sperm tissues and 0.997 in sperm) (**Extended Data Fig. 9f**), which may be either due to endogenous or *ex vivo* cytosine deamination. No other ssDNA signature was identified, likely due to the low ssDNA call burdens in healthy tissue samples. Further studies of ssDNA mismatch patterns in healthy tissues, and whether these increase with age in some tissues, will require higher throughput single-molecule sequencing instruments.

### Single- and double-strand DNA calls in the mitochondrial genome

Prior studies have measured an ∼20-40-fold higher somatic dsDNA mutation rate with age in the mitochondrial genome than the nuclear genome^18^. However, the mechanism by which the mitochondrial genome mutates remains unclear^76-81^. While it was long assumed to be due to oxidative damage from mitochondrial oxidative metabolism^78,82^, recent studies have not identified oxidation-related mutational signatures such as G>T mutations from 8-oxoguanine, and instead have found patterns supporting a mechanism closely linked to replication^76-78,80,83,84^. Specifically, A>G and C>T dsDNA mutations are highly enriched on the mitochondrial heavy strand (the G+T-rich ‘-‘ strand of the reference genome, which is the template strand for most genes)—i.e., A>G and C>T changes on the heavy strand with complementary changes on the opposite light strand— with a gradient in frequency that decreases with distance from the origin of replication in the direction of heavy strand synthesis^76,77,80,83^. Several potentially overlapping hypotheses have been proposed for these findings: a) the mitochondrial genome’s strand-displacement mechanism of replication leaves the heavy strand exposed for a longer time as single-stranded DNA, making it vulnerable to deamination of adenine and cytosine that are then mispaired during replication with cytosine and adenine, respectively^75-77,80,85,86^; b) strand asymmetries in polymerase misincorporation of canonical nucleotides^77,78,81^; and, c) strand asymmetries in DNA repair^77^. Importantly, if DNA repair is not substantially more efficient in mitochondria than the nuclear genome^87^, we would expect HiDEF-seq to detect the latter two possibilities as ssDNA events, since HiDEF-seq detects an increased burden of ssDNA mismatches of canonical nucleotides in the nuclear genomes of mismatch repair-deficient and *POLE* PPAP samples that have even lower dsDNA mutation rates than mitochondria: 8.1 and 5.4-fold lower, respectively (**Figs. 6a,b and Extended Data Fig. 7d**). Because HiDEF-seq captures molecules from one third of the mitochondrial genome (**Methods**), we investigated mitochondrial dsDNA and ssDNA call burdens and patterns to distinguish among these hypotheses.

**Figure 6.**
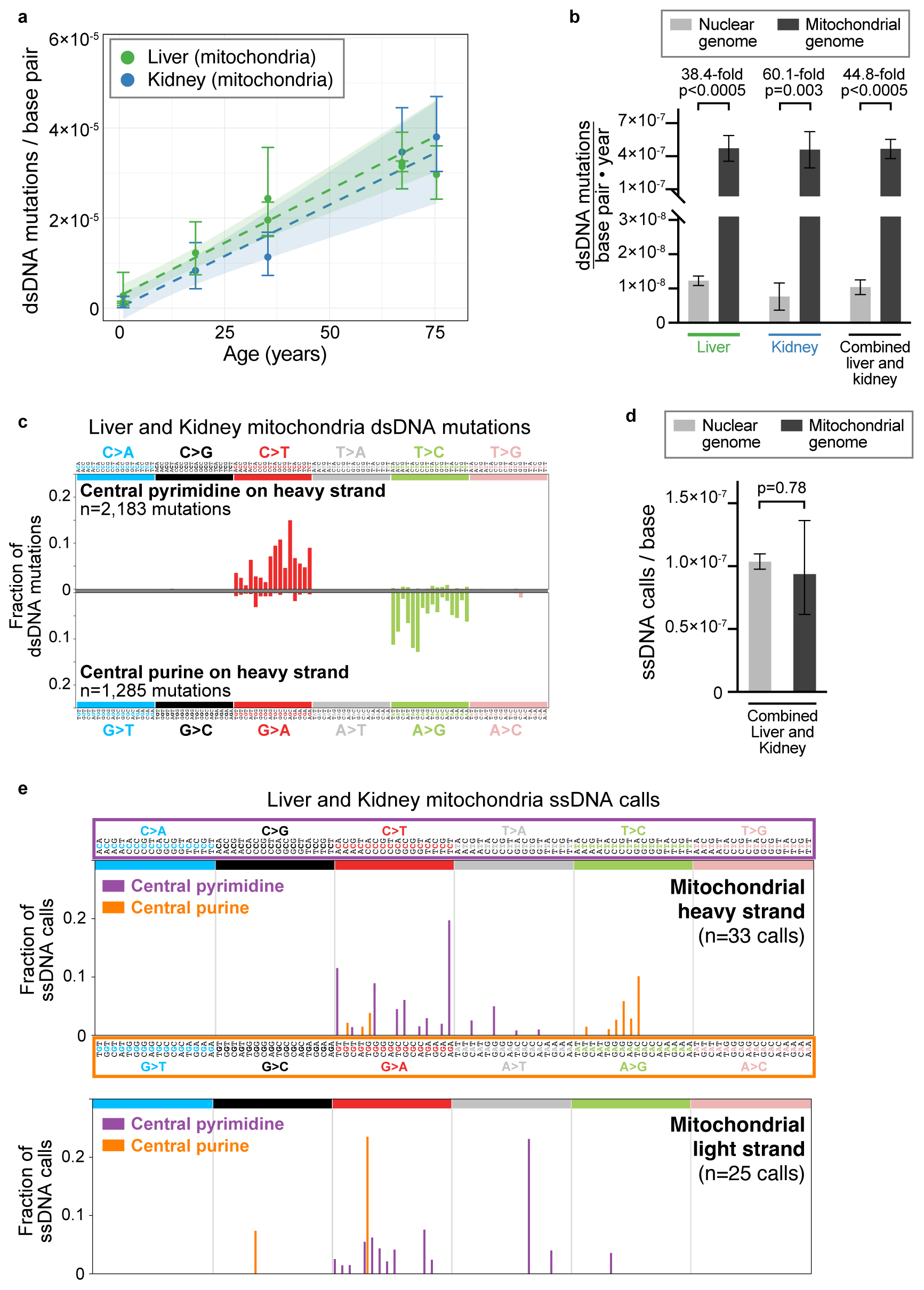
Mitochondrial genome dsDNA and ssDNA call burdens and patterns. **a,** dsDNA mutation burdens versus age in the mitochondrial genome of liver and kidney samples, including liver samples from which mitochondria were enriched. Dashed lines: weighted least-squares linear regression (p < 0.0005 and p = 0.003 for regression slope for liver and kidney, respectively), with a 95% confidence interval (shaded ribbon). **b,** dsDNA mutation burdens per year in the nuclear versus mitochondrial genome. Liver and kidney mitochondrial genome data is from the regressions in panel (a), which were similarly performed for the nuclear genome as well as for liver and kidney samples combined. P-values, comparing the nuclear versus mitochondrial genome within each tissue type, obtained from an ANOVA comparing two weighted least-squares linear regression models of mutation burden versus age and genome type covariates: one with and one without an ‘age x genome type’ interaction term (an estimate of the difference of the dsDNA mutation burden slope versus age depending on whether it is the nuclear or mitochondrial genome). **c,** dsDNA mutation spectra in liver and kidney samples for the mitochondrial genome heavy strand, separated by pyrimidine (top) and purine (bottom) contexts. **d,** ssDNA call burdens in the nuclear versus mitochondrial genomes. Calls are pooled from liver and kidney samples, including liver samples from which mitochondria were enriched (n=1126 and n=27 nuclear and mitochondrial genome calls, respectively). P-value, ANOVA. **e,** Spectrum of ssDNA calls combined from liver and kidney samples, including samples profiled by HiDEF-seq v2 with A-tailing, as well as liver samples from which mitochondria were enriched. **a,d,** Error bars, Poisson 95% confidence intervals. **b,** Error bars, 95% confidence intervals from regressions. **c,e,** Spectra are corrected for trinucleotide context opportunities.

We focused on liver and kidney samples, which had a higher yield of mitochondrial DNA (average 1% of sequenced molecules per sample) than other tissues (**Supplementary Table 1**). Additionally, we purified mitochondria from 3 liver samples, which further increased the yield of mitochondrial DNA (average 13% of sequenced molecules per sample; **Supplementary Table 1**). We detected the expected increase in mitochondrial dsDNA mutation burden with age (**Fig. 6a**), and this mitochondrial genome dsDNA mutation rate was 38.4- and 60.1-fold higher in liver and kidney, respectively, than the dsDNA mutation rate of these tissues’ nuclear genomes (**Fig. 6b**). Combining liver and kidney samples, the difference was 44.8-fold (**Fig. 6b**). HiDEF-seq also detected the expected highly asymmetric pattern of A>G and C>T dsDNA mutations on the heavy strand, though with different distributions of peaks than prior bulk cancer sequencing studies of mosaic mitochondrial mutations^77,83^ (**Fig. 6c**). There was no significant similarity of the full mitochondrial mutation spectrum to COSMIC signatures. However, there was significant similarity between the A>G portion of the heavy strand spectrum and the C>T SBS30ss* cytosine deamination and COSMIC SBS30 signatures (cosine similarities 0.96 and 0.92, respectively) (**Extended Data Fig. 10a**). The mechanism for this similarity is unclear, but this finding suggests a deamination mechanism for A>G heavy strand mitochondrial mutagenesis (37% of dsDNA mutations in our data) and that the same biophysical effects that determine the propensity of cytosine to deaminate preferentially within certain trinucleotide contexts similarly affects adenine deamination.

Notably, despite the large differences in dsDNA mutation rates in the mitochondrial and nuclear genomes, their ssDNA call burdens were not significantly different (p = 0.78, ANOVA) (**Fig. 6d**). Specifically, there were only 27 ssDNA calls in 2.7⋅10^5^ mitochondrial DNA molecules interrogating 3.8⋅10^8^ bases of mitochondrial DNA. While the number of ssDNA calls was low, these were concentrated in sequence contexts consistent with the dsDNA mutation spectrum (**Extended Data Fig. 10b**). To further assess if these ssDNA call patterns are consistent with specific mutagenic mechanisms, we increased the number of analyzed ssDNA calls (n=58) by including liver and kidney samples previously profiled by HiDEF-seq v2 with A-tailing, since the ssDNA T>A artifact that A-tailing can incur is orthogonal to the contexts of mitochondrial mutagenesis. The spectrum of this larger call set was likewise consistent with the dsDNA mutation spectrum and with the following possible mechanisms of mutagenesis: cytosine deamination on the heavy strand (15 / 20 heavy strand central pyrimidine calls are C>T), adenine deamination on the heavy strand (8 / 13 heavy strand central purine calls are A>G), and cytosine deamination on the light strand (18 / 22 light strand central pyrimidine calls are C>T) (**Fig. 6e**). Additionally, a low level of G>A calls on both the heavy and light strands (5 / 13 and 2 / 3 central purine calls are G>A in each strand, respectively) may be due to polymerase G>A misincorporation and/or guanine deamination^88^ (**Fig. 6e**). Altogether, these data strengthen the evidence that the mitochondrial genome mutates primarily during replication via deamination of cytosine and adenine on the heavy strand while it is single-stranded, and to a lesser extent via deamination of cytosine on the light strand.

## Discussion

Mosaic dsDNA mutations reflect the cumulative sum of prior ssDNA mismatches and damage that were not repaired or that were misrepaired^5,20^. Therefore, profiling dsDNA mutations and profiling ssDNA mismatches and damage differ in two important ways. First, profiling dsDNA mutations interrogates past mutational events, while sequencing ssDNA mismatch and damage provides a real-time view of DNA lesions that reflects the current equilibrium between DNA damage, repair, and replication. Second, once ssDNA mismatches and damage transform into dsDNA mutations, information is lost about the originating DNA lesions. These gaps in studying mutagenesis motivated us to develop HiDEF-seq—the first single-molecule DNA sequencing to achieve single-molecule fidelity for single-base changes—which enabled interrogation of dsDNA and ssDNA changes simultaneously. It is also the first ultra-high fidelity long-read sequencing. Our approach opens new avenues for studying DNA damage and mutation processes, which we illustrate by profiling diverse samples, both healthy and with defects in DNA replication and repair.

Mutational signatures have transformed the study of both cancer and mosaic mutations in healthy tissues^8^, but current signatures reflect only dsDNA mutations. We define here the first ssDNA signatures: SBS10ss, SBS14ss, and SBS30ss*. SBS10ss and SBS14ss, the ssDNA signatures of defective polymerase epsilon proofreading with and without functional mismatch repair, respectively, arise from misincorporation of canonical (i.e., non-damaged) nucleotides during replication. It is possible that ssDNA mismatches of canonical nucleotides also occur outside the setting of replication, including post-mitotically. For example, signature SBS5 is ubiquitous in all cells and occurs over time in post-mitotic neurons^16,89^. A recent study indicates SBS5 may be caused by translesion polymerases^90^, implying a mechanism of misincorporation of canonical nucleotides that may become detectable by HiDEF-seq with higher throughput instruments. We anticipate HiDEF-seq will spur future studies to elucidate additional ssDNA mismatch and damage signatures to create a ssDNA signature catalogue that complements the existing dsDNA signature catalogue. It will then be important to relate specific ssDNA and dsDNA signatures to each other, as these relationships will encode information about the dynamics of DNA damage, repair, and replication.

The prevailing view that single-molecule sequencers have low single-molecule fidelity and high cost, with the exception of studies investigating *in vitro* polymerase and bacterial mutagenesis^91-93^, may have deterred their use in studying mosaic mutation processes. Since HiDEF-seq captures data from both DNA strands more efficiently than high fidelity duplex sequencing approaches that utilize short-read (Illumina) sequencing, it is only ∼4-fold more expensive than short-read duplex sequencing, which will reduce to an ∼2-fold difference with the upcoming introduction of new sequencing instruments (**Supplementary Note 2**). Our work also highlights the principle of repeated measurement, in this case increasing the number of sequencing passes of single DNA molecules, to exponentially increase fidelity^24^. Indeed, an analogous approach was recently employed for single-molecule protein sequencing^94^. Further, we achieve single-molecule fidelity for dsDNA mutations with fewer sequencing passes than expected, which supports sperm DNA as an essential, readily available reagent for accurately assessing sequencing fidelity^16,29^. This also implies that with appropriate library preparation and computational filtering, single-molecule fidelity may be feasible in the context of standard PacBio long-read (HiFi) sequencing to enable concurrent detection of germline and mosaic dsDNA mutations, though not ssDNA events that require more sequencing passes.

One limitation of HiDEF-seq is that it does not achieve single-molecule fidelity for insertions and deletions (indels) due to high sequencing error rates for these events in single-molecule sequencing^26^. This may become feasible with improved sequencing fidelity and indel-tuned consensus sequence calling software^26^. Additionally, while we present the first single-molecule fidelity profiling of ssDNA damage and illustrate its effects on sequencing polymerase kinetics, we do not currently detect types of ssDNA damage that do not affect base pairing or that are not compatible with replication by the sequencing polymerase. Since diverse types of ssDNA damage alter sequencing polymerase kinetics^69^, other types of damage, such as 8-oxoguanine, may be feasible to detect in the future with single-molecule fidelity by incorporating kinetics into the initial detection.

We detect an increase in ssDNA call burdens in healthy (i.e., non-tumor) samples of individuals with congenital mismatch repair deficiency and individuals with abnormal polymerase proofreading—syndromes that harbor the highest mosaic mutation burdens found to date in healthy tissues^30,32^. These syndromes’ high mutation rates put their ssDNA events within range of currently feasible HiDEF-seq single-molecule sequencing depth. Likewise, we observe abnormal ssDNA burdens and patterns in tumors deficient in both mismatch repair and polymerase proofreading. However, we did not detect altered ssDNA burdens or patterns in cancer-predisposition syndromes involving nucleotide excision repair or base repair. This may be not only due to the limitation of currently feasible sequencing depth, but also due to mechanisms of mutation involving types of ssDNA damage that we do not detect. We anticipate that future increases in throughput of single-molecule sequencing instruments will enable sufficiently deep profiling to interrogate ssDNA events in cancer-predisposition syndromes with lower mutation rates and in tissues from individuals with normal mutation rates.

A variety of methods can profile individual types of DNA damage, either by enzymatic alteration at damage sites or by affinity enrichment^21,95^. These have revealed important information about sequence context patterns of DNA damage, but since these approaches do not have single-molecule fidelity, their derived damage patterns have low resolution^21,95^. HiDEF-seq can sequence the context of ssDNA cytosine deamination damage with single-molecule fidelity, which reveals its ssDNA signature: SBS30ss*. The correspondence between SBS30ss* and the dsDNA signature SBS30, which is associated with *in vivo* oxidative deamination damage to cytosine^96^, indicates that the SBS30ss* deamination process occurs *in vivo* and that its pattern has a biophysical basis that is independent of *in vivo* processes. It will be illuminating for future studies to attempt to explain this fundamental signature of cytosine deamination damage from first principles (for example, local structural effects on ionization energies) and by molecular dynamics simulations^97^, as this will serve as a ground truth for building predictive models for other DNA oxidation and deamination signatures.

Signatures SBS1 and SBS30/SBS30ss* arise from deamination of 5-methylcytosine and cytosine, respectively^6,96^, and both deamination processes occur at rates within ∼2-fold of each other^98^. Yet, SBS1 dsDNA mutations are detected ubiquitously in all tissues and increase with age, but not SBS30. This discrepancy—even more notable given that most cytosines in the genome are not methylated—is likely explained by differential rates of repair, i.e., the T:G mismatches leading to SBS1 are repaired less efficiently than the uracil-species:G mismatches of SBS30ss*^99,100^. However, why do we detect SBS30ss* but not a ssDNA signature corresponding to SBS1? This suggests that SBS30ss* detected in healthy tissues reflects primarily cytosine deamination that occurs *ex vivo*. Using previously estimated cytosine deamination rates at 37 °C ^98,101^ (2.6 – 7⋅10^-13^ events/second), the ssDNA C>T burdens we detect in post-mortem liver and kidney correspond to spontaneous deamination that would occur over ∼8 – 21 hours without any repair, which is in the range of these samples’ post-mortem intervals plus library preparation times. The detection of SBS30ss* at low levels even in freshly collected blood suggests this may be a difficult residual *ex vivo* burden to avoid. However, in sperm, the significantly greater burden of SBS30ss* may reflect true *in vivo* cytosine deamination damage that accumulates in the absence of effective DNA repair, which is later repaired by the egg post-fertilization^66,67,102,103^.

Our data for the mitochondrial genome is consistent with the asynchronous strand displacement model of mitochondrial genome replication, where mutations occur primarily by deamination of cytosine and adenine on the denatured heavy strand during replication^76,80,85^. However, the pattern of dsDNA C>T mutations in mitochondria does not resemble the cytosine deamination signatures SBS30/SBS30ss*. This may reflect differences in repair efficiency of different sequence contexts in the mitochondrial and nuclear genomes. Alternatively, the low-level signal of G>A ssDNA calls on both the heavy and light strands suggests that polymerase misincorporation and/or guanine deamination may also contribute to dsDNA C>T mutations, such that the mitochondrial dsDNA C>T mutation spectrum may be a combination of these signatures plus a cytosine deamination signature. Overall, our results further constrain the possible mechanisms of mitochondrial mutagenesis.

In addition to profiling primary tissues, HiDEF-seq may find utility in experimental systems to dissect the kinetics of the DNA damage, repair, and replication equilibrium—for example, combined with *in vitro* genetic and other manipulations, with synchronization of the cell cycle, and in reconstituted enzyme systems. It may also be used in biochemical studies of DNA damage. Sequencing single-strand changes to DNA with single-molecule fidelity will transform our understanding of the origins of mutations in cancer and in aging, as well as mutation processes throughout biology.

## Methods

### Sample sources

Post-mortem tissues were obtained from the NIH NeuroBioBank (University of Maryland site). Post-mortem tissues were frozen by the NIH NeuroBioBank in isopentane-liquid nitrogen baths and stored at −80 °C until use. Blood was obtained from individuals enrolled in human subjects research protocols approved by the New York University Grossman School of Medicine Institutional Review Board, the Hospital for Sick Children (SickKids) Research Ethics Board as part of the International Replication Repair Deficiency Consortium (IRRDC) biobank, and the University of Pittsburgh Institutional Review Board. All blood samples were collected in EDTA tubes and frozen immediately after collection until use. Tumor samples were obtained from the IRRDC. Semen samples were obtained at Cryos International Sperm Bank from individuals enrolled in human subjects research approved by the New York University Grossman School of Medicine Institutional Review Board. Lymphoblastoid cell lines were obtained from Coriell Institute and the IRRDC. Primary fibroblasts were obtained from Coriell Institute.

The source, sex, age at collection, and post-mortem interval of each sample are listed in **Supplementary Table 1.**

### Cell culture

Lymphoblastoid cell lines were cultured in T25 flasks with RPMI 1640 media (Thermo Fisher, product #61870036) supplemented with 15% fetal bovine serum and penicillin-streptomycin. Cells were incubated at 37 °C, 5% CO2, and ambient oxygen. Cells were passaged to new media approximately every 2-3 days.

Fibroblasts were cultured in T25 flasks with DMEM media (Thermo Fisher, product #10569010) supplemented with 10% fetal bovine serum and penicillin-streptomycin. Cells were incubated at 37 °C, 5% CO2, and ambient oxygen. Cells were passaged to new media every 3-5 days prior to reaching full confluency. Cells were harvested for DNA at 80-90% confluency using trypsin-EDTA.

### Sperm purification

After collection at Cryos International Sperm Bank, semen underwent liquefaction at room temperature for 30 to 60 minutes. Semen then immediately underwent initial purification for sperm using density gradient centrifugation followed by a wash with HEPES-buffered media^104^. For semen from individuals D1 and D2, sperm were purified from half of each semen sample by this method, and sperm were purified from the other half with a ZyMot Multi (850 µL) Sperm Separation Device (ZyMot) per the manufacturer’s instructions. After addition of cryopreservation media, sperm were stored in liquid nitrogen until further use.

Cryopreserved sperm that previously underwent initial purification by density gradient centrifugation were further purified in the laboratory with a second density gradient centrifugation and two additional washes, as follows. First, the following reagents were equilibrated to room temperature: ORIGIO gradient 40/80 buffer (Cooper Surgical, 84022010), Origio sperm wash buffer (Cooper Surgical, 84050060), and Quinn’s Advantage sperm freezing medium (Cooper Surgical, ART-8022). In a 15 mL tube, 1 mL of Origio 80 buffer was placed at the bottom, and 1 mL of Origio 40 buffer was gently layered on top. Sperm were thawed at room temperature for 15 minutes, gently pipette mixed, and carefully layered on top of the Origio 40 buffer. The tube was then centrifuged in a swinging bucket centrifuge at 400xg for 20 minutes at room temperature with low acceleration and deceleration speeds. The supernatant was aspirated, leaving 500 μL of sperm/buffer at the bottom. The sperm was transferred to a new 15 mL tube and diluted with 5 mL sperm wash buffer. The tube was mixed by inverting 10 times and centrifuged in a swinging bucket centrifuge at 300xg for 10 minutes at room temperature with maximum acceleration and deceleration. The supernatant was removed, leaving about 350 μL of sperm/buffer at the bottom. The sperm was then washed again in the same way with 5 mL of sperm wash buffer, and the supernatant was removed, leaving about 250 μL of sperm/buffer at the bottom of the tube. After pipette mixing, an aliquot of this sperm was transferred to a 2 mL DNA LoBind microtube (Eppendorf) for immediate DNA extraction and general evaluation using a hemocytometer. The remaining sperm was diluted dropwise with a 1:1 volumetric ratio of sperm freezing medium, incubated at room temperature for 3 minutes, frozen in a Mr. Frosty freezing container (Thermo Fisher) at −80 °C freezer for 24 hours, and then transferred to a liquid nitrogen freezer.

### Cerebral cortex neuronal nuclei purification

Cerebral cortex neuronal nuclei were isolated as previously described^9^ from post-mortem frontal cortex of individuals who did not have any known neurological or psychiatric disease: 1) Subject 5344 (Brodmann area 9, left hemisphere) and Subject 6371 (Brodmann area 9, left hemisphere). Specifically, approximately 1 gram of frozen tissue from each was cut into 5 mm^3^ pieces and added to 9 mL of chilled lysis buffer (0.32 M sucrose, 10 mM Tris HCl pH 8, 5 mM CaCl2, 3 mM magnesium acetate, 0.1 mM EDTA, 1 mM DTT, 0.1% Triton-X) in a large dounce homogenizer (Sigma-Aldrich D9938). While on ice, the tissue was dounced 20 times each with pestle size A and then B. The homogenate was layered on a 7.4 mL sucrose cushion (1.8 M sucrose, 10 mM Tris HCl pH 8, 3 mM magnesium acetate, 1 mM DTT) in an ultra-centrifuge tube on ice. Tubes were centrifuged (Thermo Fisher Sorvall LYNX 6000) at 10,000 rpm for 1 hour at 4 °C. The resulting supernatant was removed, and 500 μL of nuclei resuspension buffer (3 mM MgCl2 in 1x Phosphate-Buffered Saline) was added on top of the pellet and then incubated on ice for 10 minutes. The pellet was then gently resuspended. Antibody staining buffer was prepared by adding 1.2 μg of NeuN-Alexa-647 (abcam ab190565) to 400 μL of antibody staining buffer (3% BSA in nuclei resuspension buffer) and inverted gently to mix. 400 μL of antibody staining buffer was added to 1 mL of nuclei and the sample was rotated at 4 °C for 30 minutes. NeuN-positive nuclei were gated as shown in **Supplementary Note 3**. NeuN-positive nuclei were collected in 30 μL of nuclei buffer in 1.5 mL LoBind tubes (Eppendorf) via fluorescence-activated nuclei sorting (FANS) on an LE-SH800 sorter. After sorting, a 1:1 volumetric ratio of 80% glycerol was added to sorted nuclei for a final concentration of 40% glycerol to stabilize nuclei during centrifugation. Nuclei were centrifuged at 4 °C, 500xg for 10 minutes. Supernatant was removed and nuclei pellets were immediately frozen at −80 °C.

### Extraction and isolation of mitochondria

Mitochondria were extracted and isolated from 300 – 500 mg of tissue using the Mitochondria Extraction Kit (Miltenyi Biotec) and Mitochondria Isolation Kit (Miltenyi Biotec), per the manufacturer’s Extraction Kit protocol, with the following modifications: a) protease inhibition buffer was prepared with 100x HALT protease inhibitor cocktail (Thermo Fisher); b) minced tissue was resuspended with a larger 2 x 2.5 mL volume of protease inhibitor buffer instead of 2 x 1 mL; c) after homogenization, the homogenate was passed through a 30 µm SmartStrainer (Miltenyi Biotec); d) the SmartStrainer was washed with 2 x 2.5 mL of solution 3 instead of 2 x 1 mL; e) prior to adding TOM22 antibody, the homogenate was diluted with Separation Buffer to a volume of 25 mL instead of 10 mL; and, f) 125 µL TOM22 antibody was used per sample instead of 50 µL. Final mitochondria pellets were frozen at −20 °C for subsequent DNA extraction.

### DNA extraction

The DNA extraction method used for each sample is listed in **Supplementary Table 1**. Below are details of each DNA extraction method.

#### DNA extraction from sperm for HiDEF-seq

An aliquot of washed sperm (i.e., after the washes that are performed after density gradient centrifugation) was centrifuged at 300xg for 5 minutes at room temperature. The supernatant was removed, leaving approximately 50 μL of sperm/buffer at the bottom of the microtube. The tube was tapped gently 5 times to break up the sperm pellet before adding lysis buffer.

If starting with frozen sperm instead of an aliquot of washed sperm, the frozen sperm vial was rapidly thawed in a 37°C water bath, gently pipette mixed, and an aliquot is transferred to a 2 mL DNA LoBind microtube for DNA extraction. The remaining sperm was frozen again. The DNA extraction aliquot was diluted with 600 μL of Origio sperm wash buffer, centrifuged at 300xg for 5 minutes at room temperature, and the supernatant was removed to leave approximately 100 μL of sperm/buffer at the bottom. The sperm was diluted again with 600 μL of Origio sperm wash buffer, centrifuged at 300xg for 5 minutes at room temperature, and the supernatant was removed to leave approximately 50 μL of sperm/buffer at the bottom. The tube was tapped gently 5 times to break up the sperm pellet before adding lysis buffer.

Sperm DNA extraction was based on a prior study^105^, with some modifications, including optimizations we performed that showed that TCEP (tris(2-carboxyethyl)phosphine) can be reduced from 50 mM to 2.5 mM in the lysis buffer. Specifically, sperm lysis buffer was prepared by combining (for each sample) 497.5 μL of Qiagen Buffer RLT (Qiagen) without beta-mercaptoethanol, and 2.5 μL of 0.5 M Bond-Breaker TCEP Solution (Thermo Scientific) for a lysis buffer with 2.5 mM TCEP final concentration. 500 μL of sperm lysis buffer and 100 mg of 0.2 mm stainless steel beads (Next Advance, SSB02-RNA) were then added without mixing to each sample. Homogenization was then performed with a TissueLyser II instrument (Qiagen) at 20 Hz for 4 minutes (samples SPM-1002, SPM-1020, SPM-1013 HiDEF-seq v2 with A-tailing, and SPM-1004) or 30 seconds (samples SPM-1060, SPM-1013 HiDEF-seq v2 without A-tailing, D1, and D2). DNA was then extracted from the lysate using the QIAamp DNA Mini Kit (Qiagen) with a modified protocol as follows. 500 μL of buffer AL was added to each lysate and vortexed well. Then, 500 μL of 100% ethanol was added and vortexed well. Then, the mixture was applied to a QIAamp DNA Mini spin column and the remaining standard QIAamp protocol was followed. DNA was eluted with 100 μL of 10 mM Tris pH 8. RNase treatment was then performed by adding 12 μL of 10x PBS pH 7.4 (Gibco), 2 μL of Monarch RNase A (New England Biolabs (NEB)), and 6 μL nuclease-free water (NFW). The reaction was incubated at room temperature for 5 minutes and immediately purified using a 0.8X beads to sample volume ratio of SPRI beads (solid-phase reversible immobilization; made by washing 1 mL Sera-Mag Carboxylate-Modified SpeedBead [Cytiva, #65152105050250] and resuspending the beads in 50 mL of 18% PEG-8000, 1.75 M NaCl, 10 mM Tris pH 8, 1 mM EDTA, 0.044% Tween-20). DNA was eluted from beads with 35 μL of 10 mM Tris/0.1 mM EDTA pH 8.

A somatic cell contamination assay was adapted from a prior study^106^ and performed on all extracted sperm DNA samples to further confirm sperm purity. This assay amplifies 4 loci from bisulfite-treated DNA: 3 loci that are methylated in sperm but not in somatic cells (PCR7, PCR11, PCR31), and 1 locus that is methylated in somatic cells but not in sperm (PCR12). Following bisulfite treatment and PCR amplification of each locus, the PCR amplicon is only cut by a restriction enzyme if the original DNA was methylated. Therefore, this assay can detect somatic cell contamination. 350 ng of each extracted sperm DNA and 350 ng of control human NA12878 lymphoblastoid cell line genomic DNA (Coriell Institute) were bisulfite-converted using the Zymo EZ DNA Methylation kit (Zymo Research). The loci were amplified by PCR using the following primer sets: PCR7 (GGGTTATATGATAGTTTATAGGGTTATT and TCTATTACTACCACTTCCTAAATCAA), PCR11 (TGAGATGTTTGTTAGTTTATTATTTTGG and TCATCTTCTCCCACCAAATTTC), PCR12 (TAGAGGGTAGTTTTTAAGAGGG and ATTAACCAACCTCTTCCATATTCTT), and PCR31 (TTTTAGTTTTGGGAGGGGTTGTTT and CTACCAAAATTAAAAACCAACCCAC). The PCR reaction contained 1.5 μL of bisulfite-converted DNA, 10 μL of 2X ZymoTaq PCR Mix (Zymo Research), PCR primers, and NFW to a final volume of 20 μL. The PCR reactions were optimized to contain the following final concentrations of each forward and reverse primer: 0.6 μM for PCR7 primers, 0.6 μM for PCR11 primers, 0.3 μM for PCR12 primers, and 0.45 μM for PCR31 primers. The PCR reactions were cycled at: 95 °C for 10 minutes, (94 °C for 30 sec; X °C for 30 sec; 72 °C for 30 sec) x 40 cycles, 72°C for 7 minutes, 4 °C hold, where X (annealing temperature) was 49 °C for PCR7 and PCR11, 51 °C for PCR12, and 55 °C for PCR31. PCR reactions were purified by 2X volumetric ratio SPRI beads cleanup and eluted in 22 μL of 10 mM Tris pH 8. Restriction digests were performed by combining 5 μL of purified PCR product, restriction enzyme (10 units of HpyCH4IV [NEB] for PCR7 and PCR31, and 20 units of TaqI-v2 [NEB] for PCR11 and PCR12), 1 μL of 10X CutSmart buffer (NEB), and NFW for a total reaction volume of 10 μL. Restriction digestions were performed at 37 °C (HpyCH4IV) or 65 °C (TaqI-v2) for 60 minutes. Control reactions without restriction enzyme were performed for each sample/locus combination. 5 μL of each restriction digest reaction was combined with 1 μL 6X TriTrack DNA Loading Dye (Thermo Fisher) and run on a 2% agarose gel pre-stained with ethidium bromide, followed by imaging of the gel.

#### DNA extraction from solid tissues for HiDEF-seq

Approximately 50 – 300 mg of tissue was cut in a petri dish on dry ice and minced with a scalpel, followed by one of the following DNA extraction methods, as specified for each sample in **Supplementary Table 1**.

##### ‘Nucleobond HMW, MagAttract HMW, QIAamp’

In this method, DNA was extracted and purified with 3 serial kits to maximize DNA purity. DNA was extracted using the NucleoBond HMW DNA kit (Takara) per the manufacturer’s instructions with a 50 °C proteinase K incubation for 4.5 hours. The eluted DNA was then further purified with the MagAttract HMW DNA kit (Qiagen) per the manufacturer’s whole blood purification protocol, except with Proteinase K / RNase A incubation occurring at 56 °C for 20 minutes. The eluted DNA was then further purified using the QIAamp DNA mini kit (Qiagen) by diluting the DNA to a final volume of 200 μL and final 1x PBS concentration, adding 20 μL of Proteinase K (Qiagen), and continuing per the manufacturer’s body fluids DNA purification protocol with a 56 °C proteinase K incubation for 10 minutes without RNase A treatment.

##### ‘MagAttract HMW’

We used the MagAttract HMW DNA (Qiagen) per the manufacturer’s protocol for tissue, with a 2-hour proteinase K digestion at 56 °C. DNA was eluted with 10 mM Tris pH 8.

##### ‘Puregene’

Tissue was pulverized inside a microtube while in a liquid-nitrogen cooled mini mortar and pestle (Bel-Art). DNA was then extracted with the Puregene DNA Kit (Qiagen) per the manufacturer’s protocol for tissues, except: 1) the lysis step was performed at room temperature on a ThermoMixer C instrument (Eppendorf) at 1,400 rpm for 1 hour; 2) the RNase A treatment was performed at room temperature for 20 minutes, and; 3) the final DNA pellet was resuspended in 10 mM Tris pH 8 at room temperature for 1 hour.

#### DNA extraction from cerebral cortex neuronal nuclei for HiDEF-seq

DNA was extracted from nuclei pellets by two methods, as specified for each sample in **Supplementary Table 1**.

##### ‘QIAamp’

We used the QIAamp DNA Mini kit (Qiagen) per the manufacturer’s protocol, with lysis performed by adding 180 μL of Buffer ATL and 20 μL of proteinase K to the nuclei pellet, followed by a 56 °C incubation for 1 hour, and including RNase A treatment.

##### ‘MagAttract’

We used the MagAttract HMW DNA kit per the manufacturer’s protocol for blood, after resuspending nuclei with 200 µL of 1x PBS, with a 30-minute proteinase K digestion at room temperature.

#### DNA extraction from mitochondria for HiDEF-seq

DNA was extracted from mitochondria pellets with the Puregene DNA Kit (Qiagen) per the manufacturer’s protocol for tissues, except: a) the lysis step used 200 µL Cell Lysis Solution and 1.5 µL Proteinase K and was performed at room temperature for 30 minutes; and, b) the RNase A treatment was performed at room temperature for 20 minutes.

Note, due to relatively low yields of mitochondria DNA preparations, these samples were profiled with HiDEF-seq v2 with A-tailing (see ‘HiDEF-seq library preparation’).

#### DNA extraction from blood, lymphoblastoid cells, and fibroblasts for HiDEF-seq and germline sequencing

DNA from blood, lymphoblastoid cells, and fibroblasts (the latter two after resuspending cell pellets in 1x PBS)—except for blood from individuals whose tumors were profiled—was extracted with the MagAttract HMW DNA kit (Qiagen) per the manufacturer’s whole blood purification protocol, with proteinase K incubation at room temperature for 30 minutes. We also performed an experiment that excluded a measurable cytosine deamination effect by possible leached iron from MagAttract magnetic beads (**Extended Data Fig. 8d**), by extracting an additional aliquot of DNA from the blood of individual 1901 with the Puregene DNA Kit (Qiagen) per the manufacturer’s protocol for ‘whole blood or bone marrow’, except: 1) 200 µL blood was first diluted with 100 µL of 1x PBS; b) the cell lysis step was performed at room temperature; and, c) the RNase A treatment was performed at room temperature for 20 minutes.

#### DNA extraction from tumors and those individuals’ corresponding blood for Illumina tumor and germline sequencing

DNA was extracted from tumors by first homogenizing the tumor with the Precellys 24 Tissue Homogenizer followed by the DNeasy Blood & Tissue Kit (Qiagen), per the manufacturer’s protocol for animal tissues with a 56 °C incubation for 10 minutes. For individuals whose tumors were profiled, DNA was extracted from blood of those individuals with the PAXgene Blood DNA Kit (Qiagen) per the manufacturer’s protocol.

#### DNA extraction from saliva for Illumina germline sequencing

DNA was extracted with the QIAamp DNA Mini Kit per the manufacturer’s ‘DNA purification from blood or body fluids’ protocol and including RNase A treatment.

#### DNA extraction from liver and spleen for Illumina germline sequencing

DNA was extracted with the QIAamp DNA Mini Kit per the manufacturer’s ‘DNA purification from tissues’ protocol, with a 2-hour proteinase K digestion at 56 °C and including RNase A treatment.

#### DNA extraction from blood for Pacific Biosciences germline sequencing

DNA was extracted using the Chemagic DNA Blood 2k Kit (Perkin Elmer CMG-1097) on a Chemagic 360 automated nucleic extraction instrument (Perkin Elmer) following manufacturer’s protocols for DNA isolation from whole blood.

#### DNA quantity and quality measurements, and storage

Concentration and quality of all DNA samples were measured using a NanoDrop instrument (Thermo Fisher), a Qubit 1X dsDNA HS Assay Kit (Thermo Fisher), and a Genomic DNA ScreenTape TapeStation Assay (Agilent). DNA was stored at −20 °C.

### Illumina germline and tumor library preparation and sequencing

Illumina germline and tumor sequencing libraries were prepared using the TruSeq DNA PCR-Free kit (Illumina) for all samples, except GM10430 that used the TruSeq DNA Nano kit (Illumina). At least 110 Gb (∼36x genome coverage) of 150-base pair paired-end sequencing per sample was obtained using a Novaseq 6000 instrument (Illumina) by Psomagen Inc, except for tumor sequencing and those individuals’ corresponding germline sequencing that used HiSeqX and Novaseq 6000 instruments at the Centre for Applied Genomics at the Hospital for Sick Children.

### Pacific Biosciences germline library preparation and sequencing

15 μg of DNA was cleaned with a 1X AMPure PB beads (Pacific Biosciences) cleanup and sheared to a target size of 14 kilobases (kb) using the Megaruptor 3 instrument (Diagenode) with the following settings: Speed 36, volume 300 µL, Conc. 33 ng/µL. Library preparation was performed with the SMRTbell Express Template Prep Kit 2.0 (Pacific Biosciences) per the manufacturer’s instructions. Library fragments longer than 10 kb were selected using a PippinHT instrument (Sage Science). Size-selected libraries were sequenced on a Pacific Biosciences Sequel IIe system using the Sequel II Binding Kit 2.0 and Sequel II Sequencing Kit 2.0 (Pacific Biosciences), Sequencing primer v4 (Pacific Biosciences), 2-hour binding time, adaptive loading, 2-hour pre-extension, and 30-hour movies.

### Heat damage of DNA

DNA was heated in a volume of 62 µL at the temperature and time, and in the buffer, listed for each sample in **Supplementary Table 1**, followed by incubation on ice up to a total of 6 hours if the heating time was less than 6 hours. Untreated samples were incubated on ice for 6 hours.

The DNA was then input into HiDEF-seq library preparation.

### NanoSeq library preparation and sequencing

NanoSeq libraries were prepared as previously described^16^ with 50 ng DNA input from the same DNA aliquots used for HiDEF-seq.

### HiDEF-seq library preparation and sequencing

#### Choice of restriction enzymes for DNA fragmentation

We performed *in silico* digests of the CHM13 v1.0 human reference genome sequence^107^ to identify restriction enzymes that: a) maximize the percentage of the genome between 1 – 4 kb, b) are active at 37 °C, and, c) the DNA is fragmented with blunt ends, since blunt fragmentation avoids single-strand overhangs that can lead to artifactual double-strand mutations during end repair^16^. This *in silico* screen identified Hpy166II (recognition sequence: 5’-GTN/NAC-3’) as the optimal restriction enzyme, with a prediction of 37% of the genome mass fragmenting between 1 – 4 kb. The percentage of the genome fragmented to sizes between 1 – 4 kb was then empirically measured by fragmenting 1 µg of genomic DNA followed by quantification on a Genomic DNA ScreenTape assay (Agilent). Hpy166II fragments 41% of the genome to within the target size range. Note that although Hpy166II is blocked by methylated CpG when present on both sides of the recognition sequence (New England BioLabs), this will occur only with the larger recognition sequence 5’-C*GTN/NAC*G-3’ (‘*’ signifies methylation of preceding cytosine); excluding all these potential bi-methylated sites increases the *in silico* predicted percentage of the genome fragmented by Hpy166II to within the target size range by 0.2%, and 99.97% of genomic bases within the original target size range remain when excluding these as potential fragmentation sites.

For the mitochondrial genome, Hpy166II captures 3 fragments in our target 1 – 4 kb size range, at the following coordinates (CHM13 v1.0): 1) 3068 – 5116 (2048 base pairs, bp), 2) 7581 – 9439 (1858 bp), and, 3) 10441 – 11831 (1390 bp). These fragments encompass 32% of the mitochondrial genome.

#### HiDEF-seq library preparation

Input DNA amounts of 500 – 3000 ng (as measured by the Qubit 1X dsDNA HS Assay [Qubit]) were used per library, depending on available DNA. With high-quality DNA and HiDEF-seq v2 (i.e., with nick ligation, see below), input amounts of 500 ng provide sufficient library yield for approximately one full (non-multiplexed) Pacific Biosciences (PacBio) Sequel II instrument sequencing run, and lower input amounts are feasible for filling a fraction of a sequencing run. We have successfully made HiDEF-seq libraries with as low as 200 ng input DNA, producing sufficient yield for 40% of a sequencing run. For fragmented DNA samples, more than 1500 ng of input DNA is generally required. Generally, for samples other than sperm and tissues from young children that have low mutation burdens, one quarter of a sequencing run is sufficient for mutation burden and pattern analysis. Input DNA A260/A280 > 1.8 and A260/A230 > 2.0 absorption ratios were confirmed on NanoDrop prior to library preparation per the Pacific Biosciences DNA preparation guidelines; we find this quality control to be important for sequencing yield for post-mortem tissues, but not strictly necessary for other sample types.

Since some DNA fragments are < 1 kb after restriction enzyme fragmentation, these small fragments need to be removed during library preparation. We found that effective removal of < 1 kb DNA fragments with high-yield recovery of larger DNA fragments requires three size selections with a 75% dilution of AMPure PB beads (Pacific Biosciences) during library preparation. We also found that efficient size selection critically depends on a DNA concentration < 10 ng/µl in the input sample. Accordingly, before beginning library preparation, a sufficient volume of AMPure PB Beads was diluted with Elution Buffer (Pacific Biosciences) to a final 75% AMPure PB bead volume/total volume solution to be used for all subsequent bead purifications and size selections. Below, ‘diluted AMPure beads’ refers to these diluted beads.

Input DNA was fragmented with 0.14 U/µL of Hpy166II restriction enzyme (NEB) in a 70 µl reaction with 1x CutSmart buffer (NEB) for 20 minutes at 37 °C. The fragmentation reaction was scaled to a higher volume if the input DNA was too dilute to accommodate a 70 µL reaction.

Next, the fragmentation reaction was diluted with NFW to a DNA concentration of 10 ng/µl (or not diluted if DNA is already < 10 ng/µl) based on the Qubit concentration of the DNA measured prior to the fragmentation reaction. For the first bead purification/size selection, a ratio of 0.8X diluted AMPure beads volume to sample volume was used, with two 80% ethanol washes, and the DNA was eluted with 22 µL of 10 mM Tris pH 8. The DNA concentration was measured again with Qubit.

For HiDEF-seq v2 (but not HiDEF-seq v1), nick ligation was then performed in a 30 µL reaction with 3 µL of 10x rCutSmart Buffer (NEB), 1.56 µL of 500 µM β-Nicotinamide adenine dinucleotide (NAD+) (NEB), and 15U of E. coli DNA Ligase (NEB). The nick ligation reaction was incubated at 16° C for 30 minutes with the heated lid turned off.

The DNA was then diluted with 10 mM Tris pH 8 to 10 ng/µL (or not diluted if DNA is already < 10 ng/µl) based on the Qubit concentration measured after the post-fragmentation reaction bead purification. For the second bead purification/size selection, a ratio of 0.75X diluted AMPure bead volume to sample volume was used, with two 80% ethanol washes, and the DNA was eluted with 22 µL of 10 mM Tris pH 8. DNA concentration was measured with Qubit.

The DNA was then treated as in ref. ^16^ in 30 µL volume reactions with 21 µL of input DNA, 1.5 µL NFW, 3 µL 10x NEBuffer 4 (NEB), 3 µL of 1 mM each dATP/ddCTP/ddGTP/ddTTP (dATP/ddBTP) (dATP, Thermo Fisher R0141; ddCTP/ddGTP/ddTTP, Jena Bioscience NU-1019S), and 7.5 U of Klenow fragment 3’→5’ exo-(NEB), except for input DNA that is degraded (e.g., post-mortem kidney and liver) for which the reaction was performed without dATP. The reaction was incubated at 37° C for 30 minutes. Next, a third bead purification/size selection was performed: the reaction volume was diluted with 10 mM Tris pH 8 to 10 ng/µL DNA (or not diluted if DNA is already < 10 ng/µl) based on the Qubit concentration measured prior to the Klenow reaction, followed by a ratio of 0.75X diluted AMPure bead volume to sample volume, with two 80% ethanol washes, and elution of DNA with 22 µl of 10 mM Tris pH 8. The eluted DNA was then adjusted to a total of 30 µL with 3 µL of 10X NEBuffer 4 and NFW before proceeding to adapter ligation.

Ligation of hairpin adapters was performed using reagents from the SMRTbell Express Template Prep Kit 2.0 (Pacific Biosciences) by combining 30 μL of Klenow-treated DNA, 2.5 µL Barcoded Overhang Adapter (Pacific Biosciences), 15 µL Ligation Mix, 0.5 µL Ligation Additive, and 0.5 µL of Ligation Enhancer. For samples whose preceding Klenow reaction was performed without dATP, 2.5 µL of 17 µM annealed blunt adapters were used instead (see sequences and preparation below). The adapter ligation reaction was incubated at 20° C for 60 minutes with the heated lid turned off. Immediately after the adapter ligation, nuclease treatment was performed using the SMRTbell Enzyme Clean Up Kit 1.0 (Pacific Biosciences) to remove any non-circularized DNA containing nicks and/or without hairpin adapters: 2 µL Enzyme A, 0.5 µL Enzyme B, 0.5 µL Enzyme C, and 1 µL Enzyme D were combined, and this 4 µL enzyme mix was added to the ligation reaction and incubated at 37° C for 60 minutes. After the nuclease treatment, samples were purified with a ratio of 1.2X diluted AMPure bead volume to sample volume and eluted with 24 µL of 10 mM Tris pH 8.

After the post-nuclease treatment AMPure bead purification, non-A tailed HiDEF-seq v2 libraries underwent an additional 1.1X diluted AMPure bead purification to remove residual adapter dimers.

Final library concentration and size distribution were measured with Qubit and High Sensitivity D5000 ScreenTape (Agilent). The final library fragment size distribution should contain < 5% of DNA mass < 1 kb and < 5% of DNA mass > 4 kb (percentages calculated using the ScreenTape analysis software’s manual region analysis ‘% of Total’ field). Final mass yield of A-tailed libraries should be ∼6-10% of the input genomic DNA mass, and approximately half of that for non-A tailed libraries. Libraries were stored in 0.5 mL DNA LoBind tubes at −20 °C.

On ScreenTape, some non-A tailed HiDEF-seq v2 libraries had a low level of residual adapter dimers, which was removed with a final 1.3X diluted AMPure bead purification after multiplexing the libraries from the same run (see multiplexing details in the ‘HiDEF-seq library sequencing’ section).

#### Sequences and preparation of blunt adapters used for HiDEF-seq without A-tailing

Adapters (sequences below) were ordered as HPLC-purified oligonucleotides from Integrated DNA Technologies. Each adapter was reconstituted to 100 µM concentration with NFW. Annealing was then performed for each adapter at a concentration of 17 µM in a 30 µL volume containing 10 mM Tris pH 8 and 50 mM NaCl, by incubating at 95 °C for 3 minutes and cooling at room temperate for 30 minutes. Additional barcoded adapters can be designed by replacing the below barcodes with alternative sequences.

bcAd1001:

/5’Phos/ACGCACTCTGATATGTGATCTCTCTCTTTTCCTCCTCCTCCGTTGTTGTTGTTGAGAGAGATCACATATCAGAGTGCGT (barcode = CACATATCAGAGTGCG)

bcAd1002:

/5’Phos/ACTCACAGTCTGTGTGTATCTCTCTCTTTTCCTCCTCCTCCGTTGTTGTTGTTGAGAGAGATACACACAGACTGTGAGT (barcode = ACACACAGACTGTGAG)

bcAd1003:

/5’Phos/ACTCTCACGAGATGTGTATCTCTCTCTTTTCCTCCTCCTCCGTTGTTGTTGTTGAGAGAGATACACATCTCGTGAGAGT (barcode = ACACATCTCGTGAGAG)

bcAd1008:

/5’Phos/ACGCAGCGCTCGACTGTATCTCTCTCTTTTCCTCCTCCTCCGTTGTTGTTGTTGAGAGAGATACAGTCGAGCGCTGCGT (barcode = ACAGTCGAGCGCTGCG)

#### Modified HiDEF-seq library preparation trials to remove single-strand DNA T>A artifacts

Below are details of trial to remove single-strand DNA (ssDNA) T>A artifacts arising from ssDNA nicks that remain after nick ligation. The final protocol that completely removes these artifacts (HiDEF-seq v2 without A-tailing) is described in the main ‘HiDEF-seq library preparation’ section.

##### Polynucleotide kinase

The standard HiDEF-seq v2 protocol was followed with the exception that prior to nick ligation, the DNA was treated in a 30 µL reaction containing 12 U T4 polynucleotide kinase (NEB), 1 mM ATP (NEB), 4 mM DTT (Promega), and 1x CutSmart buffer (NEB) at 37 °C for 1 hour. The sample then proceeded into the nick ligation reaction in a higher reaction volume of 35 µL, with reaction components scaled proportionally to the higher volume and a final 1x CutSmart buffer concentration.

##### Alternative A-tailing polymerases

The standard HiDEF-seq v2 protocol was followed with the exception of replacing Klenow fragment 3’→5’ exo-polymerase with one of the following: 9.6 U Bst large fragment (NEB), 9.6 U Bst 2.0 (NEB), 9.6 U Bst 3.0 (NEB), or 9 U Isopol SD+ (ArcticZymes). Reaction temperatures and times for these polymerases were: a) Bst large fragment and Bst 2.0: 30 minutes at 45 °C; b) Bst 3.0: either 30 minutes or 150 minutes at 45 °C, or 210 minutes at 37 °C; and, c) Isopol SD+: either 30 minutes or 210 minutes at 37 °C.

##### Pyrophosphatase

The standard HiDEF-seq v2 protocol was followed with the exception of adding 0.15 U of E. coli inorganic pyrophosphatase (NEB).

##### Klenow reaction without dATP or without dATP/ddBTP

The standard HiDEF-seq v2 protocol was followed with the exception that the Klenow reaction was performed without dATP or without dATP/ddBTP.

##### No Klenow reaction

The standard HiDEF-seq v2 protocol was followed, except after the post-nick ligation bead purification, the DNA was diluted to 30 µL in a final 1x NEBuffer 4 concentration and taken directly to adapter ligation using blunt adapters. Following the post-nuclease treatment bead purification, an additional size selection was performed with 0.75X diluted AMPure beads since this would ordinarily have occurred after the Klenow reaction. Note: this protocol produces a CCT>CGT ssDNA artifact that does not occur when the Klenow reaction is performed without dATP or ddBTP, indicating that Klenow polymerase removes this artifact likely via a pyrophosphorolysis mechanism (**Extended Data Fig. 5d and Supplementary Table 3**).

#### HiDEF-seq large fragment library preparation

Large fragment size libraries (1-10 kb range; median 4.1 kb size) were prepared per the above HiDEF-seq library protocol, except: 1) fragmentation was performed with 30 U PvuII-HF enzyme (NEB) instead of Hpy166II, 2) post nick ligation and post A-tailing cleanups were performed with 1.8X undiluted AMPure PB beads, and DNA was not diluted to < 10 ng/µL (since size selection is not being performed), and, 3) final post-nuclease treatment bead purification was performed with 1X undiluted AMPure PB beads.

#### HiDEF-seq library sequencing

Libraries sequenced on the same sequencing run were multiplexed together based on the final library Qubit quantification, to achieve at least 50 ng of total library in no more than 15 µL volume. When necessary, the concentration of individual or pooled libraries can be increased by room temperature centrifugal vacuum concentration (Eppendorf Vacufuge) and pausing periodically (approximately every 2 minutes) to avoid increases in temperature, or with an AMPure PB bead purification.

Sequencing was performed on Pacific Biosciences Sequel II or Sequel IIe systems with 8M SMRT Cells by the Icahn School of Medicine at Mount Sinai Genomics Core Facility and the New York University Grossman School of Medicine Genome Technology Center. Sequencing parameters were: Sequel II Binding Kit 2.0 (Pacific Biosciences), Sequel II Sequencing Kit 2.0 (Pacific Biosciences), Sequencing primer v4 (Pacific Biosciences), 1-hour binding time, diffusion loading, loading concentrations between 125–160 pM (lower concentration was used for blood than for tissues) for standard size libraries (Hpy166II libraries) or 80 pM for large fragment libraries (PvuII libraries), 2-hour pre-extension, and 30-hour movies.

### Illumina germline and tumor sequencing data processing

#### Illumina germline and tumor sequencing data processing

Reads were aligned to the CHM13 v1.0 reference genome^107^ with BWA-MEM v0.7.17^108^ with standard settings, followed by marking of optical duplicates and sorting using the Picard Toolkit^109^. Variants were called from the aligned reads with two different variant callers: 1) Genome Analysis Toolkit (GATK) ^110^ v4.1.9.0 (v, version) using the HaplotypeCaller tool with parameters ‘-ERC GVCF -G StandardAnnotation -G StandardHCAnnotation -G AS_StandardAnnotation’ followed by the GenotypeGVCFs tool with default parameters; 2) DeepVariant^111^ v1.2.0 with parameter: ‘--model_type=WGS’. Both GATK and DeepVariant variant calls were used during subsequent analysis.

#### Pacific Biosciences germline sequencing data processing

Circular consensus sequences were derived from raw subreads (a subread is one sequencing pass of a single strand of a DNA molecule) using pbccs v5.0.0 (ccs, Pacific Biosciences) with default parameters, and consensus sequences were filtered to only retain high-quality ‘HiFi’ reads; i.e., reads with predicted consensus sequence accuracy ‘rq’ tag ≥ 0.99 (‘rq’ is calculated by ccs as the average of the per base consensus qualities of the read). These reads were then aligned to the CHM13 v1.0 reference genome with pbmm2 v1.4.0 (Pacific Biosciences) with the parameters ‘--preset CCS --sort’. Variants were called from the aligned reads with DeepVariant^111^ v1.2.0 with the parameter: ‘--model_type=PACBIO’.

### HiDEF-seq primary data processing

HiDEF-seq data first undergoes a two-part primary data processing pipeline that transforms the raw data into a format suitable for subsequent analysis. Primary data processing also produces quality control plots generated by custom scripts and by SMRT Link (Pacific Biosciences) software (e.g. distributions of polymerase read lengths, number of passes, etc.). Note, for simplicity, we use the term ‘call’ to refer to both double-strand DNA (dsDNA) mutations and single-strand DNA (ssDNA) mismatch and damage events. The pipeline analyzes calls in sequencing reads that are single base mismatches relative to the reference genome (i.e., not insertions and deletions).

The first part of the primary data processing pipeline utilizes a combination of bash and awk scripts to process raw subread sequencing data (a subread is one sequencing pass of a single strand of a DNA molecule) into a strand-specific aligned BAM format^112^ with additional tags needed for call analysis^112^. The steps of this first part of data processing are as follows:

1. Subreads for which an adapter was not detected on both ends of the molecule (‘cx’ tag not equal to 3) are removed.
2. Consensus sequences are created separately for each strand of the DNA molecule (i.e., forward and reverse strand separately) using pbccs v6.2.0 (Pacific Biosciences) with parameters: --by-strand, --min-rq 0.99 (minimum predicted consensus sequence accuracy > Q20 [Phred quality score] to remove low quality consensus sequences), and --top-passes 0 (unlimited number of full-length subreads used per consensus).
3. Demultiplexing of samples according to adapter barcodes using lima v2.5.0 (Pacific Biosciences) with parameters: --ccs --same --split-named --min-score 80 --min-end-score 50 -- min-ref-span 0.75 --min-scoring-regions 2.
4. Filter to remove any DNA molecules (also known as zero-mode waveguides, ZMWs, which are sequencing wells containing a single DNA molecule) that did not successfully produce both one forward and one reverse strand consensus sequence.
5. Align forward and reverse strand consensus sequences to the CHM13 v1.0 reference genome^107^ using pbmm2 v1.7.0 (Pacific Biosciences), an aligner based on minimap2^113^, with parameters: --preset CCS. Note that the CHM13 v1.0 reference genome only contains nuclear chromosomes 1-22, chromosome X, and the mitochondrial genome—but not chromosome Y, which is therefore not part of the analyses.
6. Filter to keep only DNA molecules that produce only 1 forward strand primary not-supplementary alignment and 1 reverse strand primary not-supplementary alignment, where the forward and reverse alignments overlap (reciprocally) in the genome by at least 90%.
7. Sort alignments by reference position.
8. Add five tags, detailed below, to each alignment in the final BAM file, with each tag containing a comma-separated array with a length corresponding to the number of single base mismatches in the alignment (relative to the reference genome) per the alignment CIGAR string:
  *qp:* Positions of bases in the read sequence (query) that are mismatches relative to the reference genome; 1-based coordinates with the left-most base in the alignment record’s SEQ column = 1.
  *qn:* Sequences of bases in the read (query) that are mismatches relative to the reference genome (base sequences are according to the forward genomic strand, i.e., they are taken from the SEQ column of the SAM alignment record).
  *qq:* Qualities of bases in the read that are mismatches relative to the reference genome (taken from the QUAL column of the SAM alignment record).
  *rp:* Positions in reference genome coordinates of read bases that are mismatches relative to the reference genome.
  *rn:* Sequences of the reference genome at positions of read bases that are mismatches relative to the reference genome.

The second part of the primary data processing pipeline is an R^114^ script (R v4.1.2, requiring the packages Rsamtools^115^, GenomicAlignments^116^, GenomicRanges^116^, vcfR^117^, plyr^118^, configr^119^, qs^120^) that further processes and annotates the aligned BAM file into an R data file, as follows:

1. Load the aligned BAM file into R, including the custom tags that annotated the positions of base mismatches relative to the reference genome.
2. Annotate calls (bases mismatched relative to the reference genome) for which the reference genome base is ‘N’, to exclude these from subsequent analysis.
3. Annotate the positions of insertions and deletions (indels) in each alignment, based on the alignment CIGAR string.
4. Annotate each call if it was present in any of the VCF variant call files of the corresponding individual’s germline sequencing, along with details of the VCF variant annotation.
5. Save positions of indels from the VCF variant call files of the corresponding individual’s germline sequencing.
6. Transform the dataset so that forward and reverse strand consensus reads and ssDNA and dsDNA calls (and tag information) from the same DNA molecule are linked to each other as dsDNA molecules.
7. Save the final R dataset to a file.

### HiDEF-seq call filtering

The call filtering pipeline implements a series of filters that were optimized to maximize the number of true calls while minimizing the number of sequenced bases and regions of the genome that are filtered out. During development of the pipeline, filters and filter parameters were iteratively optimized using low mutation rate samples (i.e., tissues from infants and sperm) to identify patterns common to false positives. These false positives were apparent as clusters of mutations in low-quality regions of the genome and as regions with low-quality alignment of sequencing reads. For example, when a metric of low-quality genome regions was found to correlate with clusters of low-quality calls, this metric was added as a filter, and its threshold was iteratively tuned to maximally remove false positives while minimizing the number of sequenced bases and genomic regions that are filtered.

Additional optimization of filter thresholds was performed using sperm samples that have a known low mutations burden. Specifically, we plotted the dsDNA and ssDNA burdens with a range of thresholds for 3 key filters: 1) minimum predicted consensus accuracy (0.99 to 0.999), 2) minimum number of passes per strand (5 to 20), and, 3) minimum fraction of subreads (passes) detecting the mutation (0.5 to 0.8) (**Extended Data Figs. 3c-j**). We examined these plots for threshold settings above which burden estimates are stable. Since burdens were corrected for sensitivity (based on total interrogated bases and detection of known germline variants; see ‘HiDEF-seq calculation of call burdens’), a decrease in burden estimates with increasing threshold settings indicates removal of sequencing artifacts. These plots showed that sperm dsDNA mutation burden estimates were stable even down to the lowest (most lenient) thresholds (**Extended Data Figs. 3d,e,g**). In contrast, ssDNA burdens required higher threshold settings before burden estimates stabilized (**Extended Data Figs. 3i,j**). Individually increasing the thresholds of each of the above three filters stabilized ssDNA burden estimates at approximately 20%, 15%, and 10% lower levels, respectively, compared to the least stringent settings, and applying all three filters together with these higher thresholds reduced the ssDNA burden estimate by approximately 25% (i.e., the three filters are not independent). Specific thresholds used for dsDNA and ssDNA mismatch filtering are detailed in the below sections detailing each filter.

The call filtering pipeline utilizes the following R packages: GenomicAlignments^116^, GenomicRanges^116^, vcfR^117^, Rsamtools^115^, plyr^118^, configr^119^, MutationalPatterns^>121^, magrittr^122^, readr^123^, dplyr^124^, plyranges^125^, stringr^126^, rtracklayer^127^, qs^120^; and the following software tools: bcftools^128^, samtools^128^, wigToBigWig^129^, wiggletools^130^, pbmm2 (Pacific Biosciences), zmwfilter (Pacific Biosciences), digest^131^, SeqKit^132^, and KMC^133^.

Additional filters used in the pipeline were created using REAPR v1.0.18^134^. REAPR was originally designed to identify regions with errors in reference genome assemblies, but we found that it calculates metrics useful for identifying regions of the genome prone to generating false positive and false negative variant calls in Illumina (short-read) sequencing data. First, Illumina whole-genome sequencing reads from a sperm sample were aligned to CHM13 v1.0 using SMALT v0.7.6^135^ with parameters ‘-r 0 -x -y 0.5’ and a CHM13 v1.0 index created with SMALT using parameters ‘-k 13 -s 2’. Next, reads were sorted and duplicates were marked. The REAPR perfectfrombam command was then run on the resulting BAM file using the parameters ‘min insert=266, max insert=998, repetitive max qual=3, perfect min qual=4, and perfect min alignment score=151’ (min and max insert size are the 1 and 99 percentiles of insert sizes calculated from the sequencing data using the Picard Toolkit CollectInsertSizeMetrics tool). REAPR metrics for each base of the genome were obtained from the output stats.per_base file and a bigwig^136^ annotation file was created for each metric.

The mutation analysis filters were applied serially as described below. Unless otherwise specified, the filters were applied to both ssDNA and dsDNA calls. Note, the computational pipeline has the capability to implement additional filters not listed here, as specified in the pipeline configuration documentation available online.

#### Filters based on DNA molecule quality and alignment metrics

Keep only DNA molecules that meet all the below criteria:

1. ccs predicted consensus accuracy ≥ 0.99 in both forward and reverse strand (i.e., rq tag of ccs ≥ 0.99) for dsDNA calls, and ≥ Q30 (i.e., rq ≥ 0.999) for ssDNA calls.
2. Minimum of 5 (for dsDNA calls) and 20 (for ssDNA calls) sequencing passes for each of the forward and reverse strands (using the ‘ec’ BAM file tag, which is computed by ccs as the average subread coverage across all consensus calling windows).
3. Both forward and reverse strands have mapping quality ≥ 60.
4. Maximum difference in number of ssDNA calls between the forward and reverse strands of 5, before germline variant filtering. This removes artifacts from rare chimeric molecules and residual low-quality molecules.
5. Average of the number of indels relative to the human reference genome in the forward and reverse strands ≤ 20, before germline variant filtering. This removes low-quality molecules with many indels.
6. Average of the number of soft-clipped bases in the forward and reverse strands ≤ 30. This removes low-quality molecules and molecules that align to complex regions of the genome with long stretches of mismatched bases.

#### Filters based on germline sequencing variant calls

1. Filter out calls that were also identified in any of the individual’s germline sequencing VCF files with read depth ≥ 3, allele quality (QUAL column in VCF) ≥ 3, genotype quality (GQ tag in VCF) ≥ 3, and variant allele fraction ≥ 0.05.
2. Filter out DNA molecules with > 8 dsDNA calls remaining after VCF germline filtering. This removes molecules with misalignment to complex regions of the genome leading to many clustered calls and regions of the genome for which Illumina short reads are not effective in identifying and filtering out germline variants.

For tumor analysis, variant calls were used in this step from both germline blood sequencing and standard fidelity (Illumina) tumor sequencing to focus the analysis on low-level mosaic calls.

#### Filters based on genomic regions

A. Filters that remove the entire DNA molecule if it meets any of the below criteria:

1. For analyses using either Illumina or PacBio germline sequencing data:

i. Segmental duplication regions: any overlap with the DNA molecule’s forward or reverse consensus sequence alignments. This annotation was obtained from the file chm13.draft_v1.0_plus38Y.SDs.bed created by the Telomere-to-Telomere consortium^137^. However, for analysis of mitochondrial mutations, this region filter is not used because it contains the region chrM:10000-14910 due to a similar nuclear genome sequence on chromosome 5, which would cause unnecessary filtering of reads aligning to this region of the mitochondrial genome. There is negligible risk of nuclear genome sequences falsely aligning to this mitochondrial region since we obtain long reads, we require high mapping quality, and we exclude reads with many mismatches—and these mitochondrial and nuclear genome regions have only 94% identity.
ii. Satellite sequence regions: ≥ 20% of the DNA molecule’s forward and reverse strand consensus alignments (average for the two strands) overlaps the region. The satellite sequence region annotation was created for CHM13 v1.0 using RepeatMasker v4.1.1^138^ with parameters ‘-pa 4 -e rmblast -species human -html -gff -nolow’, followed by extraction of ‘Satellite’ regions.
2. Only for analyses that use Illumina germline sequencing data, because short-read data is more prone to missing true germline variants in these regions:

i. Telomere regions: any overlap with the DNA molecule’s forward or reverse consensus sequence alignments. This annotation was obtained from the file chm13.draft_v1.0.telomere created by the Telomere-to-Telomere consortium^107^.
ii. 50-mer mappability score: ≥ 30% of the DNA molecule’s forward and reverse strand consensus alignments (average for the two strands) has a mappability score < 0.4. This annotation was created for CHM13 v1.0 using Umap v1.2.0^139^. This annotation calculates mappability for every base in the genome as [the number of *k*-mers overlapping the base that are uniquely mappable to the genome] / *k*.
iii. Fraction of Illumina short reads aligning to the region that are orphaned reads (i.e., the read’s mate is either unmapped or mapped to a different chromosome), averaged across the genome in 20 base pair non-overlapping bins, is ≥ 0.15 for ≥ 20% of the DNA molecule’s forward and reverse strand consensus alignments (average for the two strands). The fraction of orphaned reads metric used in this filter is the average of the orphan_cov and orphan_cov_r REAPR metrics, which are the fraction of forward and reverse strand reads that are orphaned, respectively.
B. Filters that remove only the portions of the DNA molecule that overlap any of the following regions, while the remaining bases of the DNA molecule are still included in analysis:

1. Regions of the reference genome whose sequence is ‘N’.
2. For analyses using either Illumina or PacBio germline sequencing data:

i. Satellite sequence regions: any base that overlaps one of these regions.
ii. Bases with gnomAD v3.1.2^140^ single nucleotide variants with ‘PASS’ flag and population allele frequency > 0.1%, lifted over from the hg38 to the CHM13 v1.0 reference genome using the liftOver tool^129^. This filter removes 27,476,828 genome bases from the analysis. It is required in order to remove residual germline variants that were not detected in the germline sequencing of the individual, and it reduces the risk of false positive mosaic event calls due to very low level contamination that may occur between samples of different individuals^16^.
3. Only for analyses that use Illumina germline sequencing data, because short-read data is more prone to missing true germline variants in these regions:

i. 100-mer mappability score: any base with a mappability score < 0.95, with mappability scores averaged across the genome in 20 base pair non-overlapping bins (binning smoothes the mappability score signal). The primary mappability scores were calculated as described for the above 50-mer mappability score.
ii. Fraction of Illumina short reads aligning to the region that are properly paired (i.e., aligned in the correct orientation and within the expected distance based on insert size distribution), averaged across the genome in 20 base pair non-overlapping bins, is < 0.7. The fraction of properly paired reads metric used in this filter is the average of the prop_cov and prop_cov_r REAPR metrics, which are the fraction of forward strand and reverse strand reads that are properly paired, respectively.
iii. Fraction of Illumina short reads aligning to the region that are orphaned reads (i.e., the read’s mate is either unmapped or mapped to a different chromosome), averaged across the genome in 20 base pair non-overlapping bins, is ≥ 0.2. The fraction of orphaned reads metric used in this filter is the average of the orphan_cov and orphan_cov_r REAPR metrics, which are the fraction of forward and reverse strand reads that are orphaned, respectively.
iv. The number of Illumina short reads aligning to the region to either the forward or the reverse strand and that are soft-clipped at the left end or the right end (i.e., sum of REAPR clip_fl, clip_fr, clip_rl, clip_rr metrics), divided by [4 x number of mapped reads / 100,000,000], averaged across the genome in 200 base pair non-overlapping bins, is ≥ 0.09.
v. The number of Illumina short reads with mapping quality 0 aligning to the region, divided by [4 x number of mapped reads / 100,000,000], averaged across the genome in 20 base pair non-overlapping bins, is ≥ 0.1. Note, this general filtering annotation was calculated using Illumina whole-genome sequencing data of one representative sample.

#### Base quality filter

Filter out dsDNA calls whose consensus sequence base quality is < 93 (from QUAL column in BAM file) in either the forward or reverse strand consensus, and filter ssDNA calls whose base quality is < 93 in the strand containing the call.

#### Filter based on location within the read

Filter out calls that are ≤ 10 bases from the ends of the consensus sequence alignment (alignment span excludes soft-clipped bases). For ssDNA calls, this filter applied to the strand containing the call, and for dsDNA calls, this filter is applied to both the forward and reverse strand consensus sequence alignments. Although this only negligibly alters call burdens (**Extended Data Fig. 3h**), it removes rare alignment artifacts.

#### Filter based on location near germline indels

Regions near germline indels are prone to alignment artifacts that can lead to false positive calls. This filter removes calls located near an indel within a distance less than or equal to twice the length of the indel or less than or equal to 15 bases of the indel (whichever range is larger), using indels called in any of the germline sequencing data of the individual (i.e., both GATK and DeepVariant indel calls when using Illumina germline sequencing data, and only DeepVariant indel calls when using PacBio germline sequencing data). For GATK indel calls, only indels with read depth ≥ 5, QUAL ≥ 10, genotype quality ≥ 5, and variant allele fraction ≥ 0.2 were used in this filtering. For DeepVariant indel calls, only indels with read depth ≥ 3, QUAL ≥ 3, genotype quality ≥ 3, and variant allele fraction ≥ 0.1 were used in this filtering.

#### Filter based on location near consensus sequence indels

Regions near HiDEF-seq consensus sequence indels are prone to alignment artifacts that can lead to false positive calls. This filter removes calls located near a consensus sequence indel within a distance less than or equal to twice the length of the indel or less than or equal to 15 bases of the indel (whichever range is larger). For dsDNA calls, the call must pass this filter on both forward and reverse consensus strands. For ssDNA calls, this filter applies only to the strand containing the call.

#### Filters based on germline sequencing read depth and variant allele fraction

1) Filter out calls in locations where the germline sequencing data has < 15 total reads coverage, since these low-coverage germline sequencing regions will be prone to false-negative germline variant calls that would then lead to false-positive HiDEF-seq calls.

2) Filter out calls detected with variant allele fraction > 0.05 or read depth > 3 in the germline sequencing data to remove variants that were not called by the prior germline variant callers (due to low variant allele fraction or due to different local haplotype assembly in GATK/DeepVariant that calls variants in a different nearby location than the bwa alignment of the consensus molecule sequence). This filter is less stringent than a recent somatic mutation analysis method^16^, but may still remove a small number of very early developmental mosaic variants shared between HiDEF-seq data and the individual’s germline sequencing.

The above two filters use the samtools mpileup command to determine total read depth and variant allele fraction, using the parameters ‘-I -A -B -Q 11 --ff 1024 -d 10000 -a “INFO/AD”’ for Illumina germline sequencing data and the parameters ‘-I -B -Q5 --ff 2048 --max-BQ 50 -F0.1 - o25 -e1 --delta-BQ 10 -M399999 -d 10000 -a “INFO/AD”’ for PacBio germline sequencing data.

For tumor analysis, this filter step used both germline blood sequencing and standard fidelity (Illumina) tumor sequencing to focus the analysis on low-level mosaic calls.

#### Filters based on fraction of subreads (passes) detecting the call and fraction of subreads overlapping the call

We filter out calls detected in < 50% (for dsDNA calls) and < 60% (for ssDNA calls) of the subreads of the DNA molecule that detected the call. For dsDNA calls, this filter is applied to forward and reverse subreads separately, and the call must pass the filter in both strands. For ssDNA calls, this filter is applied only to subreads of the strand in which the call was detected. This remove false positive calls in the consensus sequence that are not well-supported by the subreads. The filter is implemented by first extracting the subreads of all the DNA molecules containing calls from the raw subreads BAM file using the zmwfilter tool (Pacific Biosciences) and aligning them to the CHM13 v1.0 reference genome with pbmm2 v1.7.0 with parameters ‘-– preset SUBREAD --sort’. The bcftools mpileup command is then used with parameters ‘-I -A -B – Q 0 -d 10000 -a “INFO/AD”’ to calculate the fraction of subreads detecting the call (excluding subreads with the supplementary alignment SAM flag).

In rare DNA molecules, a large fraction of subreads are soft-clipped, leading to false positive calls in the small fraction of remaining subreads aligned to the soft-clipped region. We therefore also filter out calls for which the percentage of subreads overlapping the call (regardless of whether they contain the call) out of the total subreads aligned to the genome is < 50%, calculated separately for subreads of each strand for the molecule in which the call was made. This filter is applied to the strand containing the call for ssDNA calls, and to both strands for dsDNA calls (i.e., a dsDNA call must pass this filter in both strands).

### HiDEF-seq calculation of call burdens

Following application of all the above filters, DNA molecules are further filtered to keep only those with a maximum of 1 dsDNA call for dsDNA call burden calculations, and a maximum of 1 ssDNA call per strand for ssDNA call burden calculations. This removes a small number of remaining DNA molecules that contain multiple post-filtering calls that upon manual inspection are due to residual regions of the genome prone to false positives.

The raw dsDNA mutation burden (i.e., mutations per base pair) of a sample is then calculated as the [# of dsDNA calls] / [# of interrogated dsDNA base pairs], and the raw ssDNA call burden (i.e., calls per base) is calculated as the [# of forward strand calls + # of reverse strand calls] / [# of interrogated forward strand read bases + # of interrogated reverse strand read bases]. Note, we subsequently use the term ‘interrogated bases’ for simplicity, even though for dsDNA mutation analysis it refers to interrogated base pairs. The number of interrogated bases takes into account all of the relevant filters that were applied, both filters that remove entire DNA molecules and filters that remove only portions of DNA molecules. Specifically, the number of interrogated bases is the total number of bases of DNA molecules that passed all the filters that remove full DNA molecules (i.e., ‘Filters based on DNA molecule quality and alignment metrics’ and ‘Filters based on genomic regions – section A’), excluding the bases of those remaining DNA molecules removed by the filters that only remove portions of DNA molecules: a) ‘Filters based on genomic regions – section B’, b) ‘Base quality filter’, c) ‘Filter based on location within the read’, d) ‘Filter based on location near germline indels’, e) ‘Filter based on location near consensus sequence indels’, and, f) ‘Filter based on germline sequencing read depth— minimum germline sequencing total read coverage’.

We also calculated ‘corrected’ call burdens that correct both for: 1) differences in trinucleotide sequence context of the genome relative to interrogated bases, and, 2) sensitivity of detection. These corrections were applied as follows:

First, we corrected raw call counts for the trinucleotide frequency distribution of the genome (specifically, the CHM13 v1.0 sequences of chromosomes being analyzed; i.e., chromosomes 1-22 and X for nuclear genome analysis, and the mitochondrial sequence for mitochondrial genome analysis) relative to the trinucleotide frequency distribution of interrogated bases in sequencing reads. This correction for ‘trinucleotide context opportunities’ is necessary because interrogated bases may have a different distribution of trinucleotides than the genome due to restriction enzyme fragmentation and computational filters, and this may affect burden estimates^16^. Specifically, we first calculate the distribution of trinucleotides (fraction of each trinucleotide out of all trinucleotides) across the genome. We then calculate the distribution of trinucleotides across interrogated bases of sequencing reads in the sample. Next, for each trinucleotide, we calculate the ratio of its fractional distribution in the full genome to its fractional distribution in the interrogated bases. The trinucleotide-corrected count of HiDEF-seq calls is then obtained by multiplying the raw call count for each trinucleotide context by that context’s genome/interrogated bases trinucleotide ratio. For dsDNA calls, trinucleotide context corrections are performed using all possible 32 trinucleotide contexts where the middle base is a pyrimidine. For ssDNA calls, trinucleotide context corrections are performed using all 64 possible trinucleotides and using strand-specific trinucleotide sequences of calls, interrogated bases, and the genome. The trinucleotide contexts of ssDNA calls reflect the original DNA molecule’s ssDNA change—i.e., for calls in strands aligning to the forward strand of the reference genome, the reverse complements of the call, interrogated read sequences, and genome are used for trinucleotide context corrections, and vice versa for calls in strands aligning to the reverse strand. This is because the sequence data produced by the sequencer has the directionality of the sequencer-synthesized strand rather than the original (template) DNA molecule.

Second, we corrected call counts for sensitivity of detection separately for each sample using a set of high-quality, true-positive heterozygous germline (dsDNA) variants detected in the HiDEF-seq data of the sample. This specifically accounts for single-molecule sensitivity loss due to the ‘Filters based on fraction of subreads (passes) detecting the call and fraction of subreads overlapping the call’ that are applied to calls detected in the final interrogated bases (they are applied to each strand separately, and dsDNA calls must pass the filters in both strands). All other filters remove DNA molecules and bases from the final set of interrogated bases and therefore do not require sensitivity correction. To generate the true-positive set of heterozygous germline variants for each sample, we extracted all the autosomal dsDNA calls detected in the final interrogated HiDEF-seq bases of the sample that were also called in all the germline variant call sets of the individual with ≥ 50^th^ percentile VCF QUAL score, ≥ 50^th^ percentile VCF genotype quality, ≥ 50^th^ percentile total read depth, and variant allele fraction between 30% to 70%. We keep only calls that meet these criteria across every one of the variant call sets of the individual and that are present in gnomAD v3.1.2 with ‘PASS’ flag and population allele frequency > 0.1%. If more than 10,000 such true-positive germline calls are identified, a random subset of 10,000 calls is selected for the sensitivity calculation. We then extract subreads corresponding to the DNA molecules that detected these calls in the sample, realign them to the genome with pbmm2 v1.7.0 with ‘--preset SUBREAD --sort’ settings, and annotate the variants using the same process described in the ‘Filters based on fraction of subreads (passes) detecting the call and fraction of subreads overlapping the call’ step of the call filtering pipeline. Next, we calculate germline variant sensitivity for the sample as the number of true-positive germline variant calls that pass the same filtering thresholds used in the ‘Filters based on fraction of subreads (passes) detecting the call and fraction of subreads overlapping the call’ step of the call filtering pipeline, divided by the total number of true-positive germline variant calls. Each sample’s dsDNA call counts are then corrected for sensitivity by dividing by that sample’s calculated germline variant sensitivity. Each sample’s ssDNA call counts are corrected by dividing by the square root of that sample’s germline variant sensitivity, because the above dsDNA germline variant sensitivity estimate corrects for filters applied to both strands separately.

Finally, ssDNA and dsDNA burdens corrected for both trinucleotide context and sensitivity are calculated as the sum of the trinucleotide context-and sensitivity-corrected call counts divided by the number of interrogated bases (ssDNA burdens) or base pairs (dsDNA burdens). For all analyses and figures, unless otherwise specified, we use burden estimates corrected for both the full genome trinucleotide distribution and sensitivity.

The Poisson 95% confidence intervals of a sample’s corrected burden were calculated as the corrected burden x [Poisson 95% confidence interval of raw call counts, calculated by the poisson.test function in R] / [raw call counts]. Weighted least-squares linear regressions of call burdens versus age were performed with the ‘lm’ function in R (via the ggplot^141^ package), with weights equal to 1/[raw call counts].

### HiDEF-seq estimate of fidelity for dsDNA mutations

The fidelity for dsDNA mutations was estimated for each sample as follows: a) for each of the 192 possible trinucleotide contexts (i.e., both central pyrimidine and central purine contexts), the number of single-strand mismatches at that context was divided by the total number of interrogated bases to obtain a ssDNA call burden for that context; b) for each central pyrimidine trinucleotide context, a dsDNA mutation error probability was calculated by multiplying the single-strand call burdens of the corresponding central pyrimidine and reverse complement central purine trinucleotide contexts; and, c) all of the resulting central pyrimidine trinucleotide context dsDNA mutation error probabilities were summed.

### Comparison of HiDEF-seq and standard PacBio HiFi molecule characteristics

Standard PacBio HiFi raw subreads data for comparison to HiDEF-seq (**Fig. 1b and Extended Data Figs. 1d,f**) were obtained from the Human Pangenome Reference Consortium (HPRC) public data repository^142^ (samples HG02080, HG03098, HG02055, HG03492, HG02109, HG01442, HG02145, HG02004, HG01496, HG02083). Circular consensus sequences were derived from raw subreads using the same ccs version and ccs parameters used to analyze HiDEF-seq data.

### Comparison of HiDEF-seq mutation burdens in sperm to paternally-phased de novo mutation burdens

Paternally-phased de novo mutation (DNM) burdens were calculated for each paternal age (in 1-year intervals) from data published in a prior study of 2,976 trios^28^ (aau1043_datas5_revision1.tsv and aau1043_datas7.tsv supplementary files), and using additional methodological details obtained from its associated study by Jonsson, et al^143^. Paternally-phased DNM burdens were first calculated for each child as [total number of DNMs] x [fraction of phased DNMs across the full cohort that are paternally phased] x [Jonsson et al’s correction factor of 1.009 that accounts for its false positive and negative rate] / [Jonsson et al’s interrogated genome size of 2,682,890,000]^28,143^. We then compare the dsDNA mutation burden of each HiDEF-seq sperm sample to the DNM burdens of children whose fathers’ age at their birth is 1-year higher than the age at which the sperm sample was collected (to account for ∼ 9 months difference between the father’s age at conception and the child’s birth).

### Comparison of HiDEF-seq and NanoSeq call burdens and patterns

NanosSeq data was processed using the NanoSeq analysis pipeline available at https://github.com/cancerit/NanoSeq. NanoSeq dsDNA burdens corrected for trinucleotide context opportunities were obtained from the ‘results.mut_burden.tsv’ output file of the NanoSeq pipeline. NanoSeq ssDNA call burdens were calculated as the sum of the values in the ‘results.mismatches.subst_asym.tsv’ output file, divided by this number plus double the number of interrogated dsDNA base pairs obtained from the ‘results.mut_burden.tsv’ output file. NanoSeq ssDNA call counts for each context were obtained from the ‘results.SSC-mismatches-Pyrimidine.triprofiles.tsv’ and ‘results.SSC-mismatches-Purine.triprofiles.tsv’ output files. Since the NanoSeq pipeline does not correct ssDNA calls for trinucleotide context opportunities, we compared the burdens of NanoSeq ssDNA calls in each context to the burdens of HiDEF-seq ssDNA calls that are also not corrected for trinucleotide context opportunities (i.e., to HiDEF-seq burdens corrected only for sensitivity).

For more informative comparison of Poisson 95% confidence intervals of HiDEF-seq and NanoSeq (**Figs. 1c,e,f and Extended Data Fig. 4a**) despite a different number of interrogated bases (for ssDNA calls, or base pairs for dsDNA calls) measured by each method, for each sample the number of calls of the method with the higher number of interrogated bases (or base pairs) was downsampled proportionally to the ratio of the number of interrogated bases of the two methods. The downsampled method’s burden was then recalculated as the downsampled call count divided the number of interrogated bases of the method with fewer interrogated bases, and the downsampled method’s Poisson 95% confidence interval was recalculated using the downsampled number of raw call counts. This downsampling does not affect burden estimates, and it normalizes the two methods’ confidence intervals to reflect an equivalent number of interrogated bases (or base pairs). Confidence intervals prior to downsampling are in **Supplementary Table 2**.

### Signature analysis

Signature analysis for dsDNA mutations was performed using the ‘sigfit’ package^144^, with input of raw mutation counts for each trinucleotide context, and the ‘opportunities’ parameter set to the ratio of the fractional abundance of each trinucleotide context in interrogated bases of that sample versus the fractional abundance of that trinucleotide context in the human reference genome. The correction for trinucleotide context opportunities performed above for burden analyses used the fractional abundance of trinucleotides in CHM13 v1.0, but the correction for trinucleotide context opportunities performed here for signature analysis and figures used the fractional abundance of trinucleotides in the full GRCh37 genome (for both nuclear and mitochondrial genome analyses and figures) so that the obtained spectra and signatures can be compared to standard COSMIC signatures. The ‘plot_gof’ function was used to determine the optimal number of signatures to extract. Since COSMIC SBS1 was not well separated from other signatures during de novo extraction^145^, we utilized the ‘fit_extract_signatures’ function to fit SBS1 while simultaneously extracting additional signatures de novo. De novo extracted signatures were compared to the COSMIC SBS v3.2 catalogue^35^ to identify the most similar known signature by cosine similarity. To obtain more accurate estimates of signature exposures, the fitted COSMIC SBS signature and the extracted signatures were then re-fit back to the mutation counts using ‘fit_signatures’ function, along with correction for trinucleotide context opportunities. SBS5 is a ubiquitous clock-like signature^35^, and often de novo extraction produced more than one signature highly similar to SBS5, for example, both SBS5 and SBS3 (cosine similarity 0.79) or both SBS5 and SBS40 (cosine similarity 0.83) or both SBS3 and SBS40 (cosine similarity 0.88). In these cases, we either reduced the number of de novo extracted signatures so that only one of these similar signatures was extracted, or we instructed ‘fit_extract_signatures’ to fit both COSMIC SBS1 and COSMIC SBS5.

ssDNA signatures were extracted by taking advantage of sigfit’s capability to analyze 192-trinucleotide context mutational spectra that distinguish transcribed versus untranscribed strands. Instead, we use this feature to distinguish central pyrimidine versus central purine contexts. We do this by arbitrarily setting central pyrimidine and central purine ssDNA calls to the transcribed and untranscribed strands, respectively (by setting the strand column to ‘-1’ for all calls that are input into sigfit’s ‘build_catalogues’ function, without collapsing central pyrimidine and central purine contexts). We then extract ssDNA signatures as described above for dsDNA signatures, with correction for trinucleotide context opportunities. Cosine similarities of ssDNA and dsDNA signatures are calculated after projecting ssDNA signatures to 96-central pyrimidine contexts, which is performed by summing values of central pyrimidine contexts with values of their reverse complement central purine contexts.

### Replication strand asymmetry (fork polarity) analysis

ENCODE replication timing (Repli-seq) data^146^ (wavelet-smoothed signal) was obtained from the UCSC Genome Browser^129^ (hg19) for the lymphoblastoid cell lines GM12878, GM06990, GM12801, GM12812, and GM12813. We calculated the average of the Repli-seq signal (higher values indicate earlier replication) across these samples at each position, and then lifted over the data to CHM13 v1.0. For each analyzed HiDEF-seq call, we calculated fork polarity^147^ as the slope versus position of the Repli-seq data points spanning -5 to +5 kilobases from the call using the ‘lm’ function in R. Positive and negative fork polarities indicate the genome non-reference (-) strand is synthesized more frequently in the leading and lagging strand direction, respectively. This was also performed for a set of 50 iterations of 1,000 randomly selected genomic positions with either the sequence or the reverse complement of the sequence corresponding to the trinucleotide context being analyzed (i.e., AGA or TCT for *POLE* samples). We then calculated the fork polarity quantile values at quantiles ranging from 0 to 1.0 in 0.1 increments, and then for each of these quantile bins (combining 0.4-0.5 and 0.5-0.6 quantile bins into one bin, as these span fork polarity 0) we counted the number of loci whose sequence is AGA in the genome non-reference (-) strand and the number of loci whose sequence is AGA in the reference genome (+) strand. Loci without annotated Repli-seq data were excluded. Next, for each genome strand, we calculated normalized call counts by dividing the quantile bin call counts by the total number of calls in that strand. For each of the 9 quantile bins, we then calculated the ‘strand ratio’ as the ratio of non-reference to reference strand normalized call counts. We also calculated this strand ratio for positive and negative fork polarities (i.e., two bins rather than 9 quantile bins), since there were not enough ssDNA calls in individual quantile bins for analysis. Analyses were also repeated after excluding loci within genic regions annotated in the CHM13 v1.0 LiftOff Genes V2 annotation obtained from the UCSC Genome Browser.

### Kinetics analysis

For each sample, consensus sequences for each strand were created with pbccs v6.4.0 (Pacific Biosciences) with parameters: --by-strand --hifi-kinetics --min-rq 0.99 --top-passes 0. pbccs v6.4.0 was used because with these parameters it outputs consensus kinetics values for each strand separately, which prior versions of pbccs do not. Consensus sequence reads were then aligned to the CHM13 v1.0 reference genome with pbmm2 with the parameters ‘--preset CCS -- sort’.

Next, we extracted the list of ssDNA C>T sequence calls in the heat-treated blood DNA and sperm samples (sequenced by HiDEF-seq v2). Due to the very high number of ssDNA C>T calls in blood DNA samples that were heat-treated in water-only or Tris-only buffer, for these samples we selected a random subset of 800 calls. We then extracted from these samples and from 88 other HiDEF-seq samples all the consensus reads that overlapped the C>T call positions, from the strand synthesized by the sequencing polymerase opposite to the strand on which the call is present in the molecule. Since kinetics is affected by sequence context^69^, this allows calculation of differences in kinetics between molecules with and without the event within the same sequence context. Next, we performed kinetic analyses of interpulse duration (IPD) and pulse width (PW). Kinetics values (IPD or PW) for each consensus read were transformed into units of time (seconds) and normalized by the average kinetics values of all bases in the consensus read to correct for baseline sequencing kinetics differences between molecules. For each C>T call, we extracted the kinetics values of all overlapping reads for ± 30 base pairs flanking the event position relative to the reference genome coordinates using each read’s CIGAR value to account for insertions or deletions in the read relative to the reference genome. Next, for each C>T call, we calculated the ratio of kinetics values for each base position by dividing the kinetics values (IPD or PW) of the molecule with the call by the weighted average kinetics values of molecules without the call (the weighted average weights by each molecule’s number of passes; i.e., its ‘ec’ tag value). Finally, for each flanking and mutant base position, we calculated the average and standard error of the mean of the kinetics value ratios across all C>T calls of each sample or sample set of interest. The same kinetic analysis was performed for dsDNA C>T mutation calls (i.e., bona fide cytosine to thymine double-strand mutations) in non-heat treated blood DNA, 56 °C and 72 °C heat treated blood DNA, sperm, kidney, and liver samples (all sequenced by HiDEF-seq v2), for the strands synthesized by the sequencing polymerase opposite the strand containing the C>T mutation; this shows the kinetic profile of true C>T changes, as a comparator for C>T calls arising from cytosine damage. Note, the dsDNA C>T mutations used for this kinetics analysis were called with the same thresholds used for ssDNA C>T calls. Both these ssDNA and dsDNA analyses were additionally conducted after randomization of labels among molecules with and without the C>T call to confirm the kinetic signal was specific to molecules with the C>T call. The kinetic profile heatmap and clustering was performed with the ‘ComplexHeatmap’ R package^148^.

## Acknowledgements

This work was supported by grants from the Eunice Kennedy Shriver National Institute of Child Health and Human Development (R21HD105910; G.D.E and J.E.S), the NIH Common Fund (DP5OD028158; G.D.E), the Sontag Foundation (G.D.E), the Pew Foundation (G.D.E.), and the Jacob Goldfield Foundation (G.D.E). Sequencing performed at the New York University (NYU) Grossman School of Medicine Genome Technology Center was supported in part by the National Cancer Institute (P30CA016087). The computational work was supported in part by the New York University Information Technology High Performance Computing resources, services, and staff expertise, and by the New York University Grossman School of Medicine High Performance Computing Core. U.T. is supported by a Stand Up To Cancer–Bristol-Myers Squibb Catalyst Research Grant (SU2C-AACR-CT-07-17), SickKids Foundation donors Harry and Agnieszka Hall, Meagan’s Walk (MW-2014-10), BRAINchild Canada, the LivWise Foundation, the Canadian Institutes for Health Research (CIHR; grant 108188), and a Canadian Cancer Society/CIHR/Brain Canada Spark Grant (Spark-21, 707089). J.E.S. is supported by the Damon Runyon Cancer Research Foundation, the Vinney Family Scholars Award, and the Bristol Myers Squibb Foundation. M.G.P. was supported by NIH grants T32AG052909 and F32AG076287. We thank Benjamin Neel (NYU), Hannah Klein (NYU), and Aravinda Chakravarti (NYU) for helpful discussions; Maya Fridrikh, Nancy Francoeur, and Robert Sebra (Genomics Core Facility at the Icahn School of Medicine at Mount Sinai) for assistance with sequencing; Shenglong Wang (New York University Information Technology) for assistance with high performance computing; and the NIH NeuroBioBank for providing human tissues.

## Author Contributions

G.D.E. conceived the technology, with input from M.H.L and J.E.S. G.D.E., M.H.L., B.C., U.C., and J.E.S. designed the experiments. M.H.L., B.C., and U.C. performed technology development experiments. M.H.L. and B.C. prepared sequencing libraries. U.T., V.B., L.S., T.P., and R.E.B. collected some of the samples. E.L., D.R., and A.B.S. recruited research subjects for sperm samples. D.R. prepared ZyMot sperm samples. M.H.L., R.C.B., A.S., Z.M., C.A.L., T.K.T., and G.D.E prepared tissues and cell samples. M.G.P. prepared NanoSeq libraries. G.D.E. created the computational pipeline with input from U.C. M.H.L., B.C., and G.D.E. performed the analysis. M.H.L. and G.D.E. wrote the initial manuscript, with input from B.C. and J.E.S. All co-authors contributed to the final manuscript.

## Supplementary Information

### Extended Data Figures

**Extended Data Figs. 1-10**, including Extended Data Figure Legends and Extended Data References.

### Supplementary Notes

**Supplementary Notes 1-3**, including **Supplementary Figs. 1-2** and Supplementary Notes References.

### Supplementary Tables

This file contains Supplementary **Tables 1-4. Supplementary Table 1** contains details of samples profiled in the study and sequencing statistics. **Supplementary Table 2** lists ssDNA and dsDNA call burdens for all samples. **Supplementary Tables 3 and 4** show the raw counts and spectra of ssDNA calls and dsDNA mutations, respectively. Spectra (normalized to sum = 1) are corrected for the trinucleotide content of the genome relative to the trinucleotide content of the interrogated bases, except for ssDNA calls of NanoSeq.

## Data availability

Data generated in this study (FASTQ files for Illumina sequencing; consensus sequence BAM files for PacBio data) will be made available at the NCBI database of Genotypes and Phenotypes (dbGaP) [accession ID pending].

## Code availability

The source code for the analysis pipeline is available at https://github.com/evronylab/HiDEF-seq.

