## Extended Data Figures for "Single-strand mismatch and damage patterns revealed by single-molecule DNA sequencing"

#### Extended Data Figure Legends

Extended Data Fig. 1. HiDEF-seq library preparation and sequencing metrics. **a**, Representative DNA sizing electropherogram after Hpy166II restriction enzyme digestion (top) and after completion of the HiDEF-seq library preparation, which removes fragments < 1 kb (bottom). **b**, Two-dimensional histogram of all molecules from a representative HiDEF-seq sequencing run of each molecule's longest strand read length (bp, base pairs) versus its total polymerase read length (PRL). Dashed line signifies the expected strand length distribution. The red diagonal line reflects 18% of molecules with < 1 strand pass, which is typical in PacBio sequencing. **c**, Histogram (200 bp bins) for representative HiDEF-seq samples (n=51) of molecule consensus sequence lengths (i.e., molecule sizes). Line and shaded region show average and standard deviation, respectively, across samples for each bin. The average of these samples' median lengths is 1.7 kilobases (kb). **d**, Histogram as in panel (c), showing HiDEF-seq (n=51 representative samples) yields smaller molecule lengths than standard PacBio (HiFi) samples (n=10 samples). The average of samples' median lengths are 1.7 kb and 18.3 kb for HiDEF-seq and HiFi, respectively. **e**, Two-dimensional histogram of the number of passes (bin width of 5 passes) vs. consensus sequence lengths (bin width of 200 bp) for molecules from the 51 representative HiDEF-seq samples plotted in panels (c,d). Bins are colored if there is at least one molecule in the bin. **f**, Box plots of the fraction of a molecule's consensus sequence bases (average of forward and reverse strands) that have the maximum predicted quality (quality=93, as predicted by ccs, **Methods**) versus the number of passes per strand, across all molecules of the same samples included in panels (c-e). Note: 93 is the quality required for HiDEF-seq analysis. This plot illustrates that the number of passes is a key determinant of consensus quality in both HiDEF-seq and HiFi.
**b**, Plot generated by SMRT Link (Pacific Biosciences) software. **c**-**e**, The single-molecule consensus sequence length is the average of the forward and reverse strand lengths. Bin values are normalized to the bin with the highest molecule count. **e**,**f**, The number of passes per strand is the average of the forward and reverse strand 'ec' tags (**Methods**). **c**-**f**, Plots show data of HiDEF-seq molecules that are output by the primary data processing step of the HiDEF-seq analysis pipeline and standard PacBio HiFi molecules that are output by the ccs HiFi pipeline (**Methods**). **f**, Box plot: middle line, median; boxes, 1st and 3rd quartiles; whiskers, the maximum/minimum values within 1.5 x interquartile range. X-axis: square brackets and parentheses signify inclusion and exclusion of interval endpoints, respectively.

Extended Data Fig. 2. Schematic of analysis pipeline. Primary data processing (blue) is followed by call filtering (green) along with germline sequencing analysis (orange), which is then followed by call burden and signature analysis (purple). See **Methods** for full details. On the left of primary data processing steps are the average percentage of molecules filtered by each step across 17 representative HiDEF-seq sequencing runs. Approximately half of molecules filtered by the 'Generate consensus sequence' step are molecules with less than 3 full-length passes (default setting of the ccs tool that creates consensus sequences), and the other half are due to molecules with read quality ('rq' tag) < 0.99. At the end of the call filtering steps are listed the percentage of bases filtered by all the call filtering steps, calculated out of the total bases of molecules that pass primary data processing, for the same 17 representative HiDEF-seq sequencing runs. The filter for 'low-quality genomic regions and gnomAD variants with allele frequency (AF) > 0.1% in the population' covers approximately 15% and 7% of the genome when using Illumina and PacBio germline sequencing data, respectively (i.e., when PacBio germline sequencing data is used, the pipeline uses less restrictive filters due to fewer genome alignment errors and artifacts). WGS, whole-genome sequencing.

Extended Data Fig. 3. Analysis thresholds and comparison of analyses using short- versus long-read germline sequencing. **a**, Histogram of predicted consensus sequence accuracy ('rq' tag, bin width=0.0001) for DNA molecules that pass primary data processing steps from 3 representative HiDEF-seq v2 sperm samples (21yo: SPM-1002; 39yo: SPM-1004; 44yo: SPM-1020; yo, years old). Note, these are consensus sequence accuracies predicted by the ccs consensus calling software (**Methods**), which are used to filter low-quality molecules, but these accuracies do not reflect the true accuracy that is significantly higher. **b**, Box plot of passes per strand for different consensus sequence accuracy bins, for molecules from the 3 samples included in the prior panel, showing that higher minimum accuracies select for molecules with higher numbers of passes. **c**, Fraction of post-primary data processing molecules that are filtered (left plot) and fraction of post-primary data processing base pairs that remain for interrogation (right plot) using different minimum passes per strand and consensus sequence accuracy thresholds. Values show average of the 3 samples included in the prior panels, after completing all steps of the mutation filtering pipeline. **d**,**e**, dsDNA mutation burdens for the 3 samples included in the prior panels using different minimum passes per strand and consensus sequence accuracy thresholds. Panel (e) shows data from (d) at consensus accuracy of 0.99 with Poisson 95% confidence intervals. These data illustrate stability of dsDNA mutation burden estimates at broad thresholds using sperm as the most stringent test of fidelity. **f**, Fraction of high-quality, known heterozygous germline variants detected using different minimum required fraction of molecule passes (i.e., subreads) that detect the variant (filter applied separately to each strand). This value is used for sensitivity correction (**Methods**). Values show average of the 3 samples included in prior panels. **g**,**h**, dsDNA mutation burdens for the 3 samples included in the prior panels using different minimum required fraction of molecule passes that detect the variant (filter applied separately to each strand), after correcting for sensitivity (g), and using different minimum required distances from the end of the read (h). Panel (g) illustrates that correcting for sensitivity maintains stable burden estimates. The analysis pipeline requires a minimum of 10 bp from the ends of reads to remove rare alignment artifacts, although this does not significantly alter burden estimates. **i**, ssDNA call burdens for the 3 sperm samples included in the prior panels using different minimum passes per strand and consensus sequence accuracy thresholds. Plot shows a small decrease in ssDNA call burdens with a higher minimum required passes per strand at low consensus sequence accuracy thresholds, and convergence to similar burdens at high consensus sequence accuracy thresholds. Data shown with minimum fraction of 0.5 molecule passes that detect the variant. **j**, ssDNA call burdens for the 3 sperm samples included in the prior panels using different minimum required fraction of molecule passes that detect the variant, after correcting for sensitivity. Data shown with a minimum consensus sequence accuracy of 0.999 and a minimum of 20 passes per strand. **k**,**l**, Concordant dsDNA mutation and ssDNA call burdens obtained by HiDEF-seq v2 using short-read (Illumina) or long-read (PacBio, Pacific Biosciences) germline sequencing during analysis, for two samples (1301 and 1901 blood).

**a-d**,**i**, Consensus sequence accuracies are the average of forward and reverse strand accuracies. **b**, Box plot: middle line, median; boxes, 1st and 3rd quartiles; whiskers, the maximum/minimum values within 1.5 x interquartile range. X-axis: square brackets and parentheses signify inclusion and exclusion of interval endpoints, respectively. **c-e**,**i**, Threshold for minimum required passes per strand is applied to both strands. **c-j,** The symbols ‡ and § mark the final thresholds chosen for dsDNA and ssDNA analyses, respectively. **c**,**f**, Error bars: standard deviation; note, panel (f) error bars are small and therefore not well visualized. **d,e,g-l**, mutation and call burdens are corrected for sensitivity and trinucleotide context opportunities of the full genome relative to interrogated bases (**Methods**). **e**,**g,h**,**j-l**, Error bars: Poisson 95% confidence intervals.

Extended Data Fig. 4. HiDEF-seq v1 dsDNA mutation burdens and removal of ssDNA artifacts by HiDEF-seq v2. **a**, dsDNA mutation burdens in two sperm samples (left to right: SPM-1004, SPM-1020) profiled by both HiDEF-seq v1 and NanoSeq, compared for each age (yo, years old) to paternally-phased de novo mutations in children from a prior study of 2,976 trios[1]. **b**, dsDNA mutation burdens versus age, measured by HiDEF-seq v1. Dashed lines (liver, kidney): weighted least-squares linear regression. Dotted lines (blood, neurons): these only connect two data points to aid visualization of burden difference, since regression cannot be performed with two samples. **c**, Mutational signature contribution to dsDNA mutations detected in HiDEF-seq v1 samples. All samples, except blood from a 62-year-old individual (1901), were jointly analyzed with fitting of SBS1 and de novo extraction of one additional signature SBSB (**Methods**). The blood sample of the 62-year-old, who has a history of end-stage renal disease (**Supplementary Table 1**), was analyzed separately together with 5 other HiDEF-seq v2 blood samples from this individual due to identification of an additional signature SBSC. Analysis of samples grouped by tissue type, excluding the 62-year-old blood sample, produced similar results. For de novo extracted signatures (SBSB and SBSC), the cosine similarities to the closest matching COSMIC signatures are shown in parentheses. Sperm samples and kidney and liver samples from an infant (1443) were not included here since the number of mutations is too low for reliable signature extraction. **d**, Burdens of dsDNA mutations (left) and ssDNA calls (right) of a blood sample (individual 1301) measured by HiDEF-seq v1 (without nick ligation), and HiDEF-seq v2 (with nick ligation). Nick ligation eliminates T>A ssDNA artifacts that match the illustrated GTTBVH motif. The motif was derived using the ggseqlogo R package[2] using all ssDNA T>A calls from the HiDEF-seq v1 sample. Gray bar is calls matching the motif with log-odds score > 2 calculated with the score_match function of the universalmotif R package. **e**,**f**, Proposed mechanism for the GTTBVH motif of ssDNA artifactual calls. The known GTNNAC motif of the Hpy166II restriction enzyme used in HiDEF-seq may arise if Hpy166II operates as a dimer (cut sites signified by triangles) with each monomer binding opposite strands, and the GTTBVH motif is due to intersection (∩) and union (U) combinatorial logic for the outer and inner 2 bases, respectively (e). ssDNA GT[T>A]BVH artifactual calls may arise from rare Hpy166II monomer nicking events, pyrophosphorolysis of the 'T' upstream of the nick, and addition of a mismatched 'A' during the Klenow dATP/ddBTP A-tailing reaction. Further extension with ddBTP does not occur due to the mismatch[3] (f). **g**, HiDEF-seq v2's nick ligation increases library yield by 66% for post-mortem tissues, likely by repairing nicks in the original input DNA so that the molecules are not eliminated in the final nuclease treatment step. Number of samples per group (left to right): 8, 8, 5, 9 (**, p=0.002; ns, not significant; two-sided unpaired t-test).
**a**, Box plots: middle line, median; boxes, 1st and 3rd quartiles; whiskers, 5% and 95% quantiles. For each sample, HiDEF-seq and NanoSeq confidence intervals were normalized to reflect an equivalent number of interrogated base pairs (**Methods**). **a**,**b**,**d** Error bars: Poisson 95% confidence intervals. **g**, Error bars: standard deviation.

Extended Data Fig. 5. HiDEF-seq v2 without A-tailing removes ssDNA artifacts of post-mortem tissues with fragmented DNA. **a**, Fraction of ssDNA calls that are T>A (corrected for trinucleotide context opportunities) versus the ssDNA T>A burden in all samples profiled with standard HiDEF-seq v2 (i.e., with A-tailing: Klenow reaction +dATP/+ddBTP) from healthy individuals and cell lines (i.e., excluding cancer-predisposition syndromes). Post-mortem kidney and liver consistently have the highest fraction of ssDNA calls that are T>A. **b**, Standard HiDEF-seq v2 (with A-tailing) ssDNA call spectrum for a liver sample with a high ssDNA T>A burden ($6.8 \cdot 10^{-7}$), corrected for trinucleotide context opportunities. Parentheses show total number of calls. **c**, Correlation between ssDNA T>A artifact burden and the input DNA's DNA Integrity Number measured by TapeStation electrophoresis[4] across all samples profiled with standard HiDEF-seq v2 (with A-tailing) from healthy individuals and cell lines (i.e., excluding cancer-predisposition syndromes). Lower DNA Integrity Number corresponds to more fragmented DNA.

**d**, Proposed mechanism for the ssDNA T>A artifact calls in fragmented DNA when performing standard HiDEF-seq v2 (with A-tailing). **e**, Modifications of the standard HiDEF-seq v2 protocol to eliminate ssDNA T>A artifacts in fragmented DNA. All trials were from the same DNA extraction aliquot (liver from individual 5697). See **Methods** for details. PNK, polynucleotide kinase; Bst, Bst large fragment; min, minutes. **f**, ssDNA call spectra for three of the samples shown in panel (e): standard HiDEF-seq v2 with A-tailing (top, same spectrum as panel (b) ), HiDEF-seq v2 with a Klenow reaction that does not contain dATP nor ddBTP (middle), and HiDEF-seq v2 with a Klenow reaction containing only ddBTP (bottom). The total number of ssDNA calls and total ssDNA call burden (calls per base) are shown. **g,** Fraction of ssDNA calls that are T>A (corrected for trinucleotide context opportunities) versus the ssDNA T>A burden in post-mortem kidney (n=5) and liver (n=5) samples profiled with HiDEF-seq v2 without A-tailing (i.e., Klenow reaction -dATP/+ddBTP). **h,** Concordant dsDNA mutation burdens in sperm sample SPM-1013 measured by standard HiDEF-seq v2 (i.e., with A-tailing) and HiDEF-seq v2 without A-tailing (i.e., Klenow reaction -dATP/+ddBTP). yo, years old. **i,** Mutational signature contribution to dsDNA mutations detected in HiDEF-seq v2 primary human tissues from individuals without cancer predisposition. Post-mortem liver and kidney samples were profiled by HiDEF-seq v2 without A-tailing. All samples, except blood from a 62-year-old individual (1901), were jointly analyzed with fitting of SBS1 and de novo extraction of one additional signature SBSB. Blood samples of the 62-year-old (with a history of end-stage renal disease; **Supplementary Table 1**) profiled by HiDEF-seq v2 were analyzed separately (plot shows average signature contributions across 5 blood samples) due to identification of an additional outlier signature SBSC. Analysis of samples grouped by tissue type, excluding the 62-year-old blood sample, produced similar results. For de novo extracted signatures (SBSB and SBSC), the cosine similarities to the closest matching COSMIC signatures are shown in parentheses. Sperm, kidney and liver samples from an infant (1443) and 18-year-old (1409), and blood from a 4-year-old (5203) are not included here since their number of mutations are too low for reliable signature extraction.
**e,h,** Error bars: Poisson 95% confidence intervals. **e-g,** Rxn, reaction.

<u>Extended Data Fig. 6. Comparison of HiDEF-seq and NanoSeq.</u> **a,** Comparison of HiDEF-seq versus NanoSeq dsDNA mutation spectra for individual 63143. **b,** Comparison of HiDEF-seq versus NanoSeq ssDNA call burdens, separated by call type. For each call type (i.e., C>A, C>G, etc.), each bar represents a different sample. Samples for each call type, from left to right, are 1105 and 6501 for healthy blood; 63143 for POLE blood; and 1443 for kidney. Comparison for sperm samples is shown in **Fig. 1g**. **c,** Comparison of HiDEF-seq versus NanoSeq ssDNA call spectra for 6501 (Blood, 43 yo), 63143 (POLE blood), and SPM-1060 (sperm, 49 yo).
**a-c,** yo, years old; mo, months old.

<u>Extended Data Fig. 7 dsDNA mutation burdens and patterns in cancer-predisposition syndromes.</u> **a,** Fraction of dsDNA mutations in each context. Non-cancer predisposition samples are (left to right): Blood (B) 5203, 1105, 1301, 6501, and 1901; lymphoblastoid cell line (LCL) GM12812; primary fibroblasts GM02036 and GM03348. Cancer predisposition samples are (left-to-right, in the same order and annotated sample types as top-to-bottom cancer predisposition samples in panel (c) ): GM16381, GM01629, GM28257, 55838, 58801, 57627, 1400, 1324, 1325, 60603, 59637, 57615, 63143 (LCL), 63143 (B), CC-346-253, CC-388-290, CC-713-555. Affected genes annotated below. Note, GM02036 (asterisk) has a significant increase in C>T mutations with a spectrum matching COSMIC SBS7a (ultraviolet light exposure), likely due to the fibroblasts deriving from sun-exposed skin. **b,** Representative dsDNA mutation spectra of a sample for each affected gene, corrected for trinucleotide context opportunities. Sample IDs are in parentheses. Ages (yo, years old) are listed for blood samples. **c,** Fraction of dsDNA mutations attributable to de novo extracted dsDNA mutational signatures.

Sample genotypes are on the right (hom., homozygous; compound heterozygous variants separated by '/'). Cosine similarity to highest matching COSMIC signature is shown in parentheses. For SBSF, the similarity to SBS18 and SBS36 are also shown since these have been previously associated with *MUTYH*. These *MUTYH* signatures were not extracted due to the normal mutation burdens of our *MUTYH* blood samples (see panel (d) ), which is expected at these sample ages and our interrogated base coverage[5]. Note that SBS40 resembles SBS18 and SBS36 in the C>A spectrum that is enriched in *MUTYH* syndrome[5]. Signature extraction was performed for samples of each DNA repair pathway (except *XPC* separately from *ERCC6*/*ERCC8*), while simultaneously fitting COSMIC SBS1 and SBS5 (**Methods**). Samples are in the same top-to-bottom order as left-to-right cancer predisposition samples in panel (a). **d,** dsDNA mutation burden per base pair divided by the age of the individual in years at the time of blood collection, corrected for trinucleotide context opportunities and sensitivity. Only blood samples are shown, since blood can be annotated with the age of the individual. Accordingly, since we did not profile blood samples nucleotide excision repair syndrome, this category is not shown. Non-cancer predisposition blood samples are the same (left-to-right) as in panel (a) (left-to-right). Cancer predisposition blood samples are the same (left-to-right) as blood samples in panel (c) (top-to-bottom). Affected genes annotated below. **e,** In *POLE* samples, dsDNA mutations occur more often with AGA>ATA on the non-reference (-) than on the reference (+) strand in genomic loci where the non-reference strand is synthesized more frequently in the leading direction (i.e., positive fork polarity) based on replication timing data (**Methods**). Random loci are the average of 50 sets of 1,000 random genomic loci with either the sequence AGA or TCT for which there is Repli-seq data at the locus. Reference (+) strand refers to the plus strand of the human reference genome. X-axis is fork polarity, divided into 9 quantile bins from 0 to 1, with higher values corresponding to a greater probability of the non-reference strand being copied in the leading rather than lagging strand direction (Methods). Y-axis for POLE HiDEF-seq samples is the 'strand ratio', calculated as the fraction of all AGA>ATA non-reference strand mutations that are in the fork polarity quantile bin divided by the fraction of all AGA>ATA reference strand mutations that are in the fork polarity quantile bin. The 'strand ratio' for random genomic loci is calculated as the fraction of all AGA non-reference strand loci that are in the fork polarity quantile bin divided by the fraction of all AGA reference strand loci that are in the fork polarity quantile bin. POLE samples are the same top-to-bottom order in the legend as top-to-bottom POLE samples in panel (c). Asterisks signify statistical significance in comparison of the POLE 4-sample average (dashed line) to random loci (heteroscedastic two-tailed t.test); p-values left-to-right for asterisks: $3.7 \cdot 10^{-17}$, 0.001, 0.009, 0.02, 0.003. An analysis excluding mutations overlapping genes, to exclude biases due to transcription strand, produced similar results with significant p-values of $p=3.1 \cdot 10^{-10}$, 0.003, and 0.04 for quantiles 0-0.1, 0.1-0.2, and 0.6-0.7, respectively, but this analysis has reduced power due to the 55% reduction in the number of mutations analyzed. Note, ssDNA calls are not plotted here due to the small number of ssDNA calls per fork polarity quantile bin. Instead, ssDNA calls are plotted in **Fig. 2h** for negative versus positive fork polarity.
**a-e**, See additional samples details in **Supplementary Tables 1-4**. **e,** Error bars, standard deviation.

<u>Extended Data Fig. 8. Burdens of ssDNA C>T calls, kinetic interpulse duration profiles, and profiling of heat treatment in varied buffers.</u> **a,** Fraction of ssDNA calls that are C>T (corrected for trinucleotide context opportunities) across all HiDEF-seq v2 samples from healthy individuals and cell lines (i.e., excluding cancer-predisposition syndromes), versus the ssDNA C>T burden. Data shown for kidney and liver samples profiled with HiDEF-seq v2 without A-tailing. Sperm consistently have the highest fraction of ssDNA calls that are C>T. LCL, lymphoblastoid cell line. **b,** Average ratio of interpulse duration of C>T sequencing calls and 30 base pairs flanking the call, of each molecule with the call relative to molecules aligning to the same locus without

the call (**Methods**). Data includes the same samples and calls as in **Fig. 4g**. **c,** Average ratio of pulse width (left column) and interpulse duration (right column) after randomizing labels of molecules with and without the calls, for the same samples and calls as in **Fig. 4g**. **d,** dsDNA mutation and ssDNA call burdens of heat-treated blood DNA in an additional experiment testing the effect of different buffers and different DNA extraction methods (orange underline, Puregene alcohol precipitation; all other samples, MagAttract with magnetic beads). MgAc, magnesium acetate; $MgCl_2$, magnesium chloride; KCl, potassium chloride; KAc, potassium acetate; Alb, albumin; Tris buffer is Tris-HCl except for the MgAc/KAc/Alb that is Tris-Acetate (see **Supplementary Table 1** for concentrations). Non-heat treated DNA samples were placed on ice for 6 hours. The percentage of ssDNA sequencing calls that are C>T are annotated above each sample. Cosine similarity to COSMIC dsDNA signature SBS30 is annotated below each sample, after collapsing ssDNA calls to central pyrimidine trinucleotide contexts and correcting for trinucleotide context opportunities, except for the no-heat treatment samples that do not have sufficient C>T calls ('N/A'). **e,** SBS30ss* signature (reproduced from **Fig. 4d**) compared to spectra of ssDNA calls after 72 °C heat damage of blood DNA for 6 hours (h) in only 10 mM Tris buffer (n=10,852 calls) or only water (n=2,751 calls). Spectra are plotted after correcting for trinucleotide context opportunities. Bottom, odds ratios of spectrum contributions at C>T contexts of the Tris-only and water-only samples compared to SBS30ss* (which was derived from sperm and salt-buffer heat-treated samples). Pyr, pyrimidine, Pur, purine. **f,** Heat map of average pulse width ratios for ssDNA and dsDNA C>T calls for positions -1 to +6, for blood DNA samples heated at 72 °C for 6 hours in different buffers or water, and for additional samples for comparison. Unbiased clustering (dendrogram) separates kinetic profiles of ssDNA C>T calls from dsDNA C>T calls and from kinetic profiles after randomizing labels of molecules with and without the calls. dsDNA 'Blood, heat': blood DNA heat-treated at 56 °C and 72 °C (both 3h and 6h for each); dsDNA 'Blood': 4 samples, not heat treated. dsDNA 'Kidney and liver': 10 samples, not heat treated.
**b,c,** Error bars, standard error of the mean. **d,** Error bars, Poisson 95% confidence intervals.

<u>Extended Data Fig. 9. ssDNA call burdens and patterns in healthy tissues.</u> **a,b** ssDNA C>T (a) and non-C>T (b) call burdens versus age across all HiDEF-seq v2 primary tissue samples from healthy individuals. Dashed lines: weighted least-squares linear regression, with a 95% confidence interval (shaded ribbon) shown for the statistically significant association for liver. P-values for the regression for liver versus age are 0.0008 and 0.002 for C>T calls, without and with including post-mortem interval (PMI) as a covariate in a multiple linear regression, respectively. P-values for the regression for liver versus age are 0.0003 and 0.002 for non-C>T calls, without and with including post-mortem interval (PMI) as a covariate in a multiple linear regression, respectively. **c,** ssDNA call burden versus PMI for liver and kidney samples. Dashed lines: weighted least-squares linear regression. The regressions are not statistically significant neither for all calls, nor for C>T or non-C>T calls. **d,e,** Fraction of ssDNA calls (d) and dsDNA mutations (e) in each context for all HiDEF-seq samples from healthy individuals and cell lines (i.e., excluding cancer-predisposition syndromes). The number of calls for each sample are listed above the plot. Note, context fractions for samples with low call counts are less reliable. Samples (left to right) are: Sperm: SPM-1013, SPM-1002, SPM-1004, SPM-1020, SPM-1060; Liver: 1443, 1409, 1104, 5697, 5840; Kidney: 1443, 1409, 1104, 5697, 5840; Blood: 5203, 1105, 1301, 6501, 1901 x 5 replicates; Neurons: 5344, 6371; Fibroblasts: GM02036, GM03348; Lymphoblastoid cell line (LCL): GM12812. Note, GM02036 (asterisk) has a significant increase in C>T mutations with a spectrum matching COSMIC SBS7a (ultraviolet light exposure), possibly due to the fibroblasts deriving from sun-exposed skin. **f,** ssDNA call spectra after pooling calls of samples from healthy individuals and cell lines, separately for each tissue, corrected for trinucleotide context opportunities. The corresponding figure for blood is also shown in **Fig. 4a**.

**a-c,** Error bars, Poisson 95% confidence intervals.

Extended Data Fig. 10. Similarity between SBS30ss* and mitochondrial genome heavy strand A>G dsDNA mutations, and mitochondrial ssDNA call spectra. **a,** SBS30ss* (cytosine deamination) spectrum is collapsed to central pyrimidine trinucleotide contexts and compared to mitochondria heavy strand A>G dsDNA mutation spectrum, for different sample sets: (i) HiDEF-seq liver and kidney samples, including liver samples from which mitochondria were enriched (i.e., same set of samples in **Figs. 6a-d**); (ii) 5697 purified liver mitochondria sample only (plot includes 81% of the mutations in (i) ); (iii) Sample set (i), excluding the 5697 purified liver mitochondria sample (plot includes 19% of the mutations in (i) ). Note, the contexts of SBS30ss* are matched with the reverse complement flanking base contexts of mitochondria heavy strand A>G mutations. The number of dsDNA A>G mutations is indicated. **b,** Spectrum of ssDNA calls combined from liver and kidney samples as well as liver samples from which mitochondria were enriched, excluding bulk (i.e., non-mitochondria enriched) samples profiled by HiDEF-seq v2 with A-tailing. This corresponds to the set of samples profiled in **Figs. 6a-d**. The spectrum is corrected for trinucleotide context opportunities.

#### Extended Data References

**Ext. Data Fig. 1**

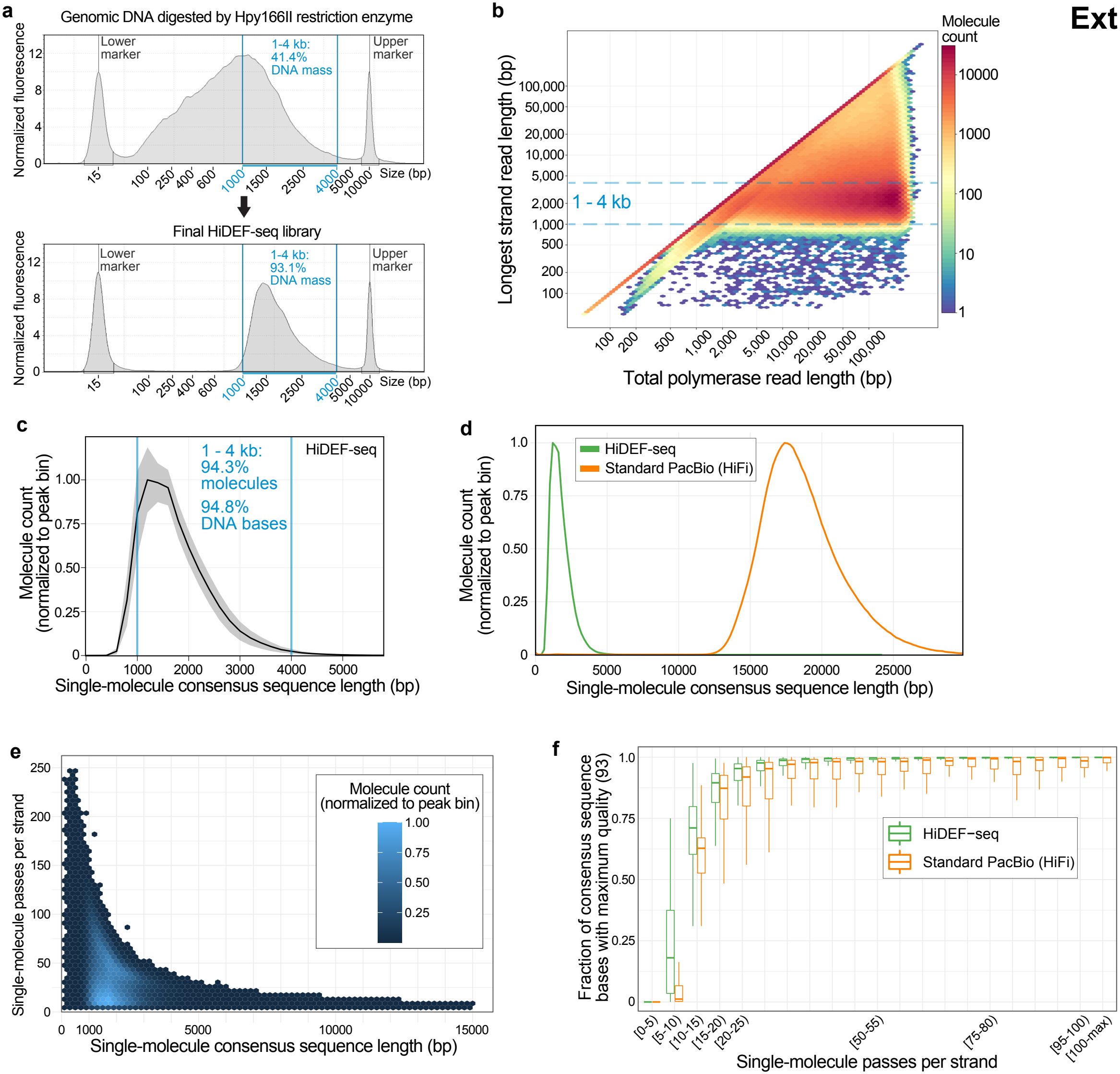

### Ext. Data Fig. 2

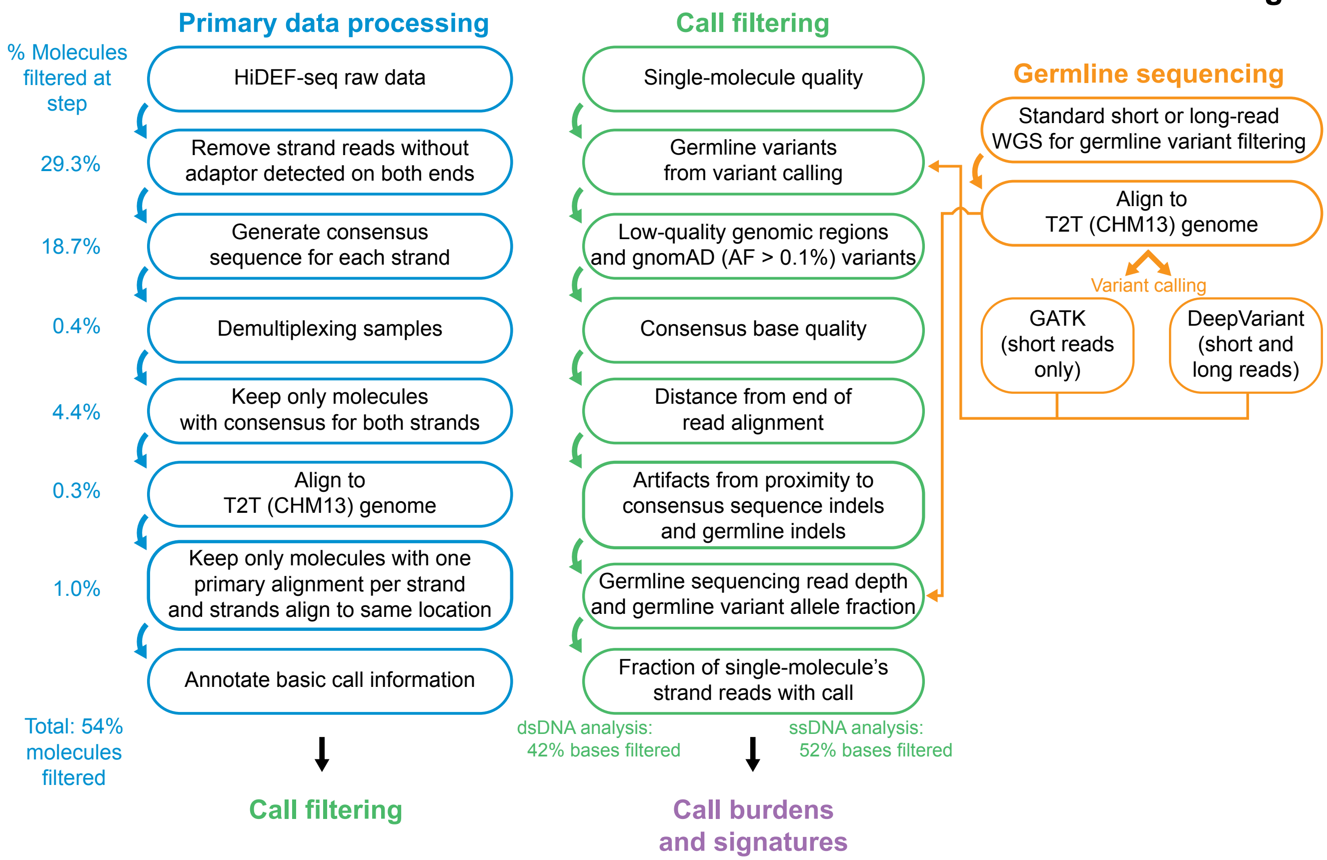

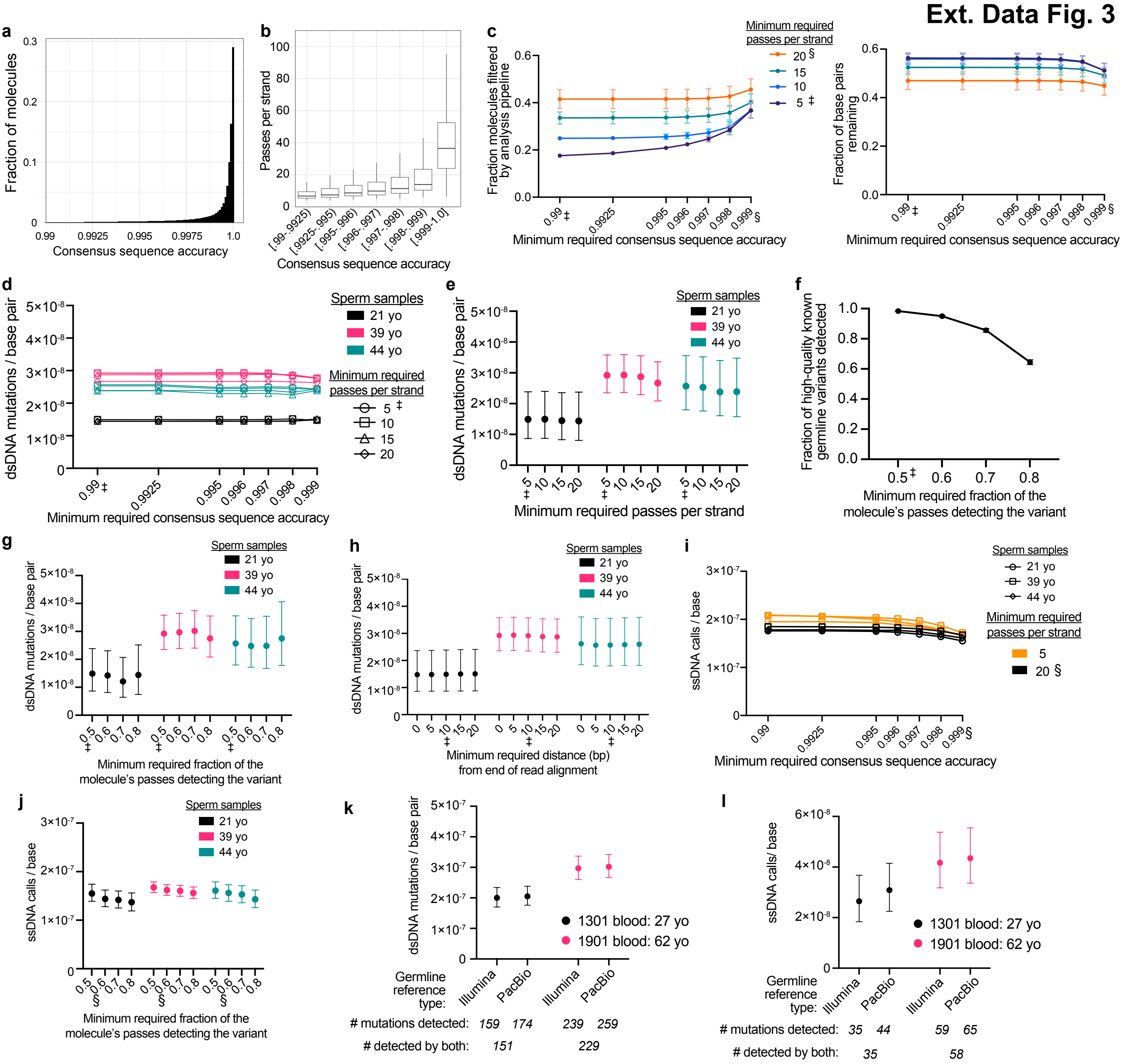
Ext. Data Fig. 3
a
Fraction of molecules
Consensus sequence accuracy
b
Passes per strand
Consensus sequence accuracy
c
Fraction molecules filtered by analysis pipeline
Fraction of base pairs remaining
Minimum required passes per strand
20 §
15
10
5 ‡
Minimum required consensus sequence accuracy
d
dsDNA mutations / base pair
Sperm samples
21 yo
39 yo
44 yo
Minimum required passes per strand
5 ‡
10
15
20
Minimum required consensus sequence accuracy
e
dsDNA mutations / base pair
Sperm samples
21 yo
39 yo
44 yo
Minimum required passes per strand
f
Fraction of high-quality known germline variants detected
Minimum required fraction of the molecule's passes detecting the variant
g
dsDNA mutations / base pair
Sperm samples
21 yo
39 yo
44 yo
Minimum required fraction of the molecule's passes detecting the variant
h
dsDNA mutations / base pair
Sperm samples
21 yo
39 yo
44 yo
Minimum required distance (bp) from end of read alignment
i
ssDNA calls / base
Sperm samples
21 yo
39 yo
44 yo
Minimum required passes per strand
5
20 §
Minimum required consensus sequence accuracy
j
ssDNA calls / base
Sperm samples
21 yo
39 yo
44 yo
Minimum required fraction of the molecule's passes detecting the variant
k
dsDNA mutations / base pair
1301 blood: 27 yo
1901 blood: 62 yo
Germline reference type:
Illumina
PacBio
Illumina
PacBio
### mutations detected: 159 174 239 259
### detected by both: 151 229
l
ssDNA calls/ base
1301 blood: 27 yo
1901 blood: 62 yo
Germline reference type:
Illumina
PacBio
Illumina
PacBio
### mutations detected: 35 44 59 65
### detected by both: 35 58

**Ext. Data Fig. 4**

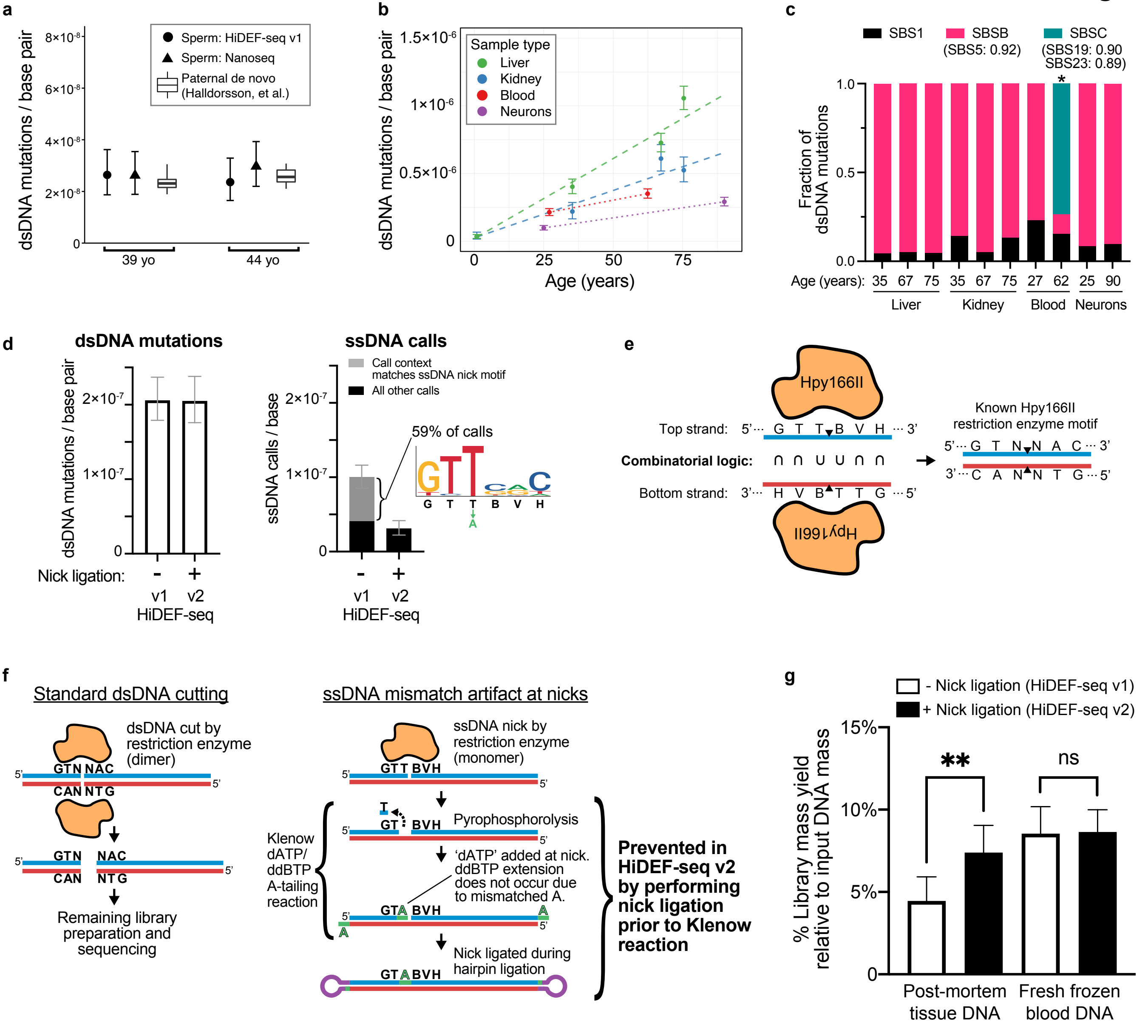

**Ext. Data Fig. 5**

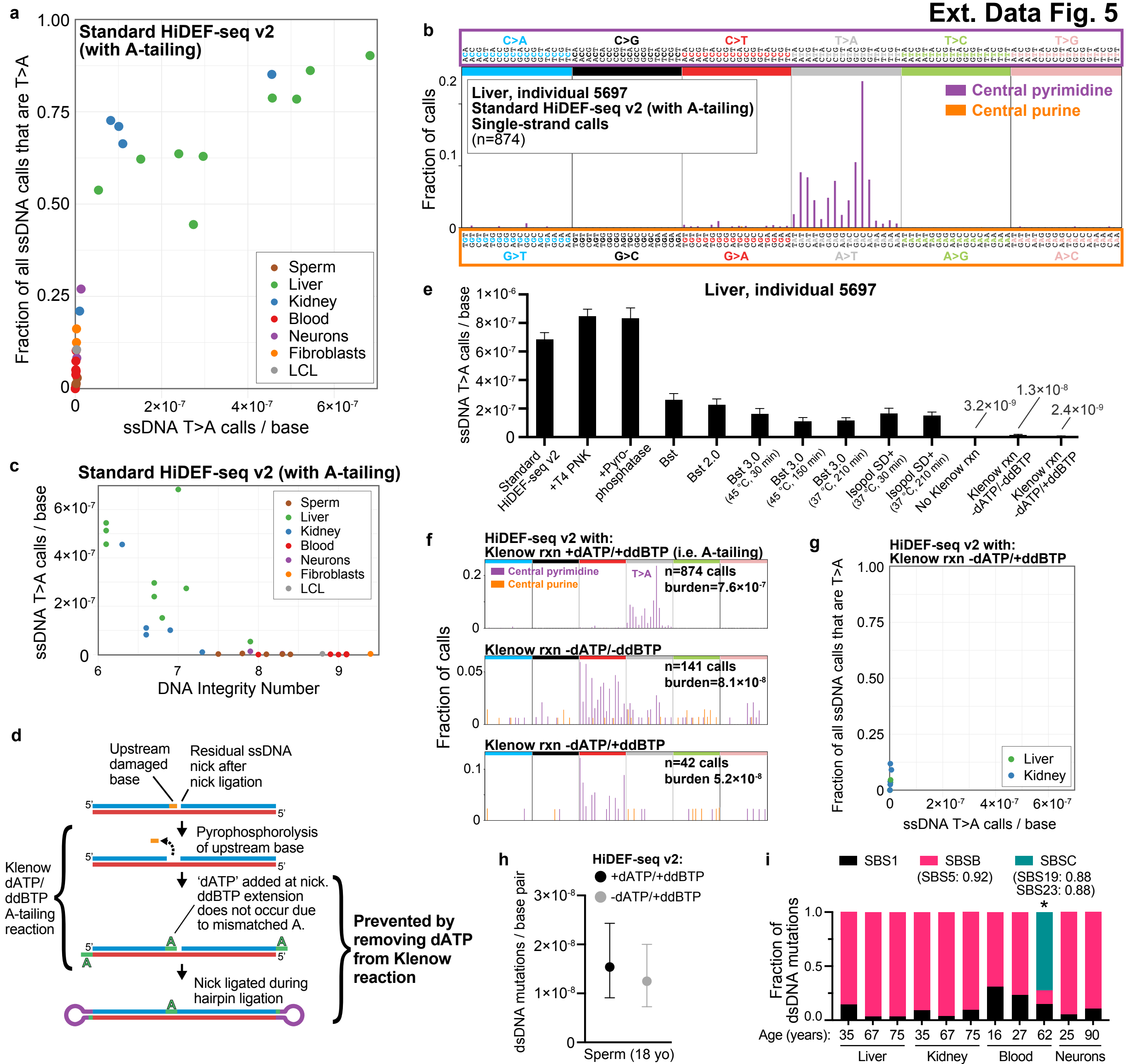

Ext. Data Fig. 6

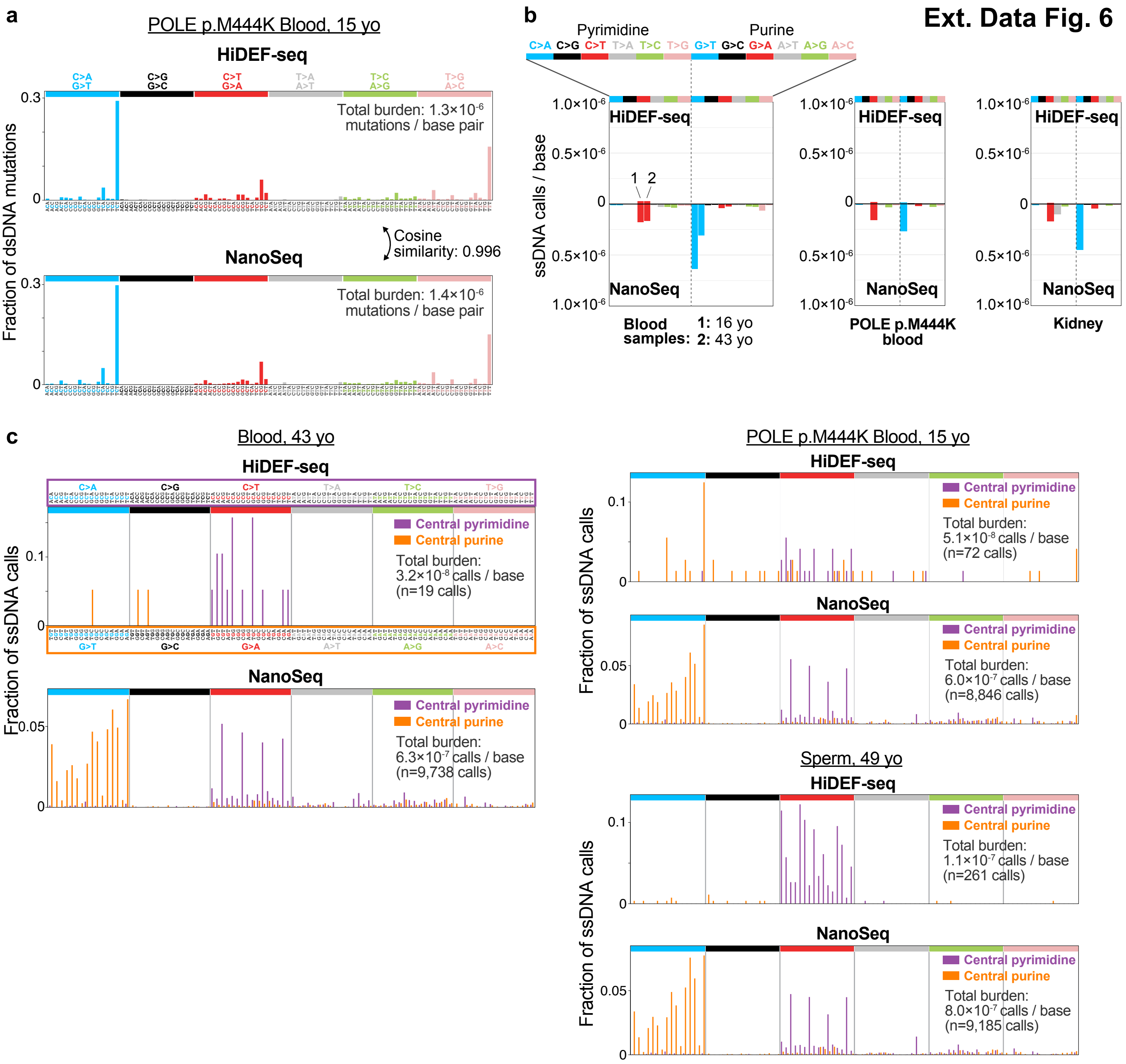

### Ext. Data Fig. 7

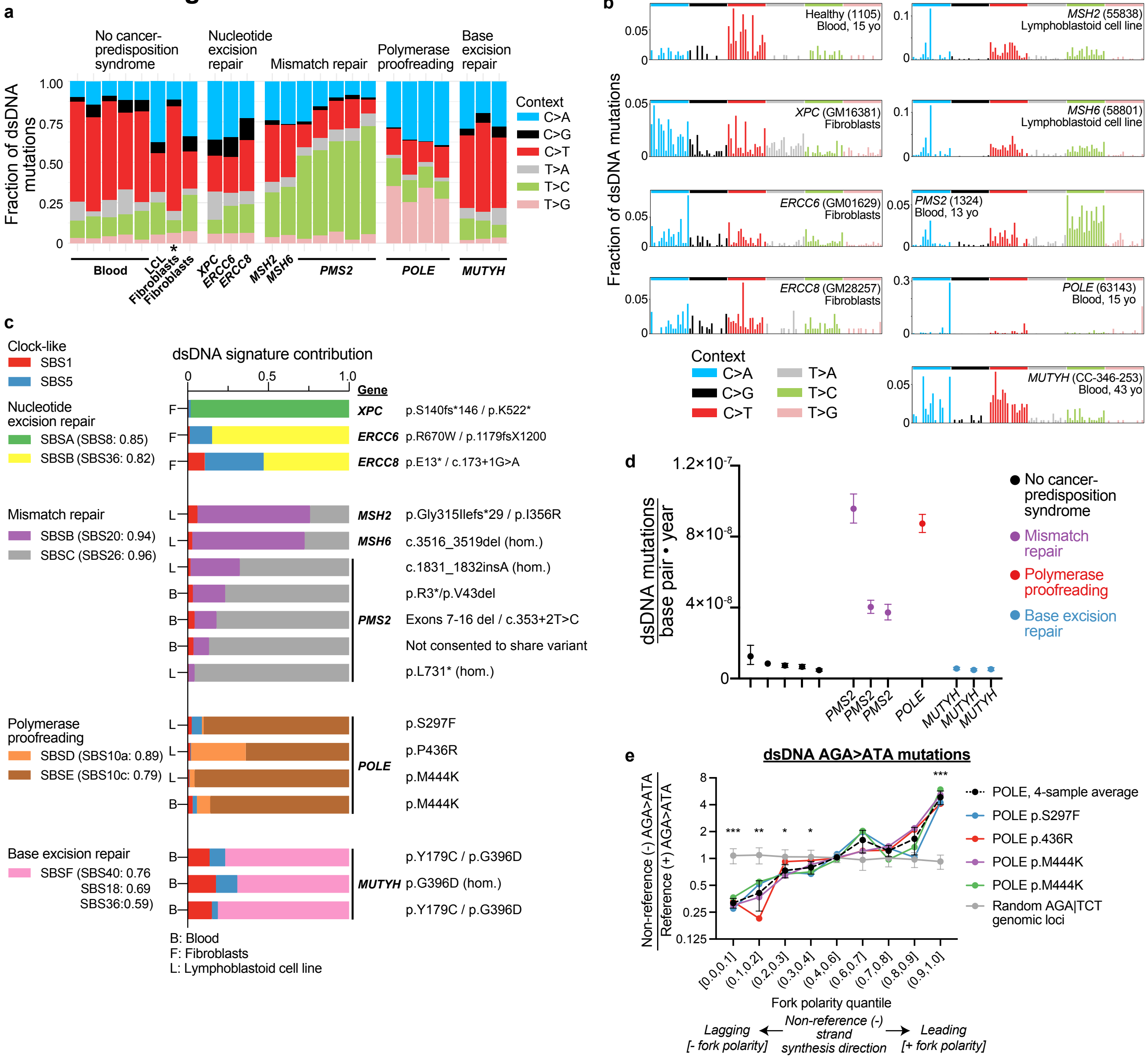

**Ext. Data Fig. 8**

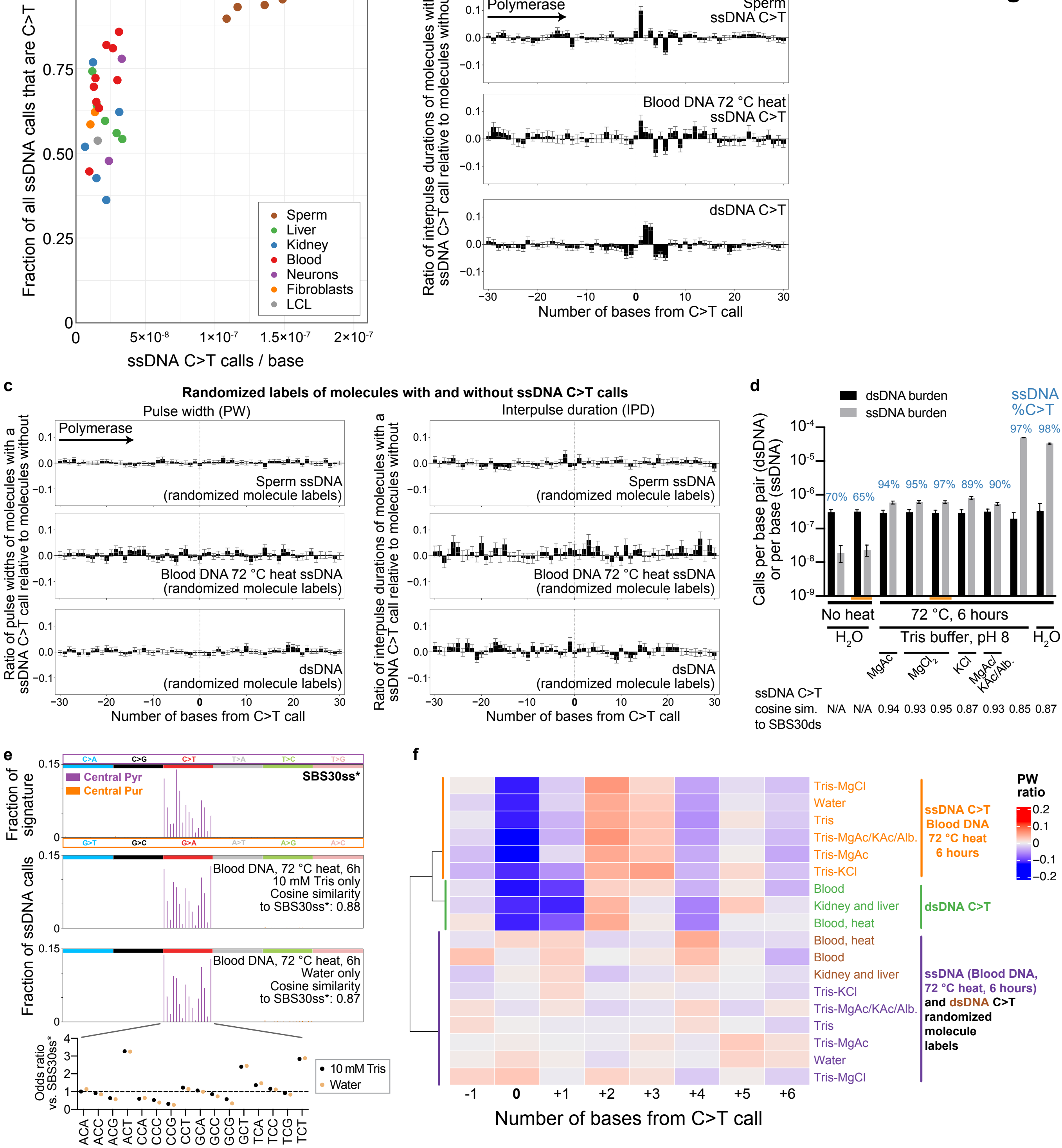

### Ext. Data Fig. 9

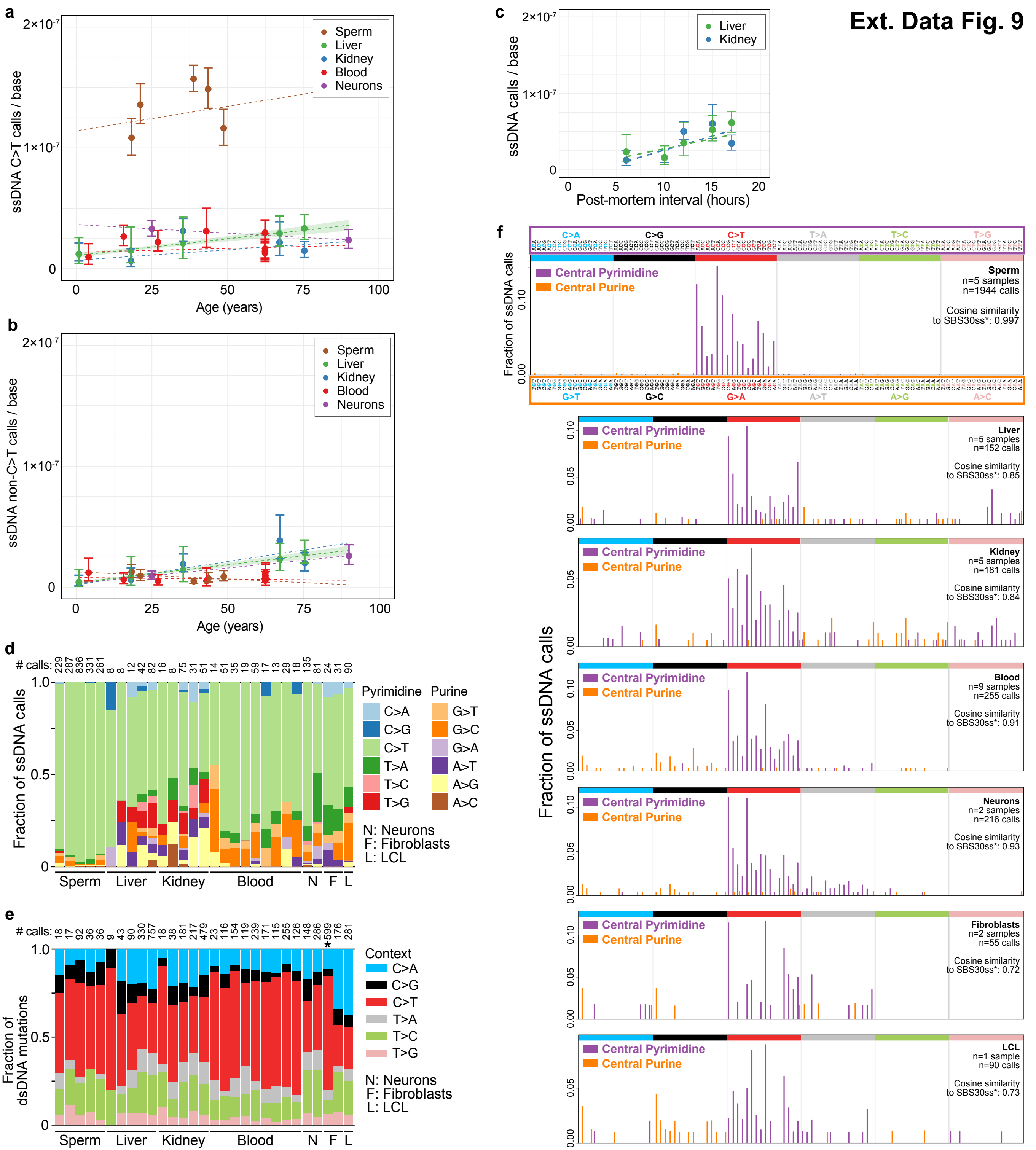

Ext. Data Fig. 10

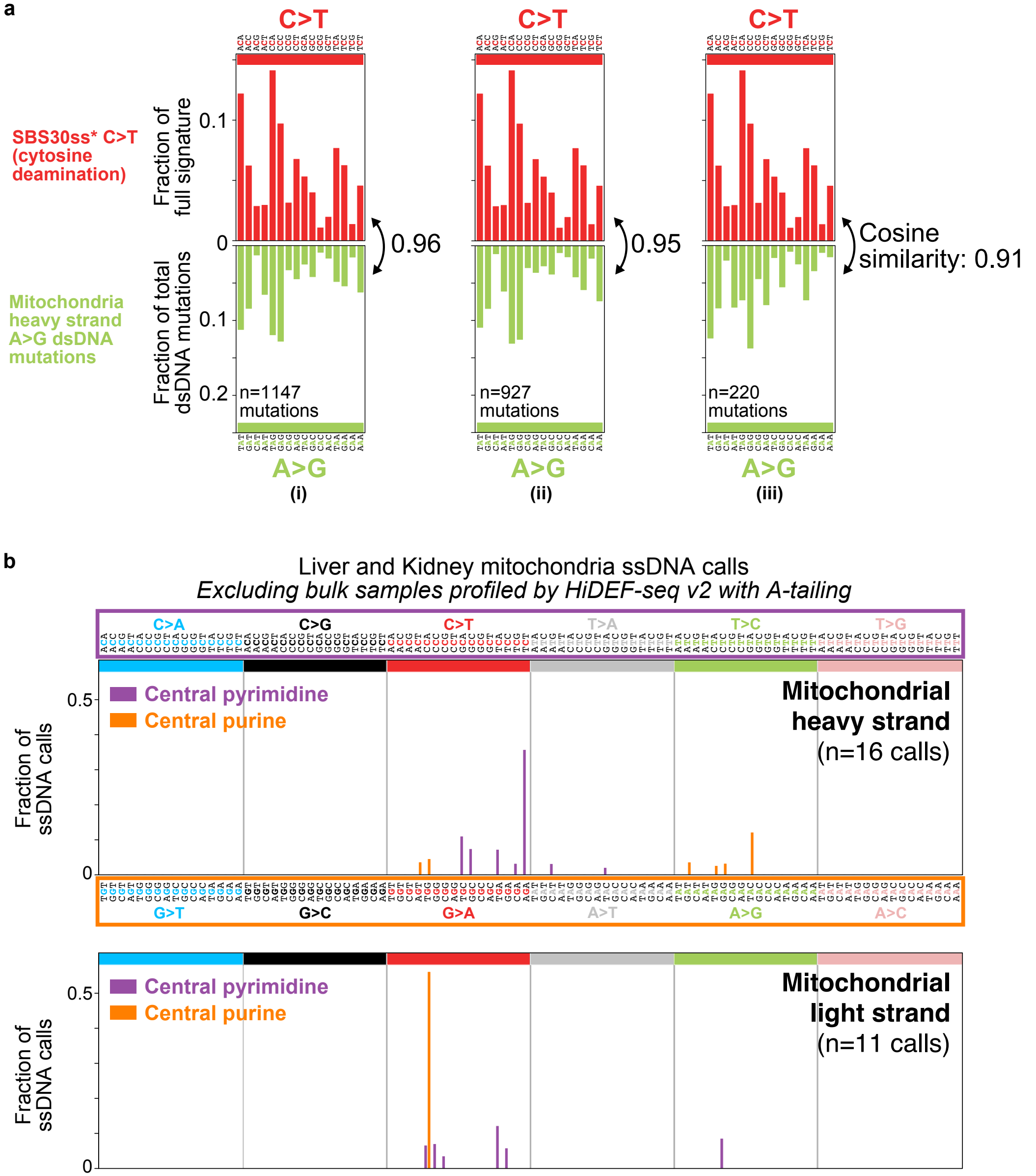
