## Supplementary Notes for "Single-strand mismatch and damage patterns revealed by single-molecule DNA sequencing"

### Supplementary Note 1. HiDEF-seq with larger DNA fragments

Since the number of passes in HiDEF-seq exceeded our initial goals, we reasoned that HiDEF-seq with larger fragments may achieve greater efficiency for double-strand DNA (dsDNA) mutation detection, albeit with reduced efficiency for single-strand DNA (ssDNA) events that require a higher number of passes. As described in the methods for HiDEF-seq utilizing Hpy166II for 1-4 kilobase (kb) libraries, we similarly performed an *in silico* computational screen and experimental screen to identify a blunt-cutting restriction enzyme that fragments the human genome to a larger size range of approximately 1-10 kb. Experimental screening of the top *in silico* candidates identified PvuII as producing the desired fragment size distribution. We prepared large fragment HiDEF-seq libraries with PvuII from two sperm samples (SPM-1002 and SPM-1020; **Supplementary Table 1 and Methods**). As expected, analysis of large fragment HiDEF-seq data showed a larger median fragment size compared to standard size HiDEF-seq (4.2 kb versus 1.7 kb, respectively) and a lower median number of passes (15.2 versus 32, respectively) (**Supplementary Figs. 1a,b**). Standard and large fragment size HiDEF-seq yielded an average of $2.6 \cdot 10^9$ and $3.9 \cdot 10^9$ interrogated dsDNA base pairs per sequencing run, respectively (the large fragment size run was in the 95%ile relative to standard fragment size runs), and $4.3 \cdot 10^9$ and $3.4 \cdot 10^9$ interrogated ssDNA bases, respectively. Therefore, HiDEF-seq with large fragment size improves efficiency for dsDNA mutation interrogation, while reducing efficiency for ssDNA event interrogation. dsDNA mutation and ssDNA call burdens were similar between large and standard fragment size HiDEF-seq for the same samples (**Supplementary Fig. 1c**).

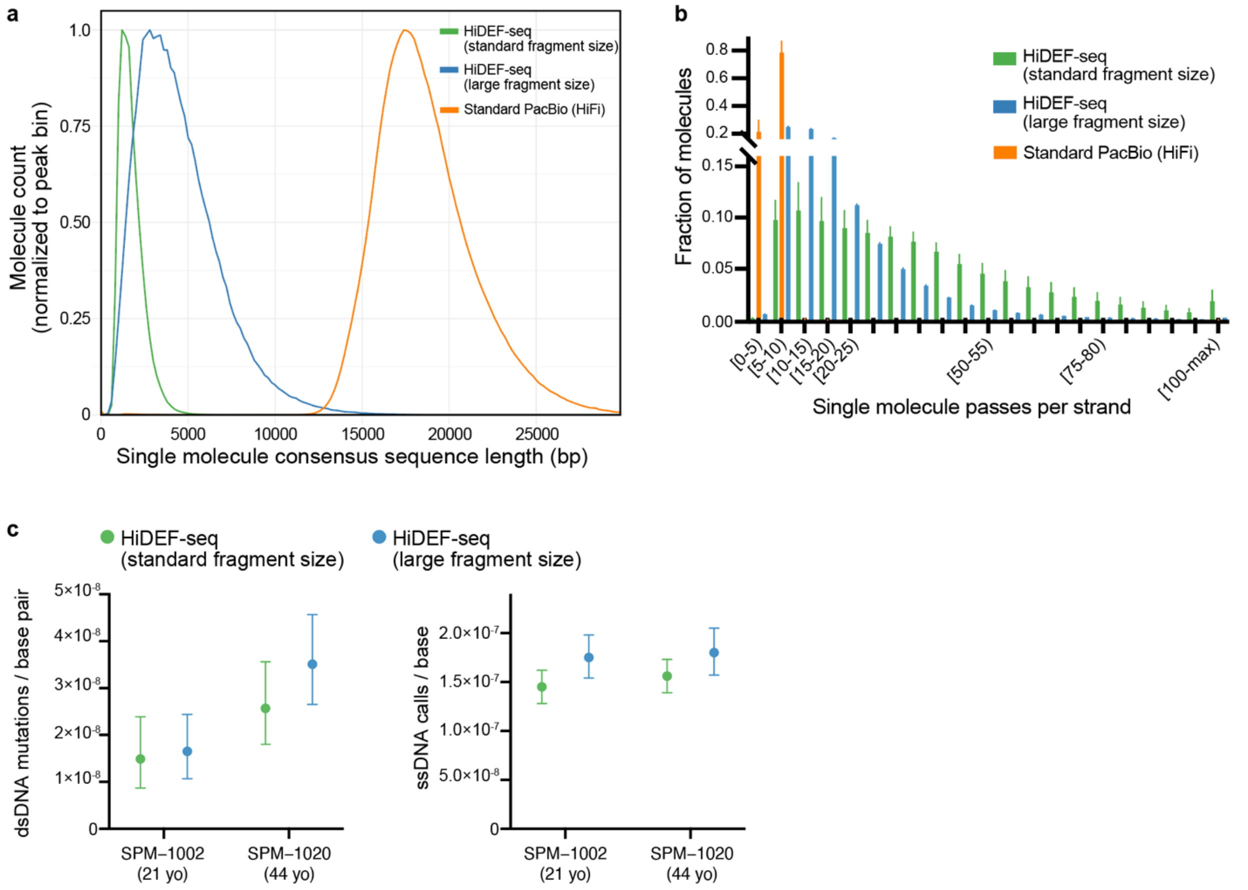

**Supplementary Figure 1. a**, Histogram of consensus sequence lengths (i.e., molecule sizes) for: HiDEF-seq standard fragment size (n=51 samples, median length 1.7 kb), HiDEF-seq large fragment size (n=2 samples, median length 4.2 kb), and standard PacBio (HiFi) (n=10 samples, median length 18.3 kb). Histogram lines show average across samples for each bin. Bin values of each sample type are normalized to the bin with the peak molecule count. **b**, Histogram as in panel (a) of the number of passes per strand (bin width of 5 passes). Average of medians across samples of sample type: 32.0 (standard fragment size), 15.2 (large fragment size), and 5.7 (HiFi) passes per strand. Error bars, standard deviation. **c** dsDNA mutation (left) and ssDNA (right) call burdens of two sperm samples profiled by both standard and large fragment size HiDEF-seq. Error bars, Poisson 95% confidence intervals.

### Supplementary Note 2. Cost efficacy of HiDEF-seq for ultra-high fidelity double-strand DNA mutations

Here, we estimate the cost efficacy of HiDEF-seq for ultra-high fidelity double-strand DNA (dsDNA) mosaic mutation detection, and we compare it to NanoSeq, another state-of-the-art method[1]. Two factors determine cost efficacy: 1) Duplex efficiency: the number of sequenced molecules that can be interrogated at ultra-high fidelity for dsDNA mosaic mutations, and, 2) cost of sequencing.

HiDEF-seq achieves significantly greater duplex efficiency per molecule than prior methods, including NanoSeq, because every sequenced molecule in HiDEF-seq is natively dsDNA on the sequencing instrument. This contrasts with Illumina-based duplex sequencing methods that computationally reconstruct dsDNA molecules after sequencing using unique molecular barcodes and require > 1 read per strand. This has low efficiency because Illumina sequencing is seeded by single-stranded DNA such that only a subset of sequencing reads can be reconstructed into dsDNA molecules[1-3]. At the same time, HiDEF-seq utilizes Pacific Biosciences sequencing that has a higher cost than Illumina sequencing.

For standard HiDEF-seq fragment sizes (1 to 4 kilobases; median 1.7 kilobase reads; Hpy166II digestion), given a cost of $1216 per SMRT cell (raw cost, excluding labor; prices current to December 31, 2022), and our observed $2.6 \cdot 10^9$ interrogated dsDNA base pairs per SMRT cell, HiDEF-seq's cost is $\$1,216 / 2.6 \cdot 10^9 = \$4.7 \cdot 10^{-7}$ per interrogated dsDNA base pair.

For large fragment HiDEF-seq (1 to 10 kilobases; median 4.2 kilobase reads; PvuII digestion), given a cost of $1216 per SMRT cell, and our observed $3.9 \cdot 10^9$ interrogated dsDNA base pairs per SMRT cell, HiDEF-seq's cost is $\$1,216 / 3.2 \cdot 10^9 = \$3.8 \cdot 10^{-7}$ per interrogated dsDNA base pair.

Given a cost of $14,465 per Novaseq S4 v1.5 300 cycle flow cell (raw cost, excluding labor; prices current to December 31, 2022), which sequences $3 \cdot 10^{12}$ bases, and assuming ideal post-filtering duplex efficiency of 0.04 for NanoSeq[1], NanoSeq's cost is $\$14,465/(0.04 \times 3 \cdot 10^{12}) = \$1.2 \cdot 10^{-7}$ per interrogated dsDNA base pair.

Therefore, for ultra-high fidelity dsDNA mosaic mutation detection, the cost of standard HiDEF-seq (1 to 4 kilboases) is 3.9-fold NanoSeq's cost, and the cost of large fragment HiDEF-seq (1 to 10 kilobases) is 3.2-fold NanoSeq's cost.

In 2023, a higher-throughput Pacific Biosciences Revio instrument will be released with a marketed sequencing cost of $995 per SMRT cell with 3.125 times higher throughput per SMRT cell relative to the SMRT cell used in this study. Therefore, the Revio instrument is projected to reduce the cost of standard HiDEF-seq to ($\$3.8 \cdot 10^{-7}$ per interrogated dsDNA base pair current cost) x ($995 / $1216) / 3.125 = $\$1.0 \cdot 10^{-7}$ per interrogated dsDNA base pair, which is less than the current cost of NanoSeq. In 2023, a higher-throughput Illumina Novaseq X instrument will also be released, with a marketed sequencing cost of \$2 per $10^9$ sequenced bases, which is projected to reduce the cost of NanoSeq to \$2 / (0.04 x $10^9$) = $\$5 \cdot 10^{-8}$ per interrogated dsDNA base pair. Therefore, after the release of both the Revio and Novaseq X instruments, the cost of standard HiDEF-seq will be 2-fold the cost of Nanoseq.

### Supplementary Note 3. Gating strategy for fluorescence-activated sorting of cerebral cortex neuronal nuclei

Flow cytometry data from a representative sample is shown in **Supplementary Fig. 2**, illustrating the gating strategy used to isolate nuclei of cerebral cortex neurons. See **Methods** for further details.

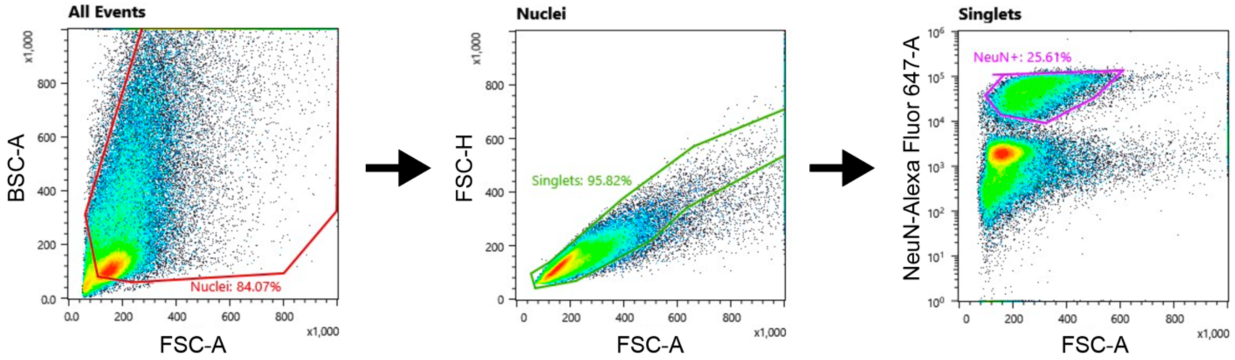


**Supplementary Figure 2.** Gating strategy for sorting cerebral cortex nuclei. Annotated percentages are calculated out of the total number of events.

### Supplementary Notes References


1 Abascal, F. *et al.* Somatic mutation landscapes at single-molecule resolution. *Nature* **593**, 405-410 (2021). https://doi.org:10.1038/s41586-021-03477-4

2 Hoang, M. L. *et al.* Genome-wide quantification of rare somatic mutations in normal human tissues using massively parallel sequencing. **113**, 9846-9851 (2016). https://doi.org:10.1073/pnas.1607794113 %J Proceedings of the National Academy of Sciences

3 Kennedy, S. R. *et al.* Detecting ultralow-frequency mutations by Duplex Sequencing. *Nature Protocols* **9**, 2586 (2014). https://doi.org:10.1038/nprot.2014.170
